# Genome-wide Quantification of Polycistronic Transcription in *Leishmania major*

**DOI:** 10.1101/2023.11.23.568479

**Authors:** Janne Grünebast, Stephan Lorenzen, Joachim Clos

**Author notes:** ***Correspondence to*** Joachim Clos, Leishmania Genetics Group, Bernhard Nocht Institute for Tropical Medicine, Bernhard Nocht St. 74, D20359 Hamburg; or: Janne Grünebast, Institute for Genome Sciences, School of Medicine, University of Maryland, Baltimore, MD 21201, USA.

## Abstract

*Leishmania major* is a human-pathogenic, obligate parasite and the etiological agent of the most prevalent, cutaneous form of leishmaniasis, which is an important neglected, tropical disease with ∼1.2 Mio new infections per year. *Leishmania*, and the whole order Trypanosomatida, are early eukaryotes with highly diverged gene expression and regulation pathways, setting them apart from their mammalian hosts and from most other eukaryotes. Using precision run-on sequence analysis, we performed a genome-wide mapping and density analysis of RNA polymerases in isolated nuclei of the protozoan parasite *Leishmania major*. We map transcription initiation sites within the chromosomes and correlate them with known sites of chromatin modifications. We confirm continuous, polycistronic RNA synthesis in all RNA polymerase II-dependent gene arrays but find varying RNA polymerase activities in polycistronic transcription units (PTUs), excluding gene-specific transcription regulation, but not PTU-specific variations as possible targets of modulatory pathways. Lastly, we find evidence for transcriptional pausing of all three RNA polymerase classes, hinting at a possible mechanism of transcriptional regulation.

**Significance Statement:** *Leishmania* spp. are pathogens of humans and animals and cause one of the most important neglected tropical diseases. Regulation of gene expression in *Leishmania* but also in the related *Trypanosoma* is radically different from all eukaryotic model organisms, dispensing with regulated, gene-specific transcription, and relying instead on highly regulated translation. Our work sheds light on the initiation, elongation and termination of transcription, maps unidirectional, polycistronic transcription units, provides evidence for transcriptional pausing at or near starting points of RNA synthesis, and quantifies the varying transcription rates of the polycistronic transcription units. Our results will further the understanding of these important pathogens and should provide a valuable ressource for researchers in the field of eukaryotic microbiology.

## Introduction

The eukaryotic genus *Leishmania* comprises a large number of parasitic protozoa which cause morbidity and mortality in humans and in a range of domestic and wild animals. The three main clinical forms of human *Leishmania* infections are cutaneous, mucocutaneous and visceral leishmaniasis. The latter form is lethal if left untreated. With 1.6 mio new infections per year and an estimated 20,000 fatalities (1), leishmaniasis is ranked second amongst the Neglected Tropical Diseases in terms of human suffering and mortality (2). There are no vaccines for human use, and the available chemotherapeutics are expensive and fraught with severe side effects (3).

The order Trypanosomatida, to which the leishmaniae belong, is an early branch of the eukaryotic tree, set apart from the crown group of eukaryota by their peculiar gene regulation mechanisms. All protein-coding genes are arranged in so-called polycistronic transcription units (PTUs), large unidirectional arrays of functionally non-related genes (4,5), separated by converging or diverging strand switch regions (cSSRs or dSSRs, respectively) which, according to the current view, are the places of transcription termination or initiation, respectively. Protein-coding genes of *Leishmania* have no distinct promoters, and the genomes harbour no genes for canonical transcription factors (6,7). However, recent studies in *Trypanosoma brucei* suggest the presence of sequence-specific motifs that act as core promoters in unidirectional PTUs (8,9). As the name implies, polycistronic transcription units are thought to be transcribed in a continuous manner. The resulting pre-mRNAs are processed by polyadenylation and trans-splicing into monocistronic mRNAs, which are translocated to the cytoplasm for translation (7). Each *Leishmania* mRNA therefore carries the same 39-nt spliced leader RNA sequence at the 5’ end which also bears a unique cap structure.

Polycistronic transcription was first investigated for loci and gene arrays by nuclear run-on transcription analysis using radioactive labelling of nascent RNA and hybridisation to immobilised gene sequences, followed by autoradiography (10–13). The results supported the concept of a polycistronic transcription and ruled out an inducible transcription of the investigated genes, mostly heat shock genes. The same methodology was also applied to extended gene arrays (14) and to small *Leishmania* chromosomes (15–17), also supporting the concept of a polycistronic transcription.

Epigenetic factors affecting transcription were analysed, too. Increased appearance of a modified thymine base, BaseJ, maps to presumed regions of transcription termination, i.e. cSSRs (18), while acetylated histone 3 (H3ac) is found predominantly in nucleosomes near transcription start regions, i.e. dSSRs (19). The combination of BaseJ followed by a region of H3ac also marks potential internal transcription termination and initiation sites within PTUs, so called head-tail regions (18,20,21). Moreover, chromatin density was found to fluctuate at dSSRs and 5’-telomeric sites, regions of transcription initiation, during in vitro stage conversion of *L. donovani* (22).

We therefore decided to investigate RNA synthesis in *Leishmania* on a genome-wide scale, applying Precision Run-On sequencing (PRO-seq) (23). By performing a nuclear run-on reaction in the presence of biotinylated CTP, we at once labelled and terminated nascent RNAs in isolated nuclei, thereby mapping the positions of RNA polymerase elongation complexes. We show that transcription is truly polycistronic and unidirectional in all PTUs in the *L. major* genome, but find that PTUs are transcribed at different rates suggesting either an underlying mechanism of transcriptional control or varying efficacies of transcription initiation sites. We verify dSSRs and cSSRs as sites of transcription initiation and termination, respectively, establish a correlation between chromatin modifications and transcription initiation/-termination, and therefore describe that BaseJ and H3ac not always lead to a termination and reinitiation of transcription within PTUs. Furthermore, we detect a significantly reduced nuclear run-on transcription in reactions with sarkosyl within transcription initiation sites in dSSRs and 5’-telomeric transcription start sites, but also in RNA polymerase I- and III-dependent genes, suggesting pausing of RNA polymerases as a new mechanism of RNA synthesis modulation.

## Materials and Methods

### Cell culture of L. major

*Leishmania major* strain Friedlin was cultured at 25°C in M199+ medium (1x M199, 20% FCS (heat-denatured), 40 mM HEPES, 10 mg/L haemin, 0.1 mM adenine, 5 μM 6-biopterin, 2 mM L-glutamine, 100 U penicillin, 100 μg/mL streptomycin). The parasites were diluted every 3-4 days to a cell count of 1×10^5^.

### Nuclei isolation from L. major

From a logarithmically growing culture, 2 ✕ 10^9^ cells were taken and sedimented at 4°C. The pellet was then washed with ice-cold 1× PBS (4°C, 15 min, 1000 ✕ g), and the cells were lysed in 5 mL ice-cold lysis buffer (10 mM Tris-HCl pH 8.0, 10 mM sodium chloride, 5 mM DTT, 1.5 mM magnesium chloride, 1 mM spermidine, 1 mM EDTA, 0.1 mM PMSF) containing 0.5% (vol/vol) Tergitol 15-S-9 for 5 min on ice. After centrifugation (4°C, 15 min, 1000 ✕ g), the cytosolic supernatant was carefully decanted, and the nuclei were washed in 5 mL of ice-cold lysis buffer without Tergitol. The nuclei were then resuspended in 200 μL of lysis buffer without Tergitol, and an equal volume of 2x nuclei storage buffer (80% (vol/vol) glycerin, 10 mM Tris-HCl pH 8.3, 0.2 mM EDTA) was added.

### Nuclear run-on and RNA isolation

The nuclear run on protocol follows a protocol by Mahat et al., 2016 (23) with slight adaptions to *Leishmania* gene expression mechanisms. Individual run-ons with biotinylated CTP were performed. First, the 2x run-on mix (2x NRO) was prepared as follows: 40µl 5x nuclear run-on buffer (80 mM Tris-HCL pH 7.5, 50 mM sodium chloride, 50 mM potassium chloride, 8 mM magnesium chloride, 0.5 mM spermidine), 2.5 µl ATP (10 mM), 2.5 µl GTP (10 mM), 2.5 µl UTP (10 mM), 1 µl CTP (0.05 mM), 5 µl biotin-11-CTP (1 mM) (Jena Bioscience), 0.8 µl DTT (0.5 M) 40 U SUPERase-InTM RNase Inhibitor (Ambion®) and 43.7 µl 2% Sarkosyl for run-ons with sarkosyl, or nuclease-free water for run-ons without sarkosyl. For the nuclear run-on reaction, 100 μL of nuclei suspension were added to 100 μL of prewarmed 2x NRO using a wide bore pipette tip. The reaction was incubated for 10 min at 37°C and stopped by the addition of 500 μl Trizol LS, followed by a vigorous mixing. 130 µl chloroform was added, the sample was shaken and then centrifuged at 14,000 ✕ g for 5 min at 4°C. The aqueous phase was transferred to a new tube and mixed with 2.5 vol 100% (vol/vol) ethanol. After incubation for 10 min at room temperature, the RNA was sedimented at 14,000 ✕ g for 20 min at 4°C. The supernatant was completely removed, and the pellet was washed once with 75% (vol/vol) ethanol. The pellet was dried and dissolved in 20 ul of nuclease free water.

### PRO-seq library preparation

The library was prepared as in Mahat et al., 2016 (23), but without hydroxyl repair of the 5’ end due to the different cap structure in trypanosomatidic spliced leader RNA. Briefly, RNA fragmentation was performed by base hydrolysis with 5µl 1 N NaOH. The sample was neutralized by 25 μL 1M Tris-HCl pH 6.8, and a buffer exchange was performed using a BioSpin P-30 column (Bio-Rad) according to the manufacturer’s instructions.

Biotin-labeled RNAs were enriched using streptavidin M280 beads. For each library, 90 µl beads were washed in preparation buffer (0,1 N NaOH, 50 mM NaCl) and then twice in 100 nM NaCl. The beads were then resuspended in 50 µl binding buffer (10 mM Tris-HCl pH 7.4, 300 mM NaCl, 0.1% (vol/vol) Tergitol 15-S-9) per reaction and incubated with 50 µl of nuclear run-on RNA for 20 min at room temperature on a rotating wheel (8 rpm). Beads were washed twice with high salt buffer (50 mM Tris-HCl pH 7.4, 2 M NaCl, 1 mM EDTA, 0.5% (vol/vol) Tergitol 15-S-9) and once with low salt buffer (5 mM Tris-HCl pH 7.4, 1 mM EDTA, 0.1% (vol/vol) Tergitol 15-S-9). Beads were resuspended in 300 µl Trizol, and after vortexing and a 3 min incubation, 60 µl chloroform was added. This was followed by a centrifugation step at 14,000 ✕ g for 5 min at 4 °C. The upper, aqueous phase was transferred to a new tube and the organic phase discarded. The same beads were extracted a second time using Trizol and chloroform as previously described. The aqueous phases were combined and extracted with 1 Vol of chloroform followed by centrifugation (14,000 ✕ g, 5 min, 4°C). To the aqueous phase, 1 μL GlycoBlue and 900 μL 100% ethanol were added and thoroughly mixed. After incubation (10 min, RT), the RNA was precipitated (14,000 ✕ g, 20 min, 4°C). The pellet was washed in 75% (vol/vol) ethanol (14,000 ✕ g, 5 min, 4°C) and dried for 5-10 min.

For 3’ adapter ligation, the pellet was resuspended in 12.5 µM VRA3 Adapter (/5Phos/ GAUCGUCGGACUGUAGAACUCUGAAC /inverted dT/), incubated at 65°C for 20 sec and placed immediately on ice. A ligation mix was added and the mixture was incubated for 4 h at 20°C. Subsequently, a second biotin enrichment was performed (see above). This was followed by a 5’ adapter ligation with VRA5 (CCUUGGCACCCGAGAAUUCCA) and a third biotin enrichment. The adapter-ligated RNA was reversely transcribed and a test PCR amplification for optimizing the cycle number was performed. After final amplification, the library was purified from an 8% polyacrylamide gel to remove adapter dimers. Concentrations were determined using the KAPA Library Quantification Kit (Roche) and librarys were sequenced on a NextSeqTM 550 system (Illumina) using a NextSeq 500/550 High Output Kit v2.5 (150 Cycles) single-end according to the manufacturer’s instructions.

The quality of the libraries was checked by the length distribution of the reads. Approximately 20 bases are protected from degradation by RNA polymerase between the RNA polymerase exit channel and the 3’ RNA end, so libraries below 20 nucleotides indicate degradation of the RNA after the nuclear run-on. A fragment size of 20-30 nucleotides is still considered as partial degradation (24). No RNA degradation could be detected in our libraries (Figure S1).

#### Bioinformatical analysis

PRO-seq reads were trimmed using cutadapt (25) and aligned to *L. major* Friedlin2021 (26) using Bowtie2 (27). Positions were extracted using Bedtools (28). Counts were normalised by aligned nucleotides per sample. Positions for acetylated Histone 3 were determined by aligning the sequences in the microarray to the Friedlin2021 genome using Bowtie2 (29) with parameter -k 5. For this alignment, only replicate 2 of (19) was used. BaseJ reads from (18) were aligned to the Friedlin genome using Bowtie (29) with parameter -k 5 and read coverage was calculated using in house scripts. BaseJ and H3ac alignments were smoothed using a running average with a window of 1000 bases. PTUs were defined as stretches from one ATG to the last stop codon of consecutive genes in the same direction. dSSRs and cSSRs were defined as the regions between two divergent or convergent genes, respectively. 5’-telomeres are the regions 5’ of a terminal gene to the end of the chromosome and 3’-telomeres are regions from 3’ end of a gene to the end of the chromosome. Positions with Ns in the reference sequence and positions covered by RNA Pol I or RNA Pol III transcribed genes were not counted for PTUs, SSRs and telomeric regions. Normalised counts of three replicates were summed up and averaged. Read counts in Figure 2, 3, 5 and 6K-M and Supplementary Figure 2-6 are plotted logarithmically with pseudocounts.

## Results

The PRO-seq nuclear run-on transcription protocol as described in (23) allows a genome-wide mapping of RNA polymerase elongation complexes in the intact chromatin of isolated nuclei, reflecting positions and densities of elongation complexes at the time of cell fractionation.

Simplified, *L. major* promastigote cells were ruptured according to our established protocol (13), replacing the banned Triton X-100 with Tergitol. Aliquots of cryogenically stored nuclei were then added to a nuclear run-on mix, with or without the detergent sarkosyl, and containing ATP, GTP, UTP, and biotinylated CTP (Figure 1A). This allowed RNA elongation to continue until the next GTP moiety in the noncoding strand DNA was encountered, at once labelling the nascent RNA with a 3’-terminal biotin tag and blocking further elongation. The nuclear run-on reaction was performed in the presence or absence of sarkosyl, an anionic detergent, that causes restarting of paused transcription complexes but not of terminated complexes (30,31). Furthermore, only transcription-competent complexes can be detected with PRO-seq as the run-on reaction is dependent on the incorporation of nucleotides (31).

**Figure 1:**
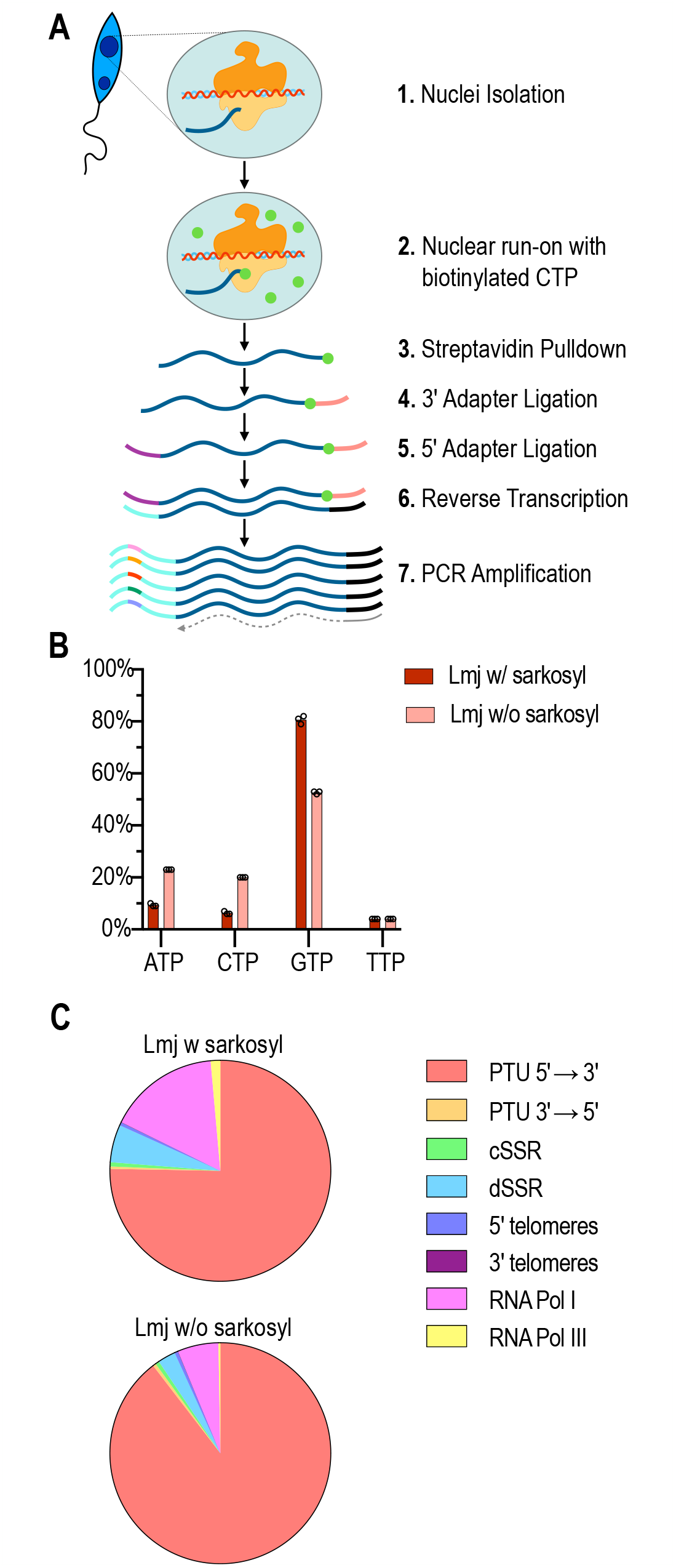
**A** Overview of PRO-seq: Nuclei are isolated and an in vitro nuclear run on reaction with biotinylated CTP is performed. After subsequent RNA isolation, the native RNA is purified with streptavidin beads. This is followed by 3’ and 5’ adapter ligation and reverse transcription. The libraries are then individually indexed by barcode PCR and sequenced with Illumina. **B** Percentage of bases at position 1 of reads in *L. major* with sarkosyl (Lmj w/ sarkosyl, dark red) and without sarkosyl (Lmj w/o sarkosyl, light red), n=3. **C** Overall distribution of reads of *L. major* with and without sarkosyl in PTUs, SSRs, telomeres and in RNA Pol I and III genes, n=3.

The nuclear run-on transcripts were then enriched by a streptavidin pull-down and further processed for Illumina sequencing by 3’- and 5’-adapter ligation, reverse transcription and PCR amplification (Figure 1 A). The resulting libraries were subjected to NextSeq 550 (High Output, 150 cycles) sequencing.

We performed PRO-seq with *L. major* logarithmic phase promastigotes in biological triplicates with or without sarkosyl. All libraries yielded comparable raw read numbers; however, 66% of reads derived from sarkosyl-treated nuclei remained after trimming, compared with 40% of reads from untreated nuclei (see Table S1). Moreover, when we analysed the 5’ dNTPs of the sequencing reads from the sarkosyl libraries (Figure 1 B, Table S1), we found an 80% majority starting with a GTP, the complementary base of the biotin-CTP used to stop elongation, while only 53% of the reads from untreated nuclei started with GTP. This indicates that RNA polymerases are blocked more efficiently by biotin-CTP under sarkosyl, given that libraries originated from a strepavidin-selected nascent RNAs.

We next aligned the reads to the *L. major* Friedlin genome (26) and performed a global RNA polymerase density analysis for PTUs in sense (5’→3’) and antisense (3’→5’) direction (according to the TriTrypDB annotation), for cSSRs and dSSRs, 5’- and 3’-telomeric regions, RNA polymerase I and RNA polymerase III genes (Figure 1 C). We find that under sarkosyl, i.e. mapping of all elongation complexes, including paused RNA polymerases, the proportions of elongating RNA polymerase I and III complexes at their dependent genes increases by over 2-fold compared with minus-sarkosyl reads, i.e. active elongation complexes only. A similar effect is seen for reads aligning to diverging SSRs. We also observe an overwhelming preference of RNA polymerases for the sense strand in the PTUs, confirming the concept of unidirectional, strand-specific transcription in the PTUs on a genome-wide level.

The mapped positions of the reads are shown in the form of a whole genome Circos plot for all 36 chromosomes of *L. major* (Figure 2) and in linear form by chromosomes (Figure S2). From the outside to the inside, the Circos plot shows the chromosomes, the open reading frames according to the TriTrypDB, the PRO-seq read density w/ sarkosyl, the PRO-seq read density w/o sarkosyl, the relative prevalence of H3ac, taken from (19), and the relative prevalence of BaseJ nucleotides, taken from (18). The latter were added to analyse the impact of both chromatin modifications on transcription initiation and termination. Again, the highly selective, unidirectional transcription of PTUs is confirmed.

**Figure 2:**
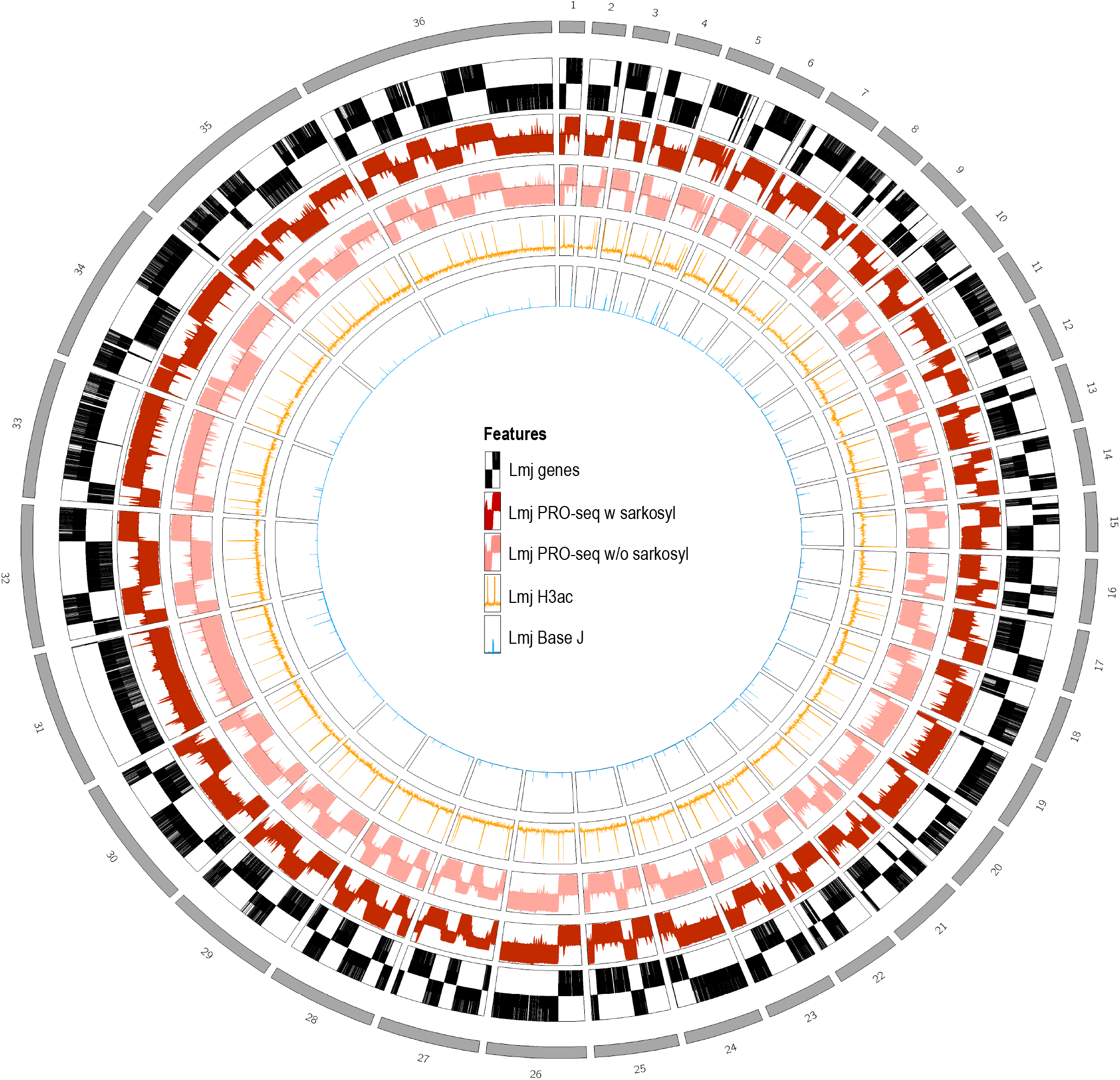
Circos plot. All 36 chromosomes of the *L. major* Friedlin 2021 genome are plotted in the outer ring (gray). Then genes follow, with UTRs plotted only halfway up and CDSs plotted over the entire length (black). Next plotted is a logarithmic representation of the PRO-seq data of *L. major* with sarkosyl (dark red) and without sarkosyl (light red), n=3. ChIP-chip data from acetylated histone H3 (H3ac) (19) follow in orange, and ChIP-seq data from glucosylated hydroxymethyluracil (BaseJ) (18) follow in blue. H3ac and BaseJ were aligned to the *L. major* Friedlin 2021 genome.

For a better understanding of transcription on a genome-wide scale, we displayed the PRO-seq reads in a higher resolution (Figures 3 and S2), allowing us to analyse interrupted transcription in head-tail regions marked by BaseJ (18) and H3ac (19).

**Figure 3:**
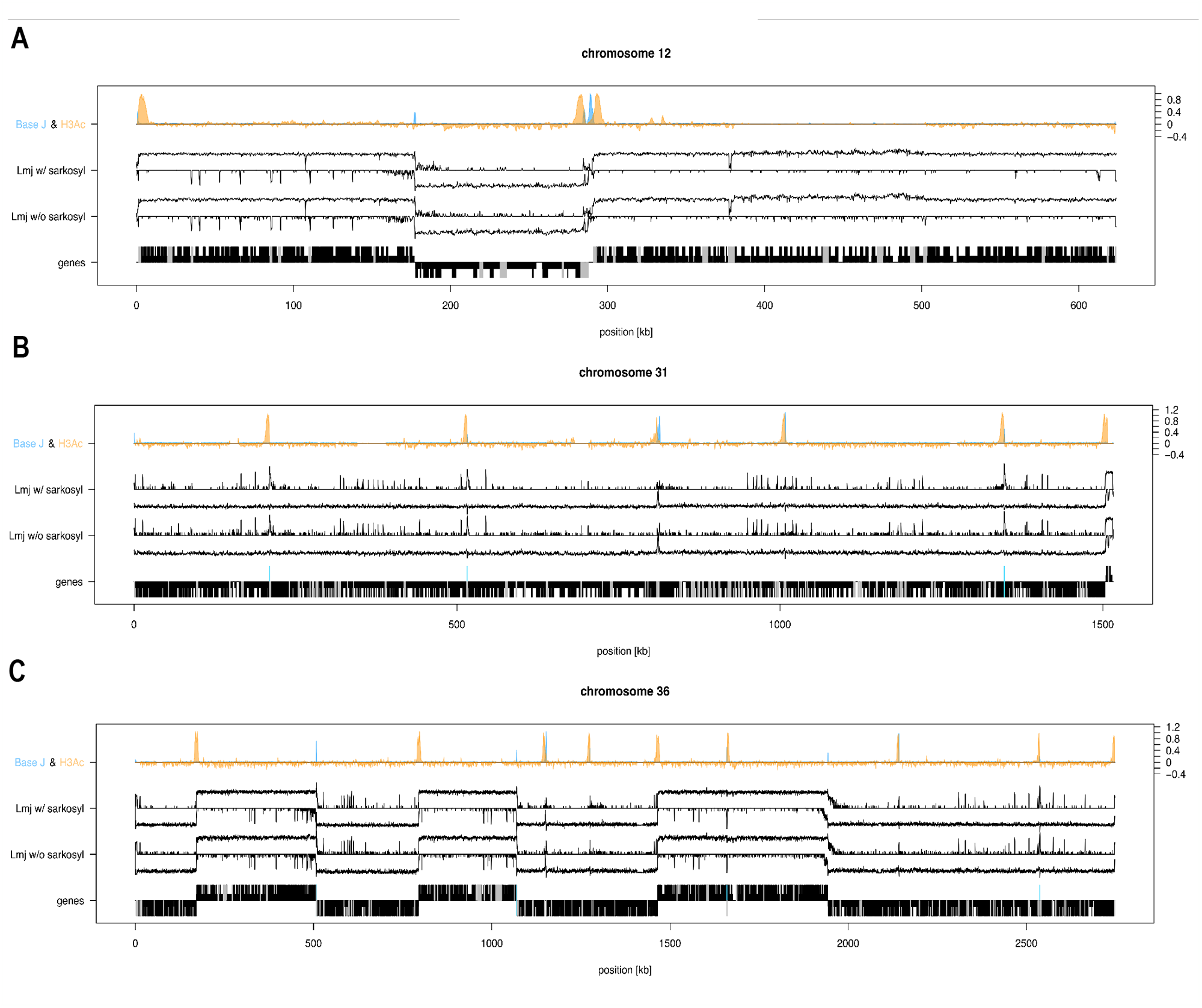
Chromosome-wide representation of PRO-seq data of *L. major* with sarkosyl (Lmj w/ sarkosyl) and without sarkosyl (Lmj w/o sarkosyl) on selected examples. PRO-seq data was plotted strand specifically and displayed logarithmically, n=3. ChIP-chip data from acetylated histone H3 (19) is plotted in orange, and ChIP-seq data from base J (18) is plotted in blue, each normalised within chromosomes. The bottom row shows genes (black), with UTRs plotted half-length and CDS plotted full-length. Noncoding RNAs are plotted in gray and RNA Pol III genes in turquoise. Examples shown are chromosome 12 (A), chromosome 31 (B), and chromosome 36 (C).

In chromosome 12 (Figure 3 A), transcription initiation in dSSRs matches H3ac peaks at a 5’ telomere and, in a double peak, at a divergent SSR. Between the twin H3ac peaks, two peaks of BaseJ abundance can be observed, likely in conjunction with the RNA Pol III transcribed genes in that dSSR (turquoise spikes), which is reflected in a spike of PRO-seq reads on that strand. Chromosome 31 consists of only two PTUs, with the 5’ PTU covering >95% of the length (Figure 3 B). Consequently, transcription proceeds unidirectionally. However, at position ∼800k, a BaseJ peak is followed by a H3ac peak, and we observe a concomitant drop of RNA polymerase density in our PRO-seq data, likely due to a termination followed by re-initiation of transcription. There are four more spikes of BaseJ and H3ac abundance of which three correspond to spikes of transcription of RNA Pol III transcribed genes on the opposite strand. However, RNA Polymerase density on the sense strand stays at the same level, meaning that no termination and re-initiation happens in the PTU despite the occurance of BaseJ and H3ac. Similar correlations can be seen on chromosome 36 (Figure 3 C). Again, BaseJ and H3ac spikes are sometimes but not always associated with localised interruptions of unidirectional transcription and corresponding spikes of polymerase activity on the opposite strand. We conclude that indeed, chromatin modifications correlate with transcription initiation and termination and that previously mapped PTUs may contain short regions of termination and reinitiation, perhaps with intermediate antisense transcription. However, the presence of BaseJ and H3ac not always results in termination and reinitiation of sense transcription within a PTU. Overall, the sense reads outnumber the antisense reads by over two orders of magnitude, both with and without sarkosyl (Figure 4 A, B). The PTUs show a different level of transcription with an average sense read count between 5 and 27 for samples with sarkosyl and 3 and 33 for samples without sarkosyl (Figure 4 A, Table S3). A closer look at single PTUs reveals varying RNA synthesis rates on one and the same chromosome (Figure 4 D, E). We displayed the PTUs by chromosome, as *Leishmania* often shows chromosomal aneuploidies, e.g. chromosome 31 being tetraploid. But even on chromosome 31, the PTUs are transcribed at different levels. In general, it can be observed that some chromosomes transcribe their PTUs at similar levels, e.g. chromosome 2, 4, 17, 18, 23, 26, 29 and 30, while some chromosomes show a large difference in read density at the PTUs, e.g. 3, 12 and 24. Chromosomes 35 and 36 show a different pattern, where the majority of PTUs are transcribed at the same level, with one PTU having a higher read density. The different read densities at the PTUs indicate either different efficiencies of the transcription initiation site involved or an additional mechanism regulating transcription.

**Figure 4:**
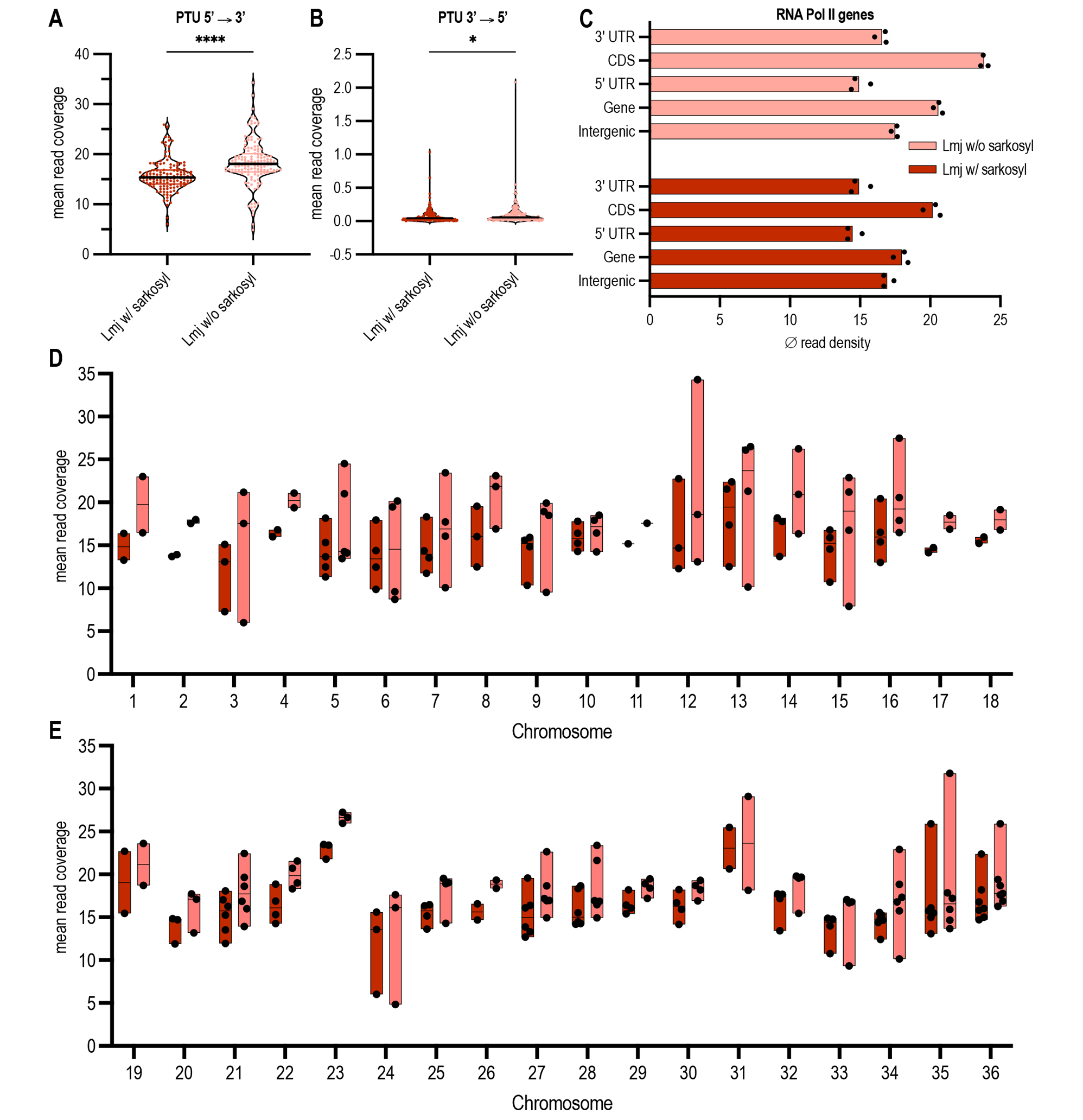
Read occupancy at PTUs. **A-B** Average occupancy of normalised PRO-seq reads within PTUs (n=132) in sense (5’→3’) **(A)** and antisense (3’→5’) **(B)** direction. Each dot represents the mean read coverage over one PTU of three biological replicates of either *L. major* with sarkosyl (Lmj w/ sarkosyl, dark red) or without sarkosyl (Lmj w/o sarkosyl, light red). Significance was tested using a paired students t-test; ****, P ≤ 0.0001; *, P ≤ 0.05. **C** Mean read coverage of normalised PRO-seq reads in RNA Pol II transcribed PTUs split by genes (UTR + CDS), 5’ UTRs, CDS, 3’ UTRs and intergenic regions within PTUs. The regions were summed and the mean over the entire regions in three biological replicates of *L. major* with sarkosyl (dark red) and without sarkosyl (light red) is shown. Each dot represents one biological replicate. **D-E** Mean read coverage of PRO-seq reads in PTUs across all 36 chromosomes of *L. major*. Each dot represents the mean read coverage of one PTU on that chromosome, n=3. Dark red boxes display samples with sarkosyl and light red boxes without sarkosyl. Chromosome 1-18 are shown in **(D)** and chromosome 19-36 in **(E)**.

We also compared PRO-seq read alignment between the 5’ and 3’ untranslated regions (UTRs) and the protein-coding sequences (CDS), but also between mRNA coding regions (genes) and intergenic regions. The results (Figure 4 C) show a ∼25% drop in alignment frequency between protein-coding sequences and the untranslated regions (UTRs), but confirm the visual impression that transcription proceeds through intergenic regions with little if any drop of polymerase density.

Next, we took a closer look at the strand switch regions to localise transcription initiation and termination regions. In Figure 5 A, we present RNA polymerase densities in 3 prototypic of 58 diverging SSRs. While in dSSR6, a central part is virtually devoid of RNA polymerase complexes, occupancy of the two DNA strands overlaps significantly in dSSR22, while in dSSR46, transcription initiates head-to-head. For RNA polymerase occupancy maps of all dSSRs see Figure S3, information about the positions can be found in Table S2.

**Figure 5:**
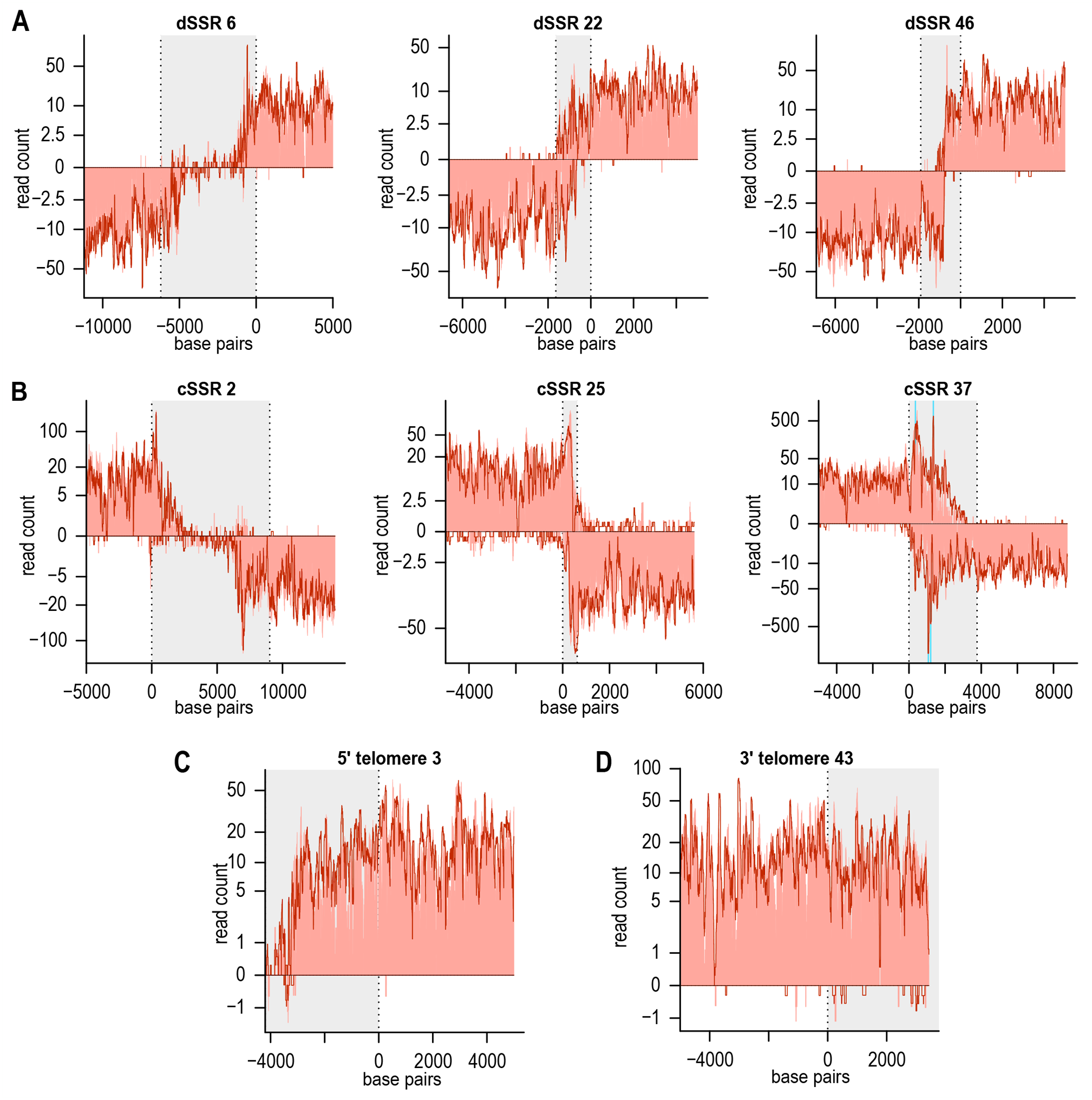
Distribution of normalised PRO-seq reads of *L. major* with and without sarkosyl at transcription initiation and termination sites. **A** Transcription initiation in divergent strand switch regions (dSSR 6, 22 and 46). dSSRs are highlighted in grey with the dashed lines representing the start of the PTUs. Normalised PRO-seq reads of *L. major* with sarkosyl are plotted logarithmically as dark red line and reads of *L. major* without sarkosyl as light red background. The first 5000 bp of each PTU are shown, n=3. **B** Transcription termination within convergent SSRs (cSSR 2, 25 and 37). cSSRs are shown in grey and the dashed lines represent the stop of the PTU with the last 5000 bp of the PTU shown. Turquoise background shows RNA Pol III genes (cSSR 37), n=3. **C-D** Transcription initiation in 5’ telomeres **(C)** and transcription termination in 3’ telomeres **(D)** are shown. The grey background represents the telomeric region with the dashed line indicating the start or stop of the PTUs. The first 5000 bp of the PTUs are plotted, n=3.

Likewise, the occupancy patterns of cSSRs are also diverse. cSSR2 shows a clear gap of polymerase occupancy (Figure 5 B), while PTUs in SSR25 are tail-to-tail. An occupancy overlap on both DNA strands can be seen in cSSR37. The occupancy patterns for cSSRs 1 to 38 can be viewed in Figure S4.

We also analysed fifteen 5’ telomeric transcription start regions (Figure S5) and 56 3’ telomeric transcription termination regions (Figure S6). The example of 5’ telomere 3 (Figure 5 C) which shows transcription starting very close to the 5’ end of the chromosome is representative for the rest (Figure S5). Likewise, transcription proceeds to the very 3’ ends of the chromosomes in 3’ telomere 43 (Figure 5 D), again reflecting a general pattern (Figure S6).

Lastly, we looked at RNA polymerase pausing based on the sarkosyl-induced restarting (Figure 6). When analysing PTUs (Figure 4 A, D, E, Table S3), we generally observe a higher read occupancy in samples without sarkosyl, i.e. active transcription elongation complexes only, compared to run-on reactions with sarkosyl, i.e. paused and active transcription elongation complexes, suggesting that sarkosyl is not in general boosting transcription but rather acting specifically in *Leishmania* as previously described in other eukaryotes (30,32). Looking at dSSRs in general, RNA polymerase II complexes show a moderate but significant occupancy increase of reads in nuclear run-on reactions with sarkosyl (Figure 6 A). At higher resolution, RNA synthesis in the absence of sarkosyl is markedly lower at the start of the PTUs (Figure 6 D), suggesting paused RNA polymerase complexes at transcription initiation sites. The same is observed at the border region and the beginning of PTUs downstream of 5’-telomeric start sites (Figure 6 F), but no significant reduction was noticed within the 5’-telomeres (Figure 6 C). However, there is no distinct peak of paused RNA polymerase complexes in dSSRs and 5’ telomeres; rather, longer stretches are affected. In regions of transcription termination i.e. cSSRs, and 3’ telomeres, no RNA polymerase II pausing is observed (Figure 6 B, E, G).

**Figure 6:**
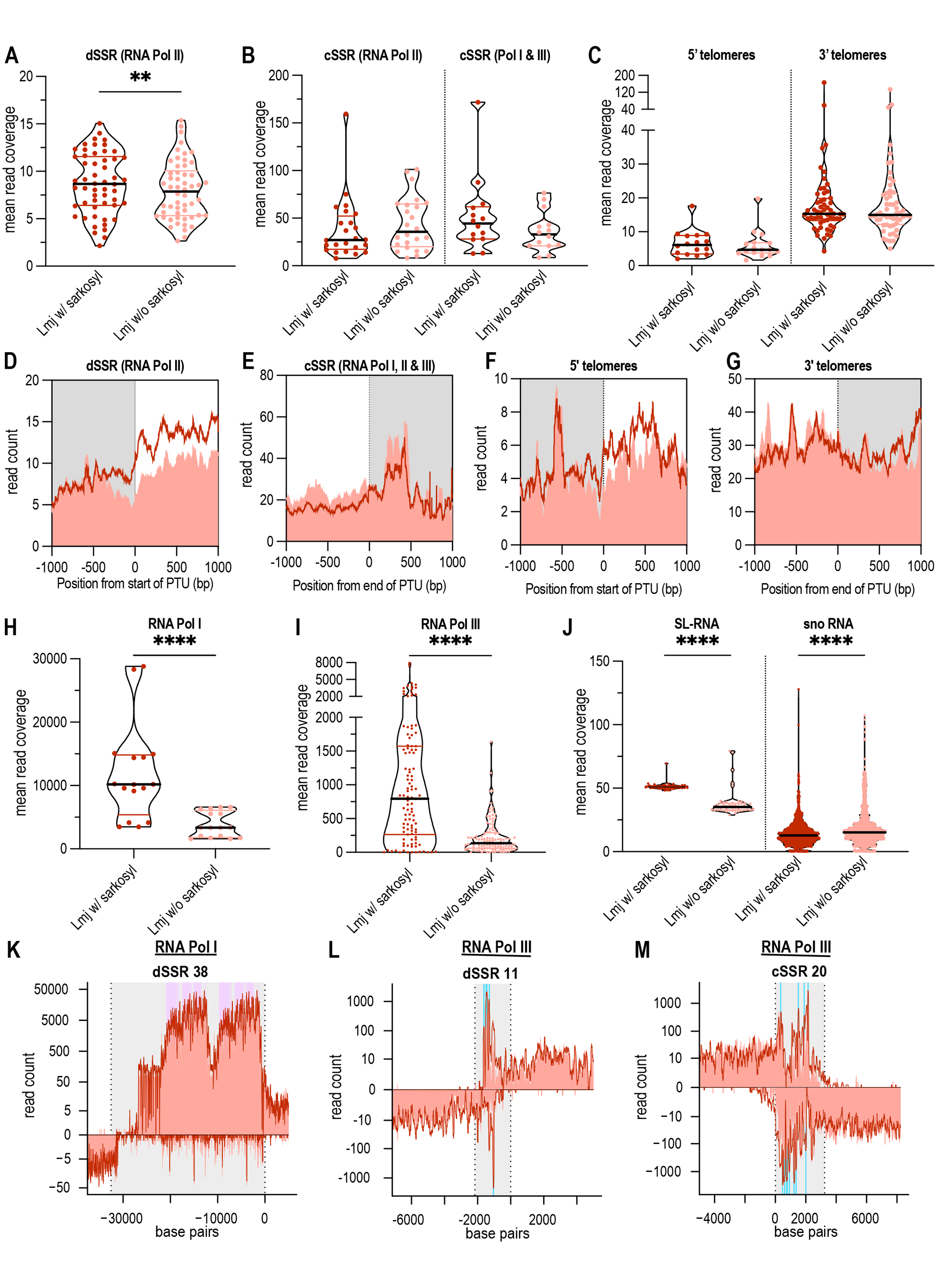
**A-C** Occupancy of dSSRs **(A)**, cSSRs **(B)**, 5’ telomeres and 3’ telomeres **(C)** with normalised reads from *L. major* with sarkosyl (dark red) and without sarkosyl (light red). Each dot shows the mean read coverage in a dSSRs, cSSRs, 5’ telomeres, or 3’ telomeres, n=3. dSSRs were only counted if there was no RNA Pol I or III gene locus within the dSSR (RNA Pol II, 55 out of 58 dSSRs). cSSRs were differentiated into cSSRs in which RNA Pol I and III gene loci are located (RNA Pol I & III, n=14) and in which no additional gene loci is located and only RNA Pol II transcription stops here (RNA Pol II, n=24). No additional RNA Pol I & III gene loci are located in 5’ telomeres (n=16) and 3’ telomeres (n=56). Significance was tested using a paired students t-test; **, P ≤ 0.01; ns, P > 0.05. **D-G** Mean read coverages of normalised PRO-seq reads at the start of a PTU in dSSRs **(D)** and 5’telomeres **(F)** and at the end of a PTU in cSSRs **(E)** and 3’telomeres **(G)**. The first and the last 1000 bp around the start/end of a PTU are shown. *L. major* with sarkosyl are plotted as dark red line and reads of *L. major* without sarkosyl as light red background, n=3. **H-J** Distribution of normalised PRO-seq reads within RNA Pol I and RNA Pol III transcribed genes and within non coding RNAs. Occupancy of PRO-seq reads in RNA Pol I (n=16) **(H)** and RNA Pol III (n=107) **(I)** regions. Each dot shows the mean read coverage of three biological replicates from *L. major* with sarcosyl (dark red) and without sarcosyl (light red) in a RNA Pol I or III transcribed region. Significance was tested using a paired students t-test; ****, P ≤ 0.0001. **(J)** Occupancy of the spliced leader RNAs (SL-RNA, n=53) and small nucleolar RNAs (sno RNA, n=1128) with PRO-seq reads. Significance was tested using a paired students t-test; ****, P ≤ 0.0001; n=3. **K** RNA Pol I gene locus of 5.8S, 18S, 28S rRNA within dSSR 38. The SSR is shown in grey and the dashed lines represent the start of the PTU with the first 5000 bp of the PTU shown. Purple background shows RNA Pol I genes. Normalised PRO-seq reads of *L. major* with sarkosyl are plotted logarithmically as dark red line and reads of *L. major* without sarkosyl as light red background, n=3. **L-M** RNA Pol III genes are shown within one dSSR and one cSSR. Plot is the same as in **K**, but for the cSSR the last 5000 bp of the PTU are shown. Turquoise background shows RNA Pol III genes. Three biological replicates of either *L. major* with or without sarkosyl are shown.

Paused RNA polymerases can be seen in genes transcribed by RNA polymerase I, i.e. large ribosomal RNA genes (28S-, 18S- and 5.8S-rRNA; Figure 6 H, K), or RNA polymerase III, (tRNAs, 5S-rRNA; Figure 6 I, L, M). Spliced leader RNA (SL-RNA) genes, transcribed by RNA polymerase II, are also occupied by a sizable percentage of paused transcription complexes, while the functionally related small nucleolar RNA genes, are not sites of paused RNA polymerases (Figure 6 J). The products of these genes, ribosomal RNAs and tRNAs, spliced leader RNAs and snoRNAs are all known to be crucial for protein synthesis and cell proliferation.

## Discussion

### PRO-seq to analyse active transcription in Leishmania

In this study, we provide a genome-wide view of transcription in *L. major* using PRO-seq analysis. Our aim was to monitor ongoing active transcription by mapping RNA polymerase elongation complexes within PTUs, telomeres and strand switch regions and to identify the transcription initiation and termination sites on all 36 chromosomes. The information derived from PRO-seq pertains to an earlier stage of gene expression than the commonly applied RNA-seq analyses, which analyses the mRNA steady state levels which are the result of transcription followed by RNA splicing minus RNA degradation (6) and are therefore not a measure of transcription.

To gain insight into active transcription there are a number of sequencing methods available. PRO-seq is a further development of GRO-seq (Global Run-On sequencing) (33), in which bromouridine-labelled nascent RNAs are purified via affinity chromatography and then sequenced. However, GRO-seq does not offer a nucleotide-precise resolution like PRO-seq, but only an accuracy of about ±20 nt. Another alternative to PRO-seq is NET-seq (Native Elongating Transcript sequencing), in which the nascent RNAs are purified and sequenced via immunoprecipitation of the RNA Pol II elongation complex (34). Nevertheless, immunoprecipitation can also lead to precipitation of other complexes due to non-specific antibody binding and, in addition, the nascent RNAs must be at least 18 nt in length. Since we are interested in transcription by all RNA polymerase classes, we opted for PRO-seq. To our knowledge, this study presents the first genome-wide PRO-seq analysis of transcription in Trypanosomatida.

PRO-seq was first used to analyse transcription initiation and pausing around the *Drosophila melanogaster* Hsp70 promoter (35) and had to be adapted for use in *Leishmania*. In *Leishmania*, capping of mRNA occurs during the process of trans-splicing (6), therefore no cap is present during elongation of nascent RNA, making the decapping step in the original protocol unnecessary (Figure 1 A). The nuclear run-on reaction was performed with a single biotin-labelled NTP, as the main purpose was to map and quantify RNA polymerase elongation complexes for which single-nucleotide accuracy was not needed. We picked biotinylated CTP due to the GC richness (>60%) of the *Leishmania* genome. Since the insertion of a biotinylated NTP causes termination of transcription, the start of each read should be a guanine base, as sequencing was performed from the 3’ end making the reads reverse complementary. Kwak et al., 2013 were able to show in their experiment that the substrate-specific base was incorporated by 67 to 88% in the presence of sarkosyl. In our analysis we achieved 80% and 53 % guanine residues in read position 1 performing the run-on with or without sarkosyl, respectively (Figure 1 B). We conclude that sarkosyl not only leads to the restarting of transcription by paused RNA polymerases, but also prevents read-through transcription over incorporated biotin-CTPs.

### Polycistronic transcription units are transcribed at varying levels

The first supporting evidence for polycistronic transcription in trypanosomes came from northern blot analyses when RNA species encoded by intergenic regions of two different gene families were detected (36–38). Nuclear run-on analyses of gene loci (11,12,39,40), a part of a chromosome (14) or full chromosomes 1 and 3 (15–17) also showed transcription in *Leishmania* to be polycistronic in the investigated loci and chromosomes. Wedel et al. provided in 2017 the first sequencing-based proof of polycistronic transcription in *Trypanosoma brucei* by applying 5′-triphosphate-containing small RNA sequencing reflecting primary transcripts (8). Our PRO-seq data provide final proof for polycistronic transcription in *Leishmania* on a genome-wide scale (Figure 2, 3). PRO-seq reads are distributed within PTUs in a unidirectional, strand-specific manner on all 36 chromosomes of *L. major*. Coverage of reads in sense direction is two orders of magnitude higher compared with antisense direction (Figure 4 A, B). The first nuclear run-on analysis in *Leishmania,* which focussed on the beta-tubulin gene cluster, already showed a strand-specific transcription (41). Later, strand-specific transcription was confirmed in other nuclear run-on analyses (15,17) and through strand-specific RNA-seq (20).

Since there are no defined gene-specific promoters for the genes transcribed by RNA Pol II in trypanosomes (with the exception of the SL gene locus), it was assumed for a long time that regulation takes place exclusively at the post-transcriptional level and that transcription is not regulated at the level of transcription initiation. Transcription elongation rates of PTUs remained unknown as sequencing-based methods mostly analyse mRNA abundance levels (6,42). A newer study from 2022, however, used ChIP-seq with an antibody binding to RNA Pol II of *T. brucei* and identified potential promotor regions of polycistronic transcription initiation (9). It was also shown that sequence-specific promoters lead to different luciferase reporter activity in transient transfection assays, which was also found in a study by Wedel et. al. analysing GT-rich promoter sequences in reporter assays (8). Our PRO-seq analysis revealed genome-wide differences in the occupancy of PTUs with transcriptional elongation complexes, which is evidence that PTUs indeed have different transcriptional activities (Figure 4 D, E, Table S3). Due to the aneuploidy for which Leishmania is known (43), we specifically analysed PTUs located on the same chromosomes and found large differences in transcription levels for some chromosomes, suggesting an underlying mechanism of transcriptional modulation of individual PTUs in *Leishmania*.

Moreover, we observe a ∼25% drop of RNA polymerase density in the 5’ and 3’ untranslated regions (Figure 4 C). We have no ready explanation for this finding, but such variations between coding sequences and flanking regions were observed before for the HSP70 gene repeat (11).

### Transcription initiation and termination

Due to the lack of defined promoters and canonical transcription factors in trypanosomes (44) transcription initiation and termination remain poorly understood. As mentioned above, only two studies have identified sequence-specific promoters in *T. brucei*, one being a long GT-rich sequence (8) and the other a 75-bp promoter element (9). Here, we provide a genome-wide view of transcription initiation and termination sites in *L. major*. In prototypic Eukaryotes, initiation and termination of transcription is regulated by trans-acting protein factors, which *Leishmania* lacks, but also by chromatin conformation changes and epigenetic markers. Therefore, we incorporated the peaks of acetylated Histone H3 (19) and BaseJ abundance (18), which are associated with transcription initiation and termination, respectively, in our analysis. In the dSSRs and cSSRs, our PRO-seq read alignment starts and stops mostly match the H3ac and BaseJ peaks (Figure 2, 3, S2). However, when a BaseJ peak (stop) followed by an H3ac peak (start) occurs within PTUs, we cannot always observe a termination and reinitiation of transcription in our PRO-seq data. Sometimes those peaks correlate with spikes on the antisense strand that overlap with RNA Pol III or other non-coding genes. Based on the ChIP-seq data (18), we do not know which strand of DNA is modified with BaseJ and it may be that the modification is on the antisense strand and therefore does not interfere with transcription on the sense strand. Moreover, the ChIP-seq data can only indicate relative abundance of chromatin modifications, with no set maxima. Therefore, some leakage must be assumed.

H3ac and BaseJ have been used in the past to identify head-tail regions within PTUs (21), but from our data, as discussed above, we suggest that BaseJ ChIP-seq peaks alone cannot specify RNA polymerase dissociation and are not a reliable marker for defining head-tail regions within PTUs, although they were shown to correlate with transcription termination in trypanosomes (18,20,45). We also observe that the transcription stops in cSSRs do not always happen abruptly, but that RNA synthesis is sometimes gradually reduced (Figure 5 B, S4). Some cSSRs harbour RNA Pol III genes, but our PRO-seq reads cannot differentiate between RNA polymerase classes. To distinguish between those, inhibition of the run-on reaction with different concentrations of α-amanitin (46,47) may be used in future analyses, since discriminative inhibition of *Leishmania* RNA polymerase classes is established (17,41).

In the 3’ telomere regions, there is not always a BaseJ peak; transcription usually continues until the end of the chromosome, resembling run-off transcription (48) (Figure S6). In general, there is no distinct pattern of transcription termination as it varies in all cSSRs (Figure S4). The same can be observed for transcription initiation, with transcription start patterns different in all dSSRs and 5’ telomeres (Figure S3, S5).

In general, open chromatin conformation is important for the recruitment of RNA polymerases and other trans-acting protein factors in eukaryotes, with nucleosome positioning also playing a role (21). For *L. donovani* promastigotes, open chromatin could be shown for dSSRs and at RNA Pol I & III gene loci (22). Another study in *L. major* showed a well-positioned nucleosome at transcription initiation sites and around RNA Pol III gene loci, whereas in cSSRs a lower nucleosome occupancy was detected (21). Especially BaseJ and the involvement of a PJW/PP1 complex seem to play a role in the termination as mutations in different proteins that are part of this complex result in readthrough transcription in trypanosomes (20,45,49,50). In addition, histone variants have been shown to play an important role. Histone variants cause nucleosomes to become more unstable, resulting in a more dynamic chromatin (51). The histone variant H3.V from *L. major* and *L. tarentolae* co-localises with BaseJ at transcription termination sites (49,52), but the loss of H3.V alone did not cause termination of transcription (53). Furthermore, the histone variants H2A.Z and H2B.V co-localise with transcription initiation sites in *L. tarentolae*, whereas nucleosomes within PTUs usually consist of core histones and not histone variants (52). H2A.Z and H2B.V were also shown to overlap with transcription initiation in *T. brucei* (8,49), while H3.V and H4.V are enriched at termination sites (51).

### RNA polymerase Pausing

Promoter-proximal pausing has been ascribed a role in gene expression control and splicing in various eukaryotes, and often occurs close to promoter regions and polyadenylation sites (32). It was especially shown to regulate transcription of the *hsp70* gene locus in *Drosophila melanogaster* in response to heat shock as it was proposed that paused RNA polymerase II after initiation can facilitate a faster response to environmental changes (35,54). In trypanosomatids, there is some suggestion that pausing exists (8,21,55), so we decided to analyse pausing using nuclear run-on reactions with and without sarkosyl in our study. Trypanosomes, as early branching eukaryotes, contain all core RNA Pol II subunits, but lack two subunits of RNA Pol I and one of RNA Pol III (56). The mechanism of promoter-proximal pausing of RNA Pol II after transcription initiation depends on NELF and DSIF in eukaryotes, but orthologues cannot be identified in trypanosomes (57,58). Nevertheless, trypanosomes contain orthologues of the transcription elongation factor TFIIS, whose binding site is blocked by NELF so that the RNA polymerase cannot elongate (59). We observe a slightly higher read occupancy in regions of transcription initiation at dSSRs and 5’telomeres, but no distinct peak was observed (Figure 6 A, D, F). In *D. melonagaster*, some genes have been shown to have a focused peak for pausing, while others have a more dispersed pausing pattern (35). In *T. brucei*, a pronounced peak 100-200 bp downstream of the transcription start on chromosome 9 was observed using primary transcription sequencing (8). Even if we do not see a distinct peak, we can exclude that sarkosyl generally leads to more transcription, as we observe a higher read occupancy within the PTUs without sarkosyl.

Furthermore, we observe a significant increase of reads within RNA Pol I and III transcribed gene loci in samples with sarkosyl indicating a release of paused RNA polymerase I and III complexes (Figure 6 H, I). Pausing has been described to support processing of rRNA co-transcriptionally via the control of RNA polymerase I progression in yeast as this is important for ribosome assembly (60–62). We conclude that we found further evidence for the existence of RNA polymerase pausing, although the mechanism of RNA polymerase pausing by RNA Pol II and especially for RNA Pol I and III in trypanosomes remains unclear. To further analyse pausing, a single-basepair resolution of the position of RNA Polymerase complexes is necessary, which can be achieved by performing four nuclear run-on experiments using a different biotinylated nucleoside triphosphates in each reaction. Also, investigating different environmental conditions e.g. heat shock, would help to observe changes in the pausing pattern as we only analysed *L. major* promastigotes under logarithmic growth conditions.

## Acknowledgements

We thank the NGS facility at the BNITM for sequencing our prepared PRO-seq libraries. We further thank all members of the Leishmania Genetics Group for fruitful discussions and suggestions.

## Data Availability

All PRO-seq read files are made available at the Sequence Read Archive of NCBI under the link: https://www.ncbi.nlm.nih.gov/bioproject/PRJNA1020779

## Author contributions

JG: methodology, data analysis, conceptualisation, figure preparation, manuscript: writing and editing of original draft. SL: bioinformatic analysis, figure preparation, manuscript: reviewing. JC: supervision, conceptualisation, manuscript: writing and editing of original draft.

## Notes

https://www.ncbi.nlm.nih.gov/bioproject/PRJNA1020779

### Competing Interest Statement

The authors have declared no competing interest.

## Bibliography

1. Alvar, J., Velez, I.D., Bern, C., Herrero, M., Desjeux, P., Cano, J., Jannin, J. and den Boer, M. (2012) Leishmaniasis worldwide and global estimates of its incidence. PloS one, 7, e35671.

2. WHO. (2010) First WHO report on neglected tropical diseases: working to overcome the global impact of neglected tropical diseases. WHO Press, Geneva, Switzerland.

3. Hendrickx, S., Maes, L., Croft, S.L. and Caljon, G. (2018) In A., P.-S. and M., P.-N. (eds.), Drug Resistance in Leishmania Parasites. Springer, Cham.

4. Ivens, A.C., Peacock, C.S., Worthey, E.A., Murphy, L., Aggarwal, G., Berriman, M., Sisk, E., Rajandream, M.A., Adlem, E., Aert, R. et al. (2005) The genome of the kinetoplastid parasite, Leishmania major. Science, 309, 436–442.

5. Peacock, C.S., Seeger, K., Harris, D., Murphy, L., Ruiz, J.C., Quail, M.A., Peters, N., Adlem, E., Tivey, A., Aslett, M. et al. (2007) Comparative genomic analysis of three Leishmania species that cause diverse human disease. Nat Genet, 39, 839–847.

6. Clayton, C.E. (2016) Gene expression in Kinetoplastids. Curr Opin Microbiol, 32, 46–51.

7. Grunebast, J. and Clos, J. (2020) Leishmania: Responding to environmental signals and challenges without regulated transcription. Comput Struct Biotechnol J, 18, 4016–4023.

8. Wedel, C., Forstner, K.U., Derr, R. and Siegel, T.N. (2017) GT-rich promoters can drive RNA pol II transcription and deposition of H2A.Z in African trypanosomes. EMBO J, 36, 2581–2594.

9. Cordon-Obras, C., Gomez-Linan, C., Torres-Rusillo, S., Vidal-Cobo, I., Lopez-Farfan, D., Barroso-Del Jesus, A., Rojas-Barros, D., Carrington, M. and Navarro, M. (2022) Identification of sequence-specific promoters driving polycistronic transcription initiation by RNA polymerase II in trypanosomes. Cell Rep, 38, 110221.

10. Brandau, S., Dresel, A. and Clos, J. (1995) High constitutive levels of heat-shock proteins in human-pathogenic parasites of the genus Leishmania. Biochem J, 310, 225–232.

11. Dresel, A. and Clos, J. (1997) Transcription of the Leishmania major Hsp70-I gene locus does not proceed through the noncoding region. Exp Parasitol, 86, 206–212.

12. Quijada, L., Soto, M., Alonso, C. and Requena, J.M. (1997) Analysis of post-transcriptional regulation operating on transcription products of the tandemly linked Leishmania infantum hsp70 genes. J Biol Chem, 272, 4493–4499.

13. Wiesgigl, M. and Clos, J. (1999) Uniform distribution of transcription complexes on the clpB gene locus of *Leishmania donovani*. Protist, 150, 369–373.

14. Monnerat, S., Martinez-Calvillo, S., Worthey, E., Myler, P.J., Stuart, K.D. and Fasel, N. (2004) Genomic organization and gene expression in a chromosomal region of Leishmania major. Mol Biochem Parasitol, 134, 233–243.

15. Martinez-Calvillo, S., Yan, S., Nguyen, D., Fox, M., Stuart, K. and Myler, P.J. (2003) Transcription of Leishmania major Friedlin chromosome 1 initiates in both directions within a single region. Mol Cell, 11, 1291–1299.

16. Worthey, E.A., Martinez-Calvillo, S., Schnaufer, A., Aggarwal, G., Cawthra, J., Fazelinia, G., Fong, C., Fu, G., Hassebrock, M., Hixson, G. et al. (2003) Leishmania major chromosome 3 contains two long convergent polycistronic gene clusters separated by a tRNA gene. Nucleic Acids Res, 31, 4201–4210.

17. Martinez-Calvillo, S., Nguyen, D., Stuart, K. and Myler, P.J. (2004) Transcription initiation and termination on Leishmania major chromosome 3. Eukaryot Cell, 3, 506–517.

18. van Luenen, H.G., Farris, C., Jan, S., Genest, P.A., Tripathi, P., Velds, A., Kerkhoven, R.M., Nieuwland, M., Haydock, A., Ramasamy, G., et al. (2012) Glucosylated hydroxymethyluracil, DNA base J, prevents transcriptional readthrough in Leishmania. Cell, 150, 909–921.

19. Thomas, S., Green, A., Sturm, N.R., Campbell, D.A. and Myler, P.J. (2009) Histone acetylations mark origins of polycistronic transcription in Leishmania major. BMC Genomics, 10, 152.

20. Reynolds, D., Cliffe, L., Forstner, K.U., Hon, C.C., Siegel, T.N. and Sabatini, R. (2014) Regulation of transcription termination by glucosylated hydroxymethyluracil, base J, in Leishmania major and Trypanosoma brucei. Nucleic Acids Res, 42, 9717–9729.

21. Lombrana, R., Alvarez, A., Fernandez-Justel, J.M., Almeida, R., Poza-Carrion, C., Gomes, F., Calzada, A., Requena, J.M. and Gomez, M. (2016) Transcriptionally Driven DNA Replication Program of the Human Parasite Leishmania major. Cell Rep, 16, 1774–1786.

22. Grunebast, J., Lorenzen, S., Zummack, J. and Clos, J. (2021) Life Cycle Stage-Specific Accessibility of Leishmania donovani Chromatin at Transcription Start Regions. mSystems, 6, e0062821.

23. Mahat, D.B., Kwak, H., Booth, G.T., Jonkers, I.H., Danko, C.G., Patel, R.K., Waters, C.T., Munson, K., Core, L.J. and Lis, J.T. (2016) Base-pair-resolution genome-wide mapping of active RNA polymerases using precision nuclear run-on (PRO-seq). Nat Protoc, 11, 1455–1476.

24. Smith, J.P., Dutta, A.B., Sathyan, K.M., Guertin, M.J. and Sheffield, N.C. (2021) PEPPRO: quality control and processing of nascent RNA profiling data. Genome Biol, 22, 155.

25. Martin, M. (2011) Cutadapt removes adapter sequences from high-throughput sequencing reads. EMBnet. journal, 17(1), 10–12.

26. Camacho, E., Gonzalez-de la Fuente, S., Solana, J.C., Rastrojo, A., Carrasco-Ramiro, F., Requena, J.M. and Aguado, B. (2021) Gene Annotation and Transcriptome Delineation on a De Novo Genome Assembly for the Reference Leishmania major Friedlin Strain. Genes (Basel), 12.

27. Langmead, B. and Salzberg, S.L. (2012) Fast gapped-read alignment with Bowtie 2. Nat Methods, 9, 357–359.

28. Quinlan, A.R. and Hall, I.M. (2010) BEDTools: a flexible suite of utilities for comparing genomic features. Bioinformatics, 26, 841–842.

29. Langmead, B., Trapnell, C., Pop, M. and Salzberg, S.L. (2009) Ultrafast and memory-efficient alignment of short DNA sequences to the human genome. Genome Biol, 10, R25.

30. Core, L.J., Waterfall, J.J., Gilchrist, D.A., Fargo, D.C., Kwak, H., Adelman, K. and Lis, J.T. (2012) Defining the status of RNA polymerase at promoters. Cell Rep, 2, 1025–1035.

31. Wissink, E.M., Vihervaara, A., Tippens, N.D. and Lis, J.T. (2019) Nascent RNA analyses: tracking transcription and its regulation. Nat Rev Genet, 20, 705–723.

32. Adelman, K. and Lis, J.T. (2012) Promoter-proximal pausing of RNA polymerase II: emerging roles in metazoans. Nat Rev Genet, 13, 720–731.

33. Core, L.J., Waterfall, J.J. and Lis, J.T. (2008) Nascent RNA sequencing reveals widespread pausing and divergent initiation at human promoters. Science, 322, 1845–1848.

34. Churchman, L.S. and Weissman, J.S. (2011) Nascent transcript sequencing visualizes transcription at nucleotide resolution. Nature, 469, 368–373.

35. Kwak, H., Fuda, N.J., Core, L.J. and Lis, J.T. (2013) Precise maps of RNA polymerase reveal how promoters direct initiation and pausing. Science, 339, 950–953.

36. Button, L.L., Russell, D.G., Klein, H.L., Medina-Acosta, E., Karess, R.E. and McMaster, W.R. (1989) Genes encoding the major surface glycoprotein in Leishmania are tandemly linked at a single chromosomal locus and are constitutively transcribed. Mol Biochem Parasitol, 32, 271–283.

37. Evers, R. and Cornelissen, A.W. (1990) The Trypanosoma brucei protein phosphatase gene: polycistronic transcription with the RNA polymerase II largest subunit gene. Nucleic Acids Res, 18, 5089–5095.

38. Flinn, H.M. and Smith, D.F. (1992) Genomic organisation and expression of a differentially-regulated gene family from Leishmania major. Nucleic Acids Res, 20, 755–762.

39. Wiesgigl, M. and Clos, J. (1999) Uniform distribution of transcription complexes over the entire Leishmania donovani clpB (hsp 100) gene locus. Protist, 150, 369–373.

40. Mishra, K.K., Holzer, T.R., Moore, L.L. and LeBowitz, J.H. (2003) A negative regulatory element controls mRNA abundance of the Leishmania mexicana Paraflagellar rod gene PFR2. Eukaryot Cell, 2, 1009–1017.

41. Adhya, S., Das, S. and Bhaumik, M. (1990) Transcription and processing of ß-tubulin messenger RNA in Leishmania donovani promastigotes. J. Biosci., Vol 15, pp. 249–259.

42. Siegel, T.N., Hekstra, D.R., Wang, X., Dewell, S. and Cross, G.A. (2010) Genome-wide analysis of mRNA abundance in two life-cycle stages of Trypanosoma brucei and identification of splicing and polyadenylation sites. Nucleic Acids Res, 38, 4946–4957.

43. Dumetz, F., Imamura, H., Sanders, M., Seblova, V., Myskova, J., Pescher, P., Vanaerschot, M., Meehan, C.J., Cuypers, B., De Muylder, G., et al. (2017) Modulation of Aneuploidy in Leishmania donovani during Adaptation to Different In Vitro and In Vivo Environments and Its Impact on Gene Expression. mBio, 8.

44. Clayton, C.E. (2002) Life without transcriptional control? From fly to man and back again. Embo J, 21, 1881–1888.

45. Ekanayake, D. and Sabatini, R. (2011) Epigenetic regulation of polymerase II transcription initiation in Trypanosoma cruzi: modulation of nucleosome abundance, histone modification, and polymerase occupancy by O-linked thymine DNA glucosylation. Eukaryot Cell, 10, 1465–1472.

46. Sugden, B. and Keller, W. (1973) Mammalian deoxyribonucleic acid-dependent ribonucleic acid polymerases. I. Purification and properties of an -amanitin-sensitive ribonucleic acid polymerase and stimulatory factors from HeLa and KB cells. J Biol Chem, 248, 3777–3788.

47. Schultz, L.D. and Hall, B.D. (1976) Transcription in yeast: alpha-amanitin sensitivity and other properties which distinguish between RNA polymerases I and III. Proc Natl Acad Sci U S A, 73, 1029–1033.

48. Weil, P.A., Luse, D.S., Segall, J. and Roeder, R.G. (1979) Selective and accurate initiation of transcription at the Ad2 major late promotor in a soluble system dependent on purified RNA polymerase II and DNA. Cell, 18, 469–484.

49. Reynolds, D.L., Hofmeister, B.T., Cliffe, L., Siegel, T.N., Anderson, B.A., Beverley, S.M., Schmitz, R.J. and Sabatini, R. (2016) Base J represses genes at the end of polycistronic gene clusters in Leishmania major by promoting RNAP II termination. Mol Microbiol, 101, 559–574.

50. Kieft, R., Zhang, Y., Yan, H., Schmitz, R.J. and Sabatini, R. (2023) Knockout of protein phosphatase 1 in Leishmania major reveals its role during RNA polymerase II transcription termination. Nucleic Acids Res, 51, 6208–6226.

51. Siegel, T.N., Hekstra, D.R., Kemp, L.E., Figueiredo, L.M., Lowell, J.E., Fenyo, D., Wang, X., Dewell, S. and Cross, G.A. (2009) Four histone variants mark the boundaries of polycistronic transcription units in Trypanosoma brucei. Genes Dev, 23, 1063–1076.

52. McDonald, J.R., Jensen, B.C., Sur, A., Wong, I.L.K., Beverley, S.M. and Myler, P.J. (2022) Localization of Epigenetic Markers in Leishmania Chromatin. Pathogens, 11.

53. Anderson, B.A., Wong, I.L., Baugh, L., Ramasamy, G., Myler, P.J. and Beverley, S.M. (2013) Kinetoplastid-specific histone variant functions are conserved in Leishmania major. Mol Biochem Parasitol, 191, 53–57.

54. O’Brien, T. and Lis, J.T. (1991) RNA polymerase II pauses at the 5’ end of the transcriptionally induced Drosophila hsp70 gene. Mol Cell Biol, 11, 5285–5290.

55. Das, A., Li, H., Liu, T. and Bellofatto, V. (2006) Biochemical characterization of Trypanosoma brucei RNA polymerase II. Mol Biochem Parasitol, 150, 201–210.

56. Kelly, S., Wickstead, B. and Gull, K. (2005) An in silico analysis of trypanosomatid RNA polymerases: insights into their unusual transcription. Biochem Soc Trans, 33, 1435–1437.

57. Li, J., Liu, Y., Rhee, H.S., Ghosh, S.K., Bai, L., Pugh, B.F. and Gilmour, D.S. (2013) Kinetic competition between elongation rate and binding of NELF controls promoter-proximal pausing. Mol Cell, 50, 711–722.

58. Florini, F., Naguleswaran, A., Gharib, W.H., Bringaud, F. and Roditi, I. (2019) Unexpected diversity in eukaryotic transcription revealed by the retrotransposon hotspot family of Trypanosoma brucei. Nucleic Acids Res, 47, 1725–1739.

59. Core, L. and Adelman, K. (2019) Promoter-proximal pausing of RNA polymerase II: a nexus of gene regulation. Genes Dev, 33, 960–982.

60. Schneider, D.A., Michel, A., Sikes, M.L., Vu, L., Dodd, J.A., Salgia, S., Osheim, Y.N., Beyer, A.L. and Nomura, M. (2007) Transcription elongation by RNA polymerase I is linked to efficient rRNA processing and ribosome assembly. Mol Cell, 26, 217–229.

61. Huffines, A.K., Engel, K.L., French, S.L., Zhang, Y., Viktorovskaya, O.V. and Schneider, D.A. (2022) Rate of transcription elongation and sequence-specific pausing by RNA polymerase I directly influence rRNA processing. J Biol Chem, 298, 102730.

62. Duval, M., Yague-Sanz, C., Turowski, T.W., Petfalski, E., Tollervey, D. and Bachand, F. (2023) The conserved RNA-binding protein Seb1 promotes cotranscriptional ribosomal RNA processing by controlling RNA polymerase I progression. Nat Commun, 14, 3013.

