## Supplementary Figure Legends for "Genome-wide Quantification of Polycistronic Transcription in *Leishmania major*"

**Figure S1:** Read length analysis for PRO-seq reads from 6 reactions.

**Figure S2:** Linear plot of PRO-seq reads against all 36 chromosomes of the *L. major* Friedlin 2021 genome. From top to bottom, the plots show BaseJ (blue) (van Luenen et al., 2012) and H3ac (orange) (Thomas et al., 2009), PRO-seq read densities from nuclei without sarkosyl (upper and lower strands), PRO-seq read densities from nuclei without sarkosyl (upper and lower strands), and *L. major* genes with UTRs at half-height and CDSs at full height. Genes in grey: non coding RNAs, genes in blue: RNA Pol III transcribed genes.

**Figure S3: Transcription starts at dSSRs.** Distribution of normalised PRO-seq reads from *L. major* nuclei incubated with sarkosyl (dark red outline) or without sarkosyl (light red shading) around dSSR regions. For chromosomal locations of dSSRs (grey shade), see Table S2. Blue shade: RNA Pol III transcribed genes.

**Figure S4: Transcription stops at cSSRs.** Distribution of normalised PRO-seq reads from *L. major* nuclei incubated with sarkosyl (dark red outline) or without sarkosyl (light red shading) around cSSR regions. For chromosomal locations of cSSRs (grey shade), see Table S2. Blue shade: RNA Pol III transcribed genes.

**Figure S5: Transcription starts at 5'-telomer ends.** Distribution of normalised PRO-seq reads from *L. major* nuclei incubated with sarkosyl (dark red outline) or without sarkosyl (light red shading) at 5'-telomeric regions. For chromosomal locations of 5' telomers (grey shade), see Table S2.

**Figure S6: Transcription at 3'-telomer ends**. Distribution of normalised PRO-seq reads from *L. major* nuclei incubated with sarkosyl (dark red outline) or without sarkosyl (light red shading) around 3'-telomeric regions. For chromosomal locations of 3' telomers (grey shade), see Table S2.
