## Supplementary figures and images for "Genome-wide Quantification of Polycistronic Transcription in *Leishmania major*"

### Figure S1

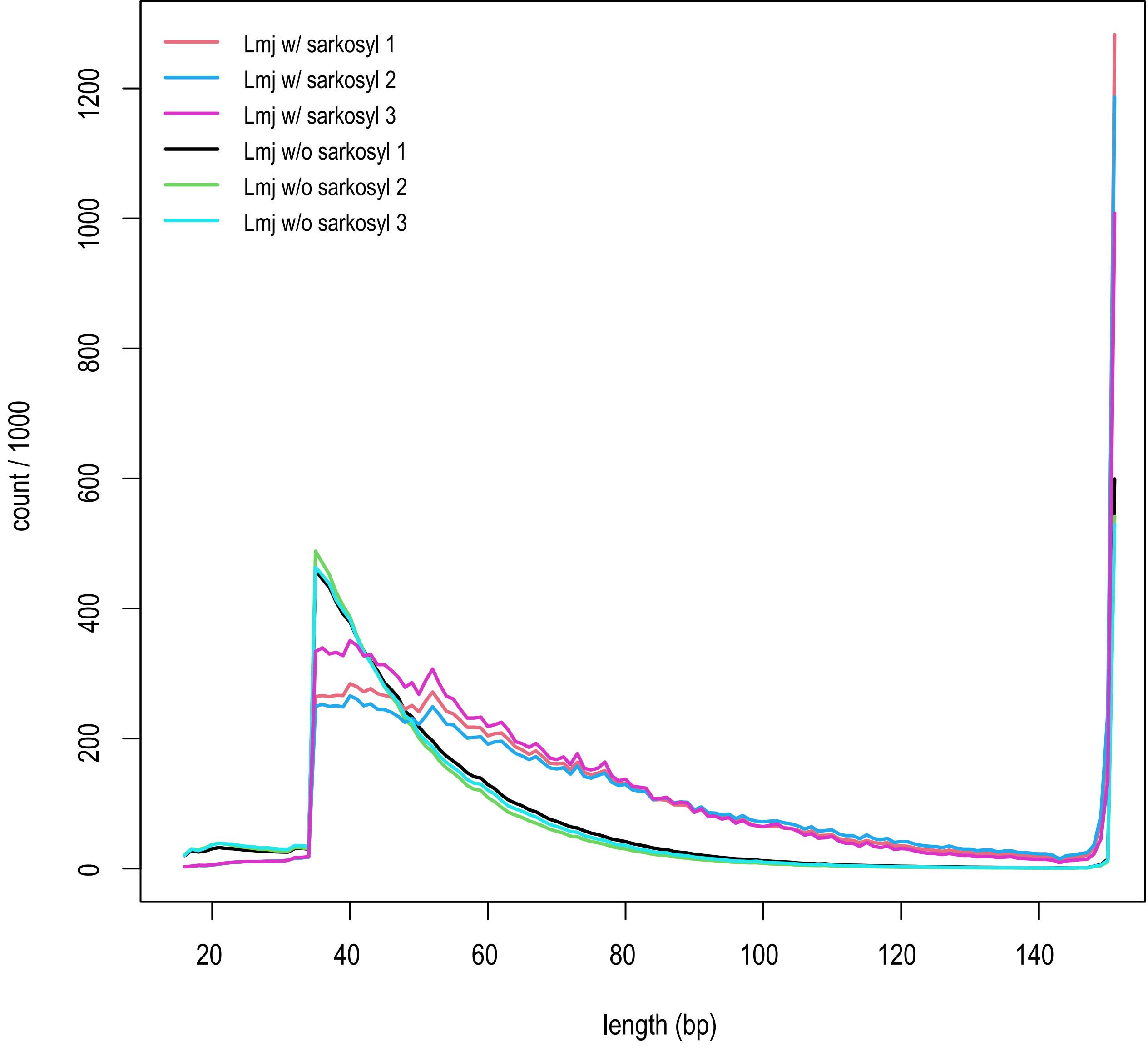

### Figure S3

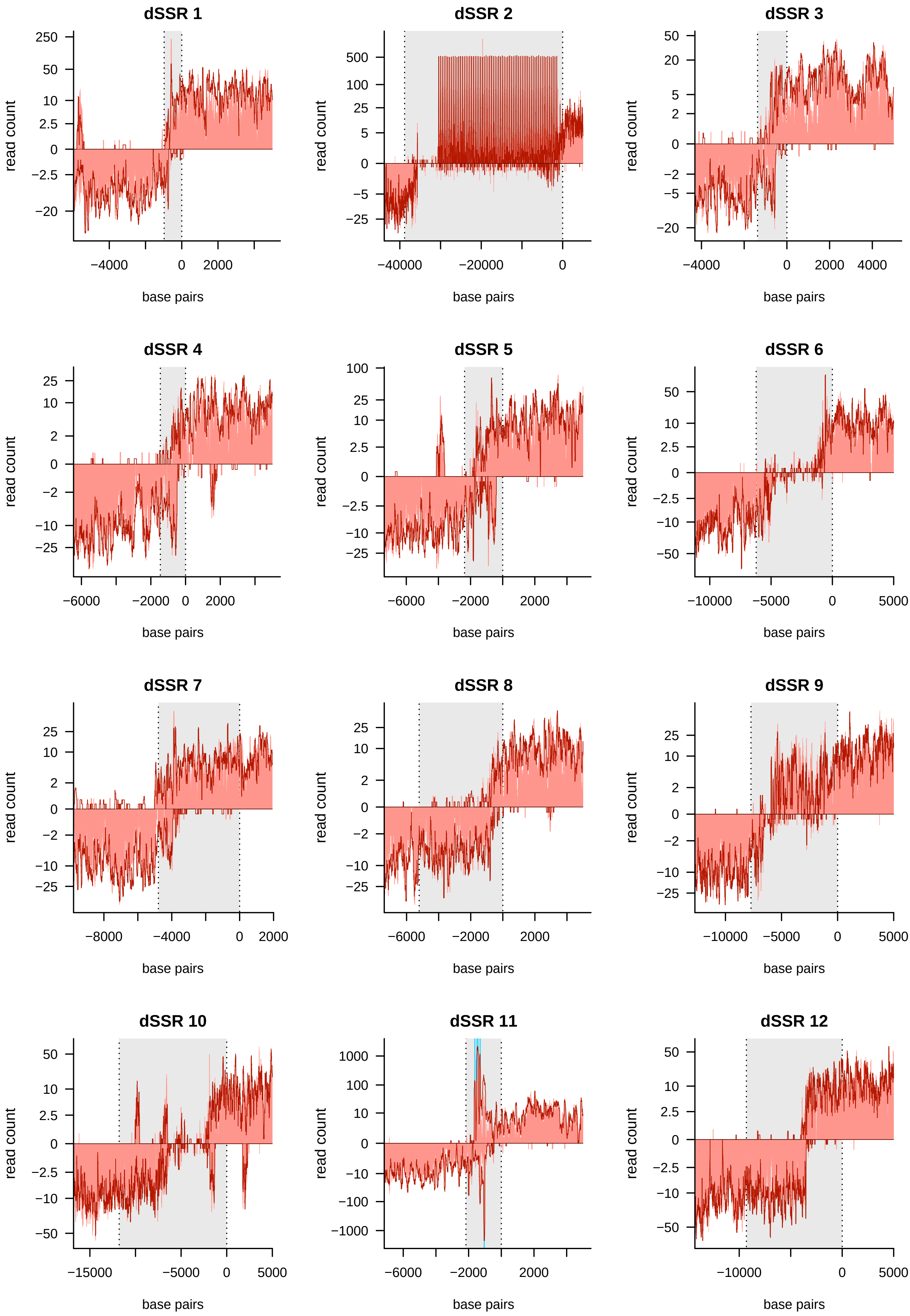

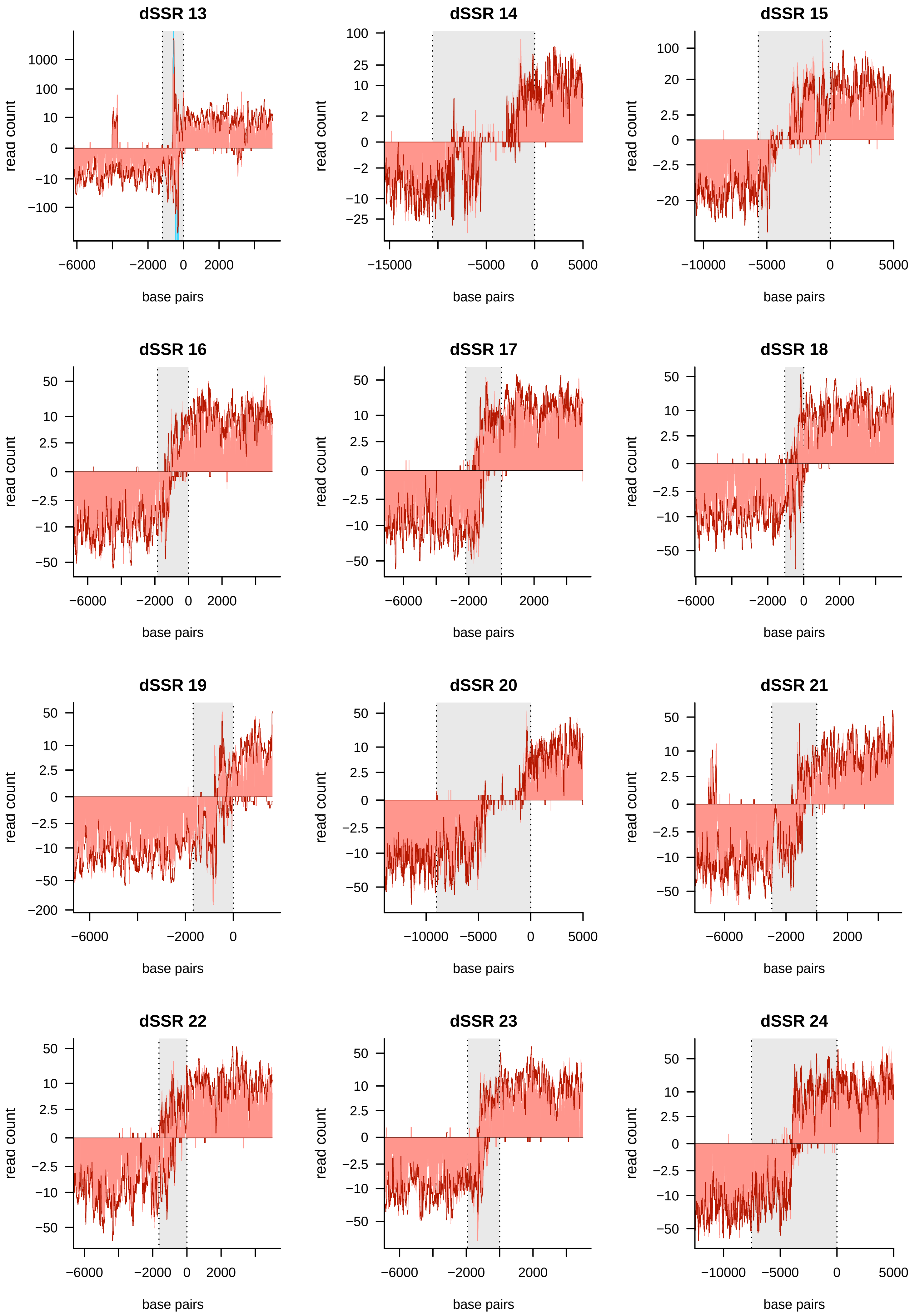

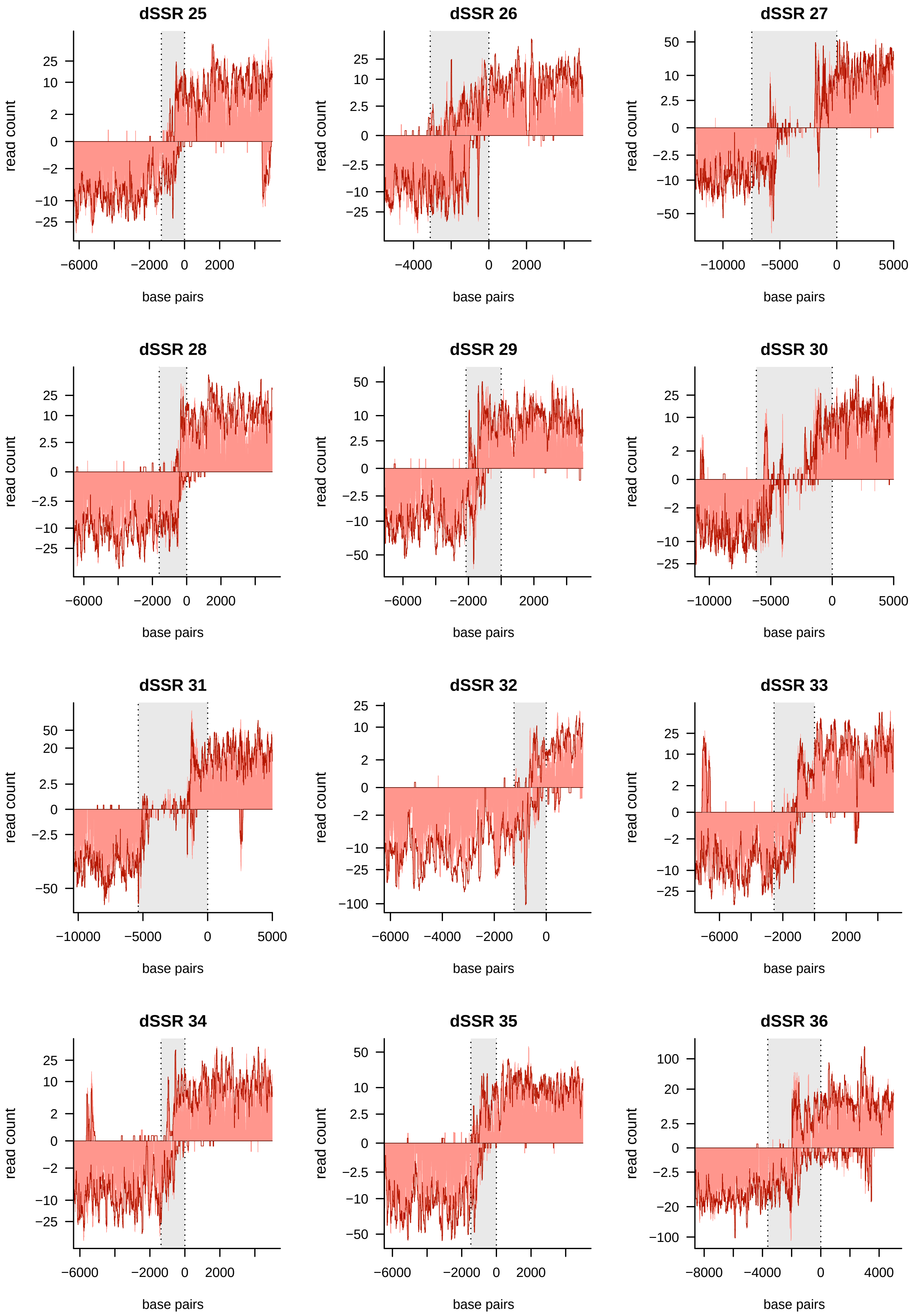

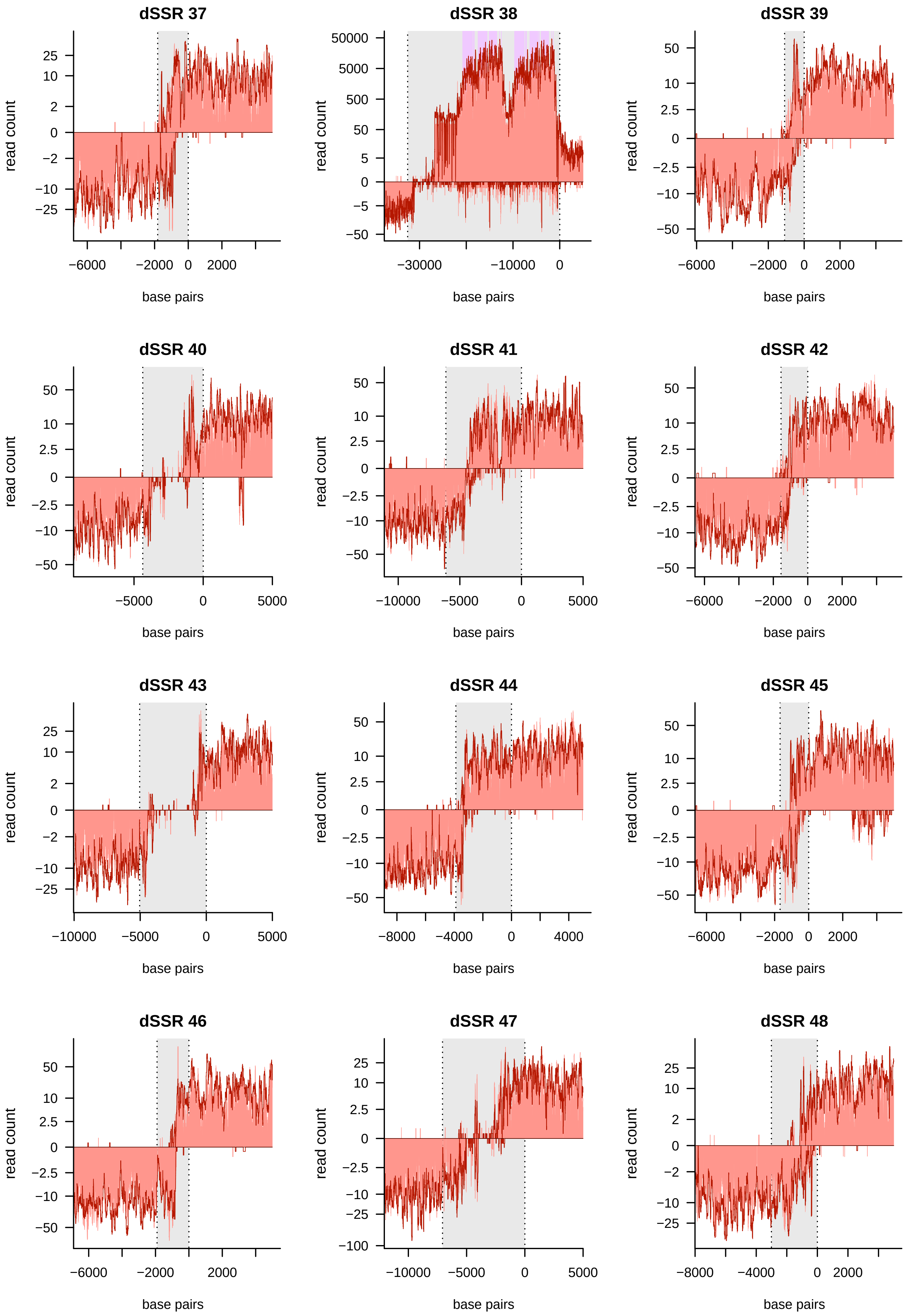

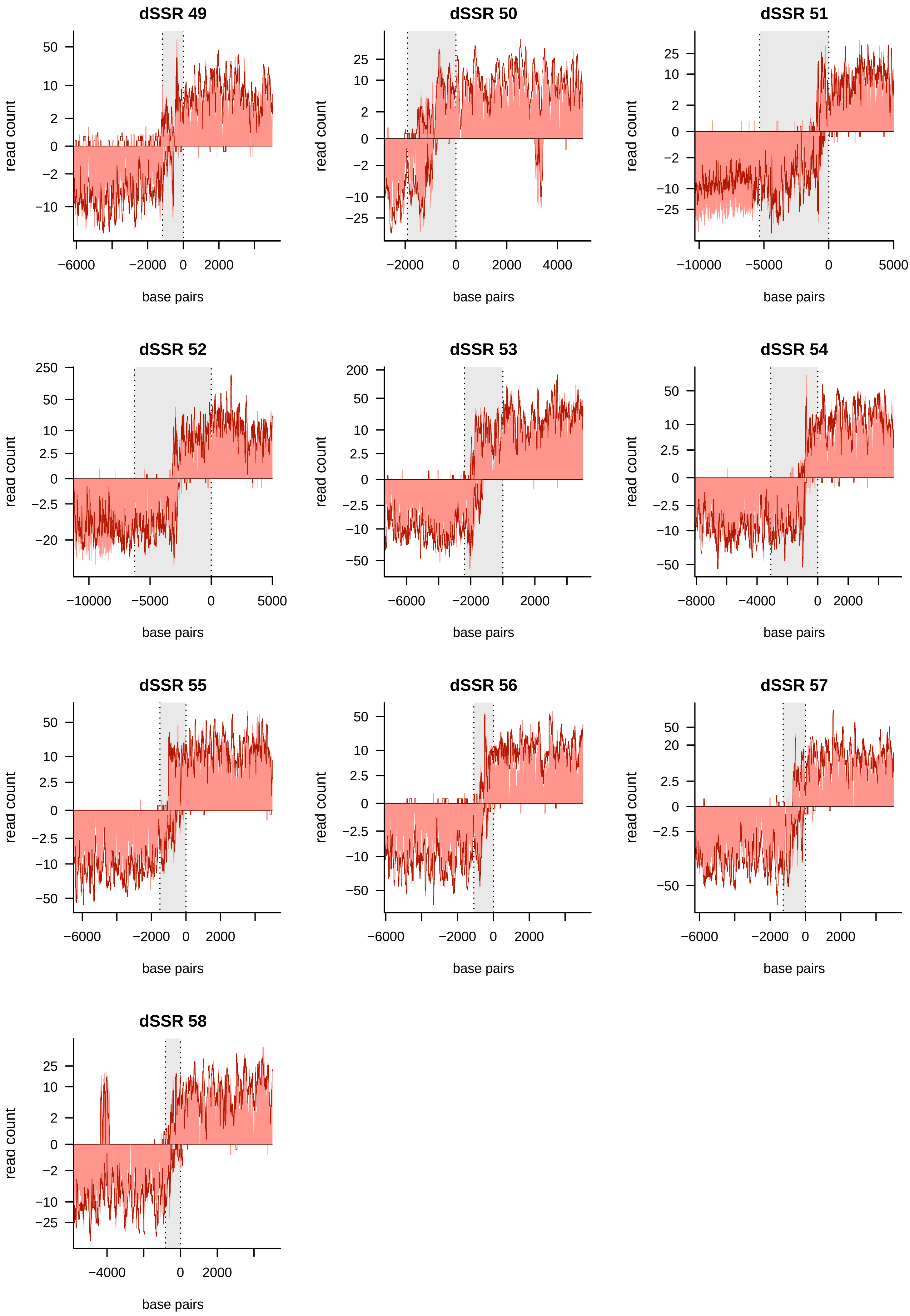

### Figure S4

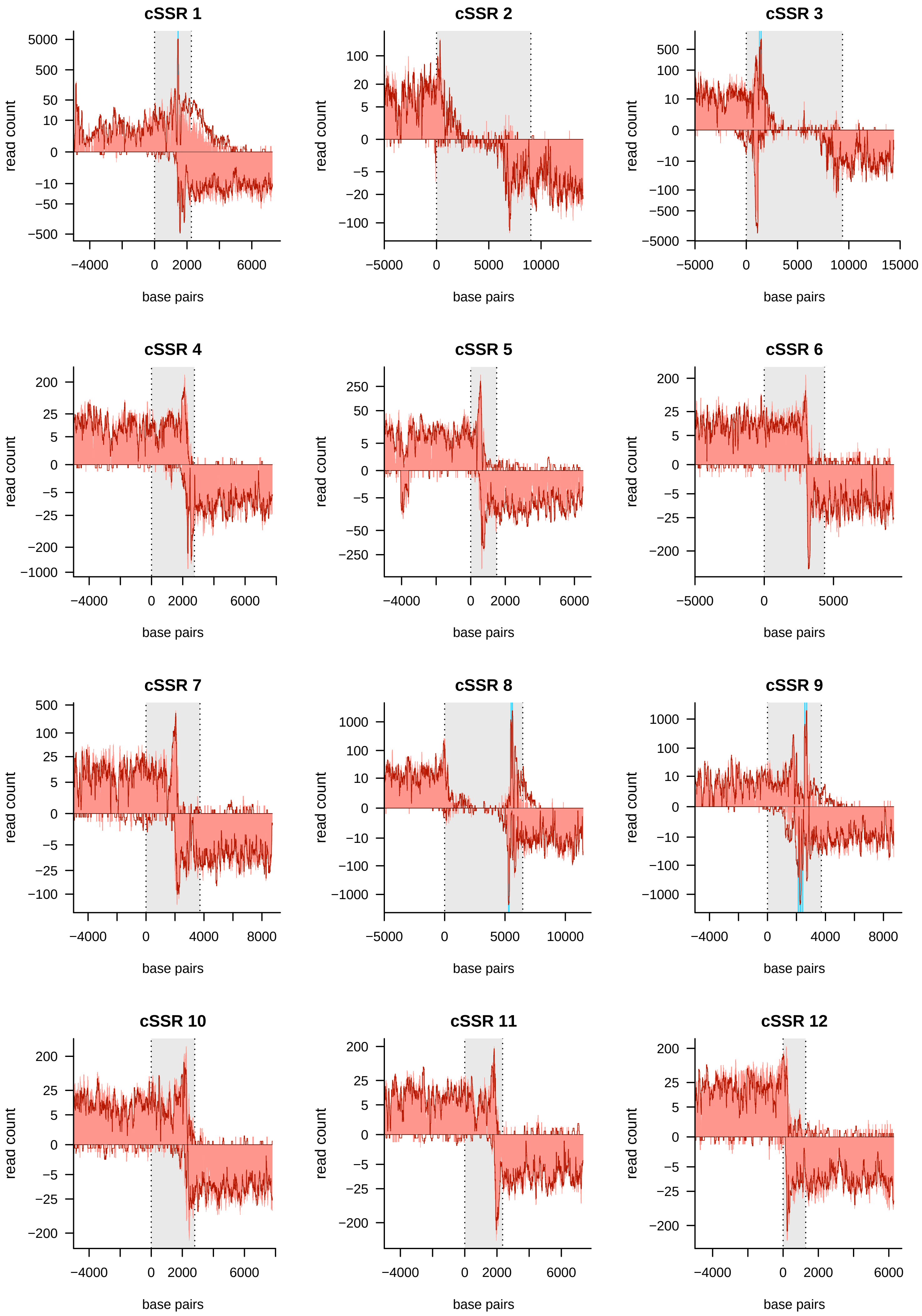

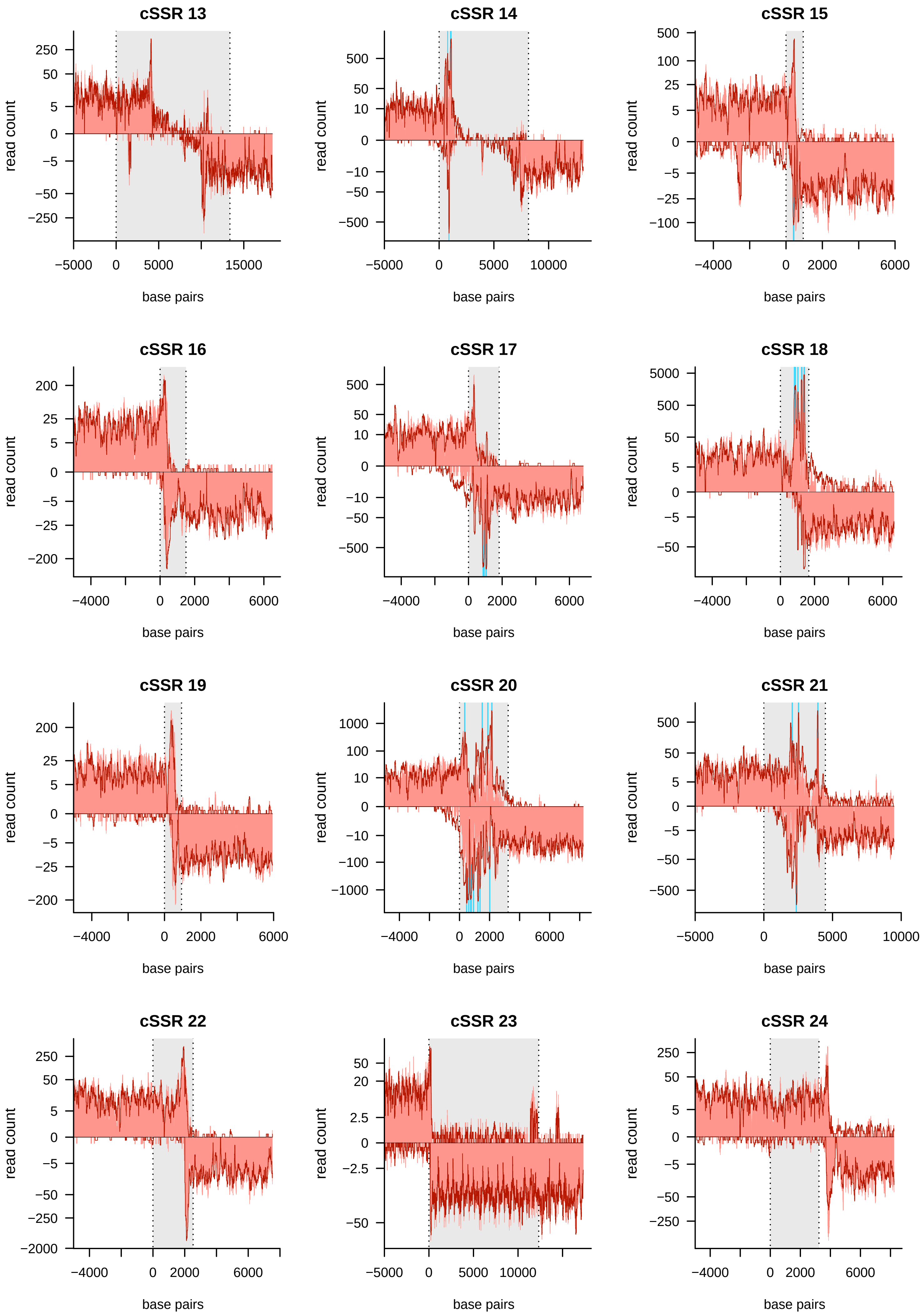

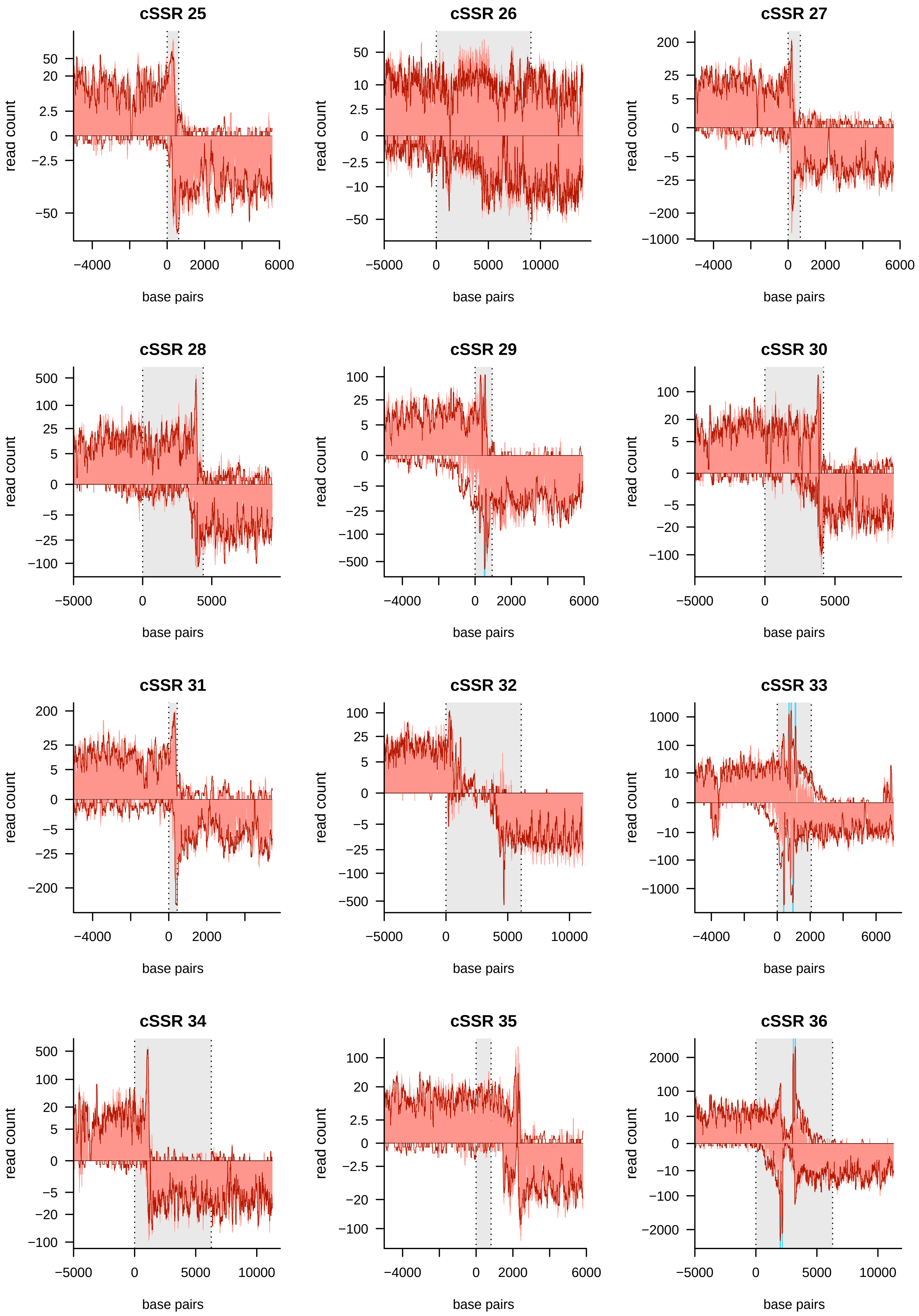

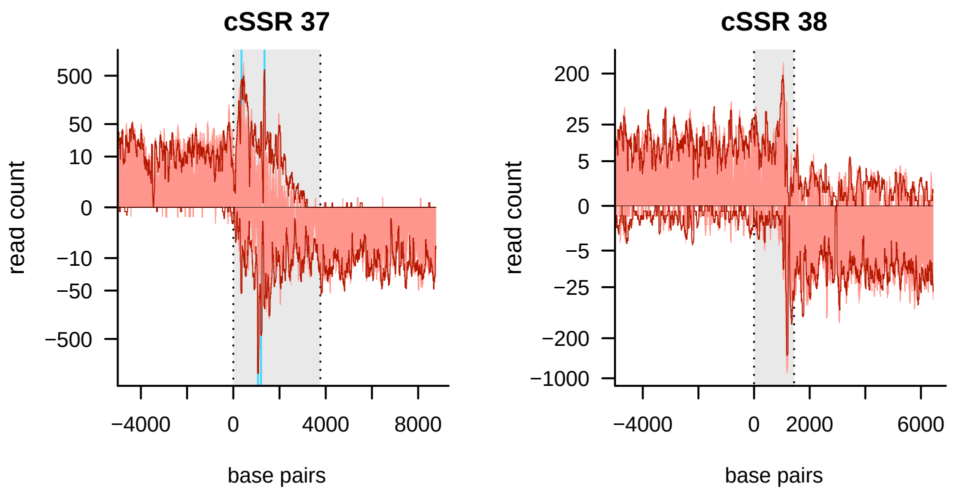

### Figure S5

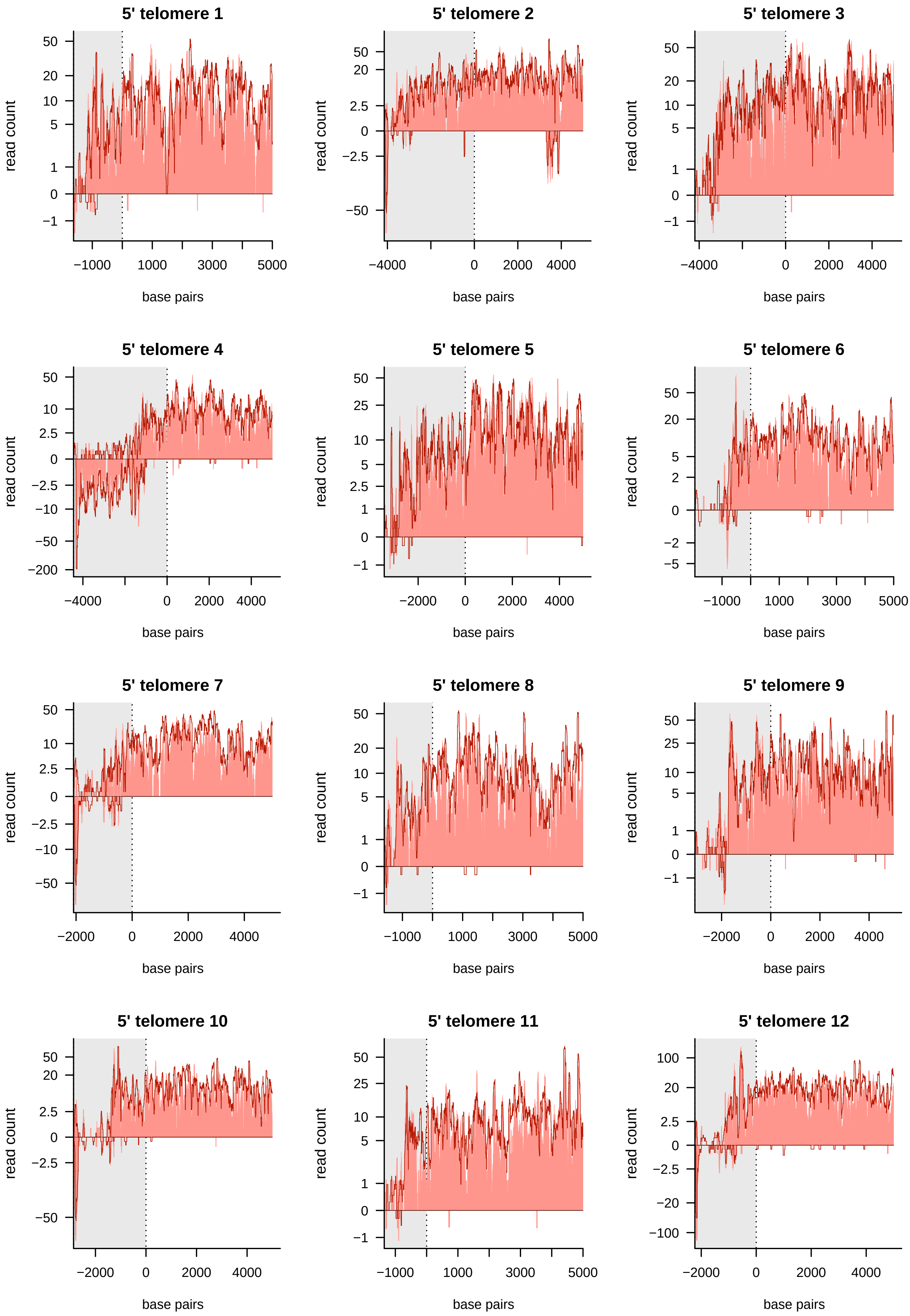

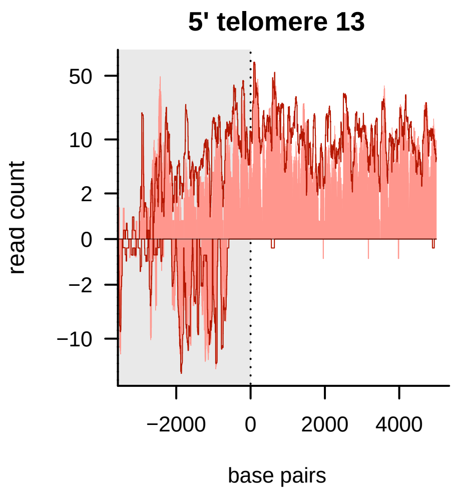

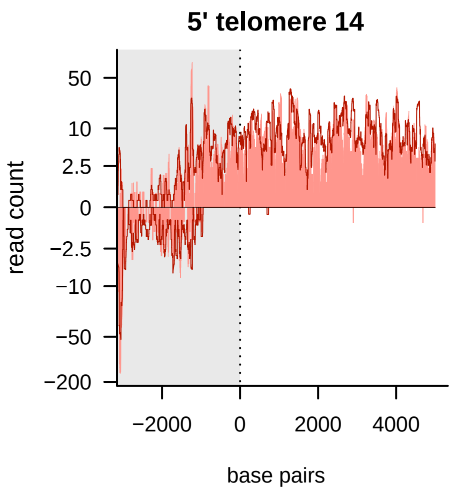

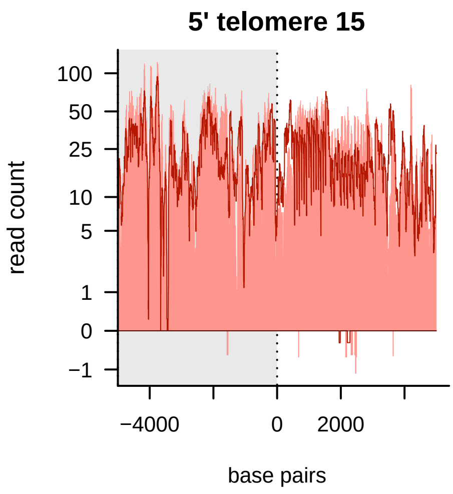

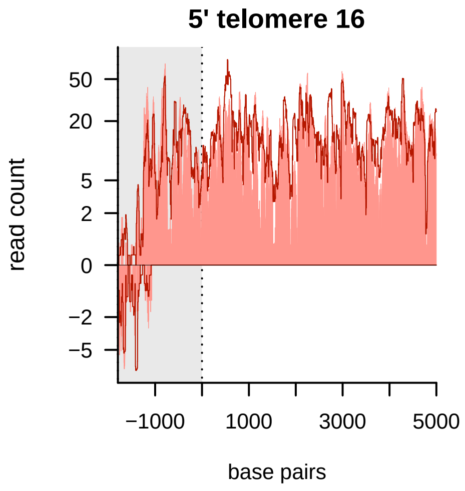

### Figure S6

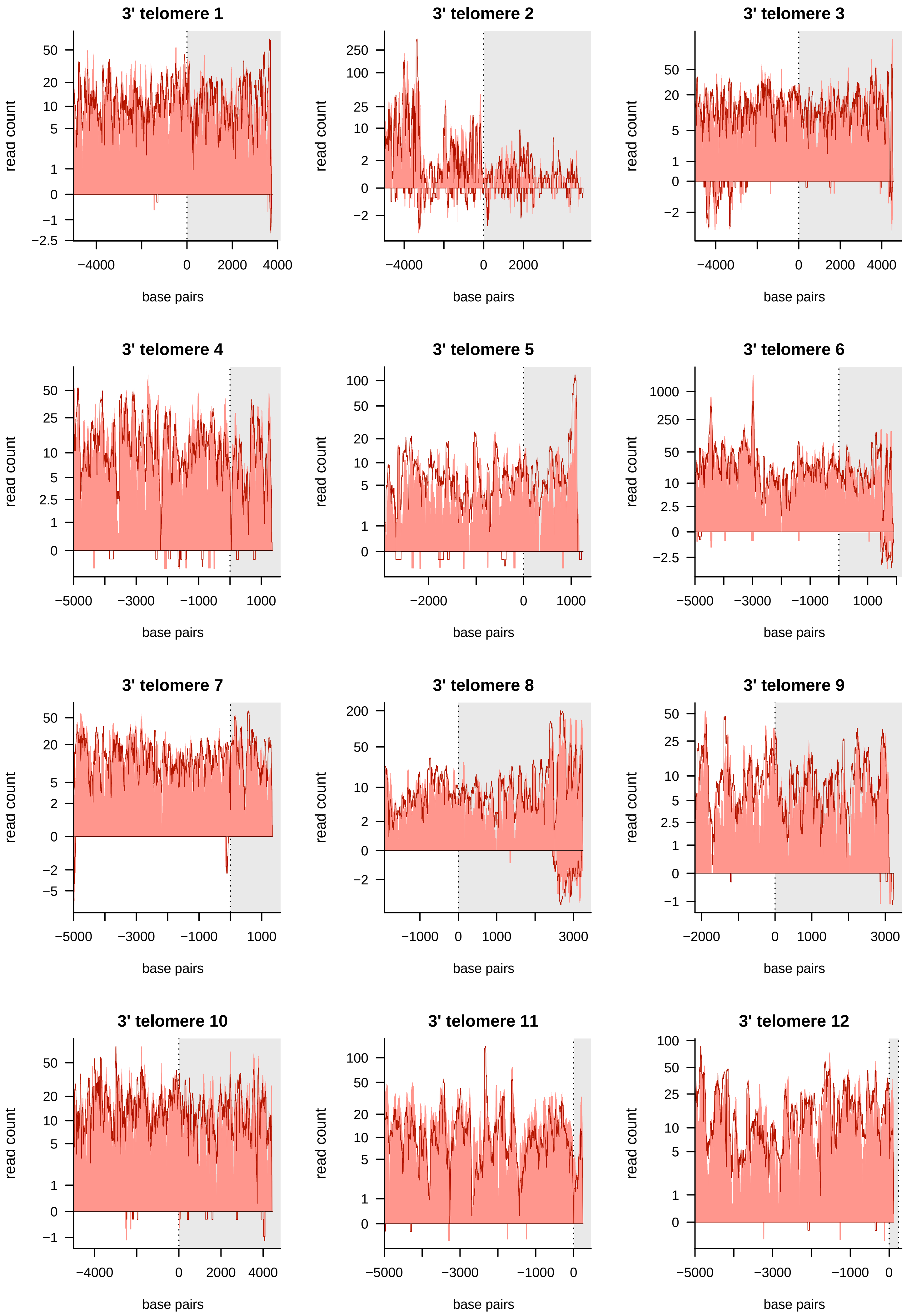

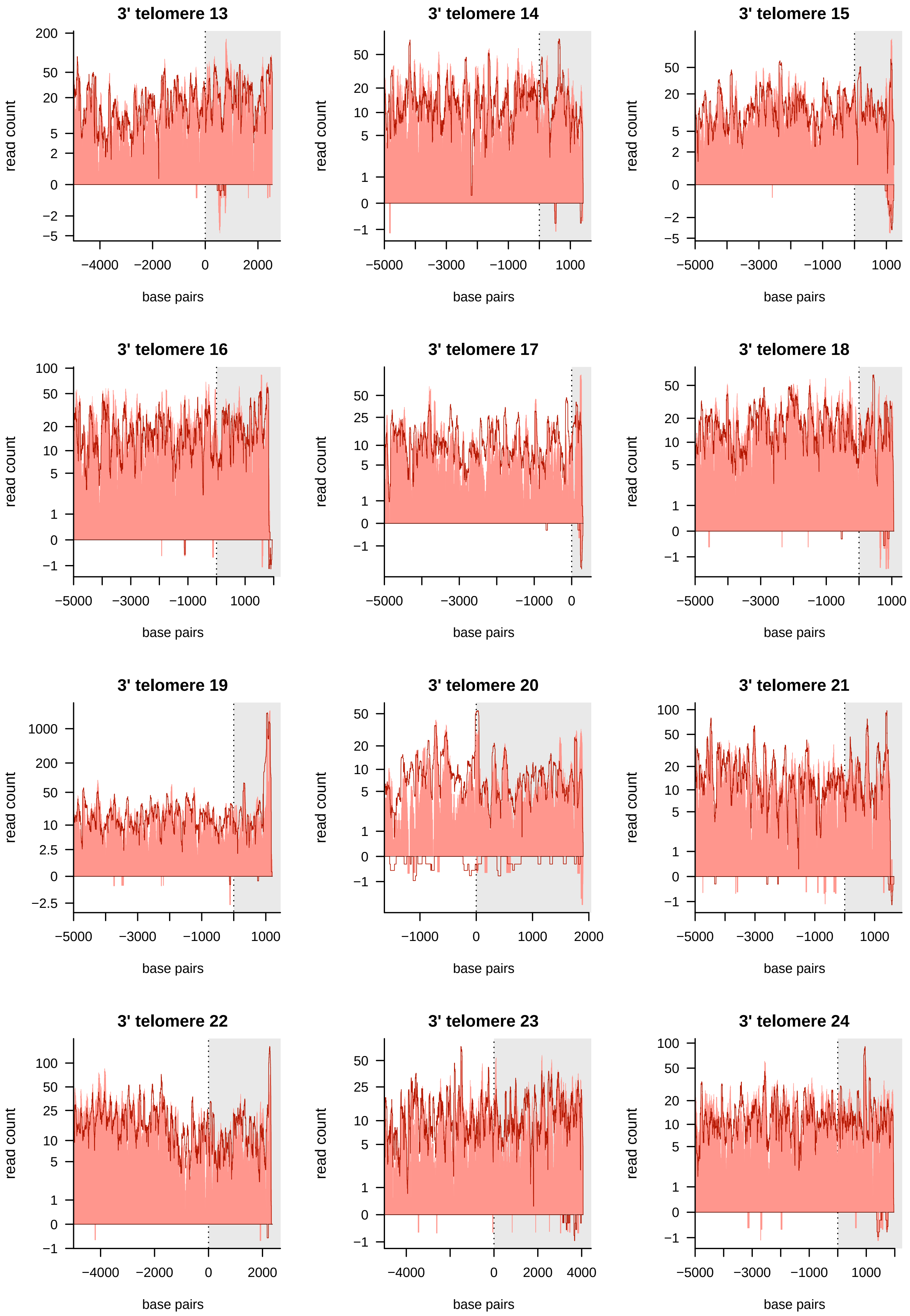

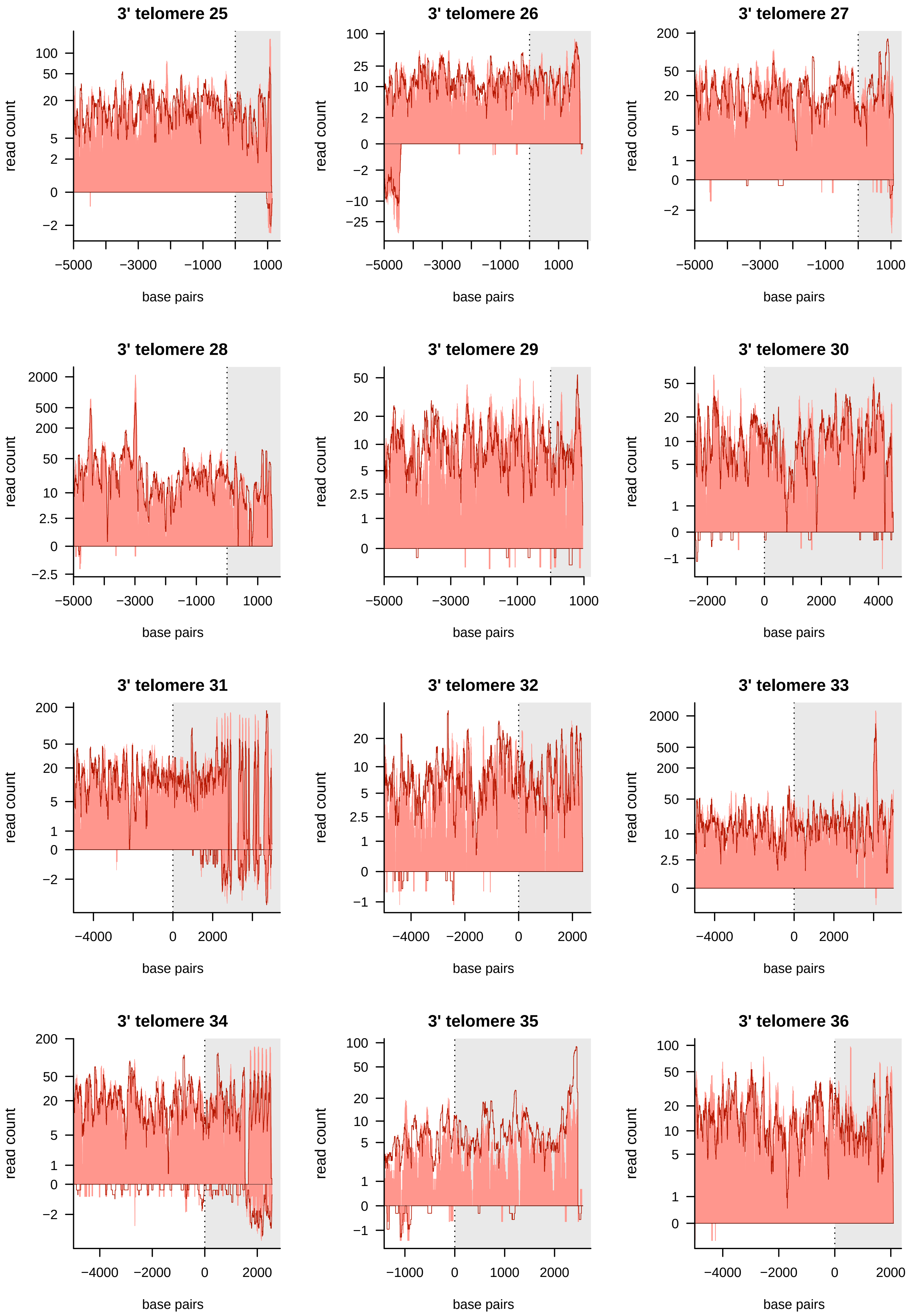

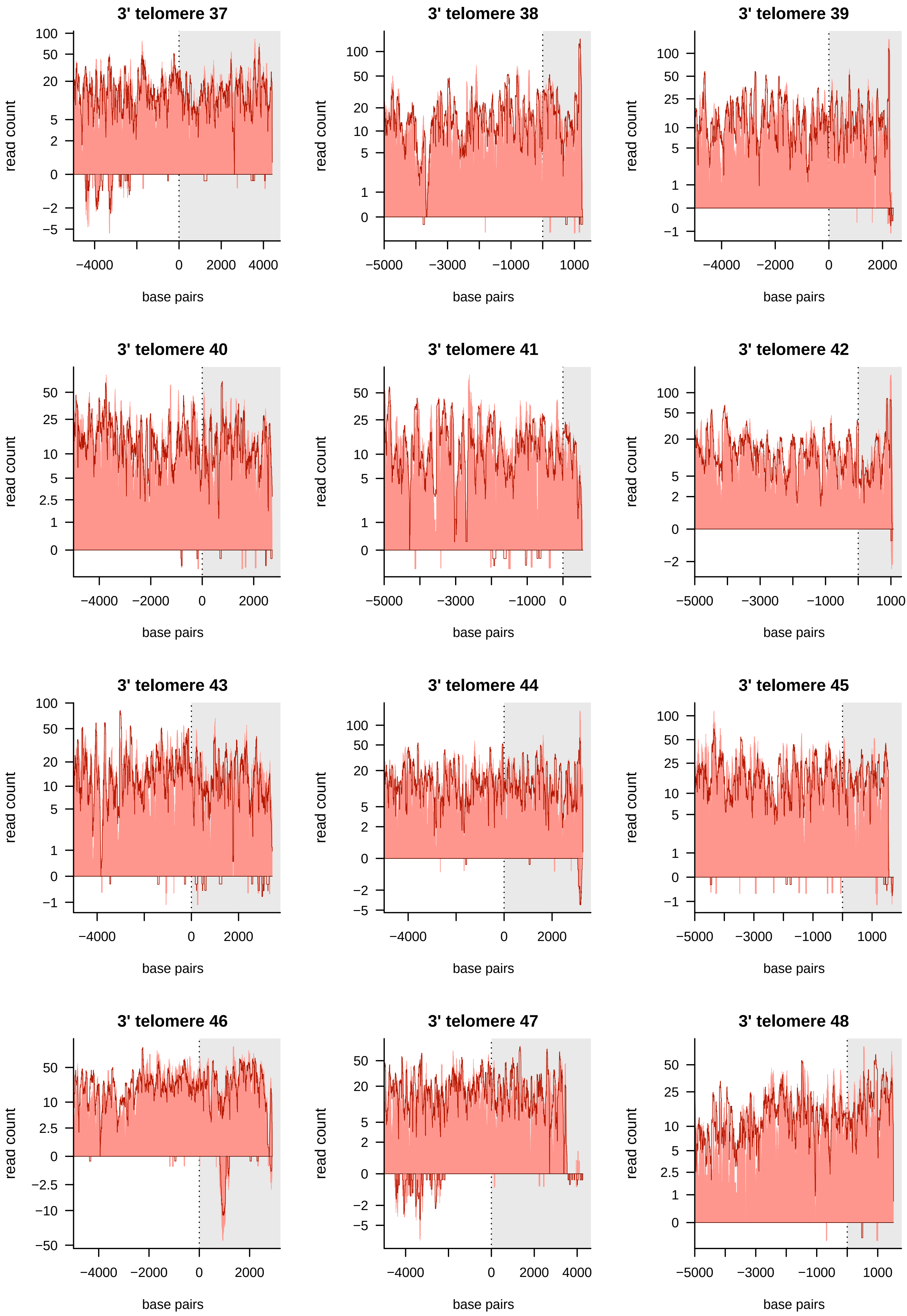

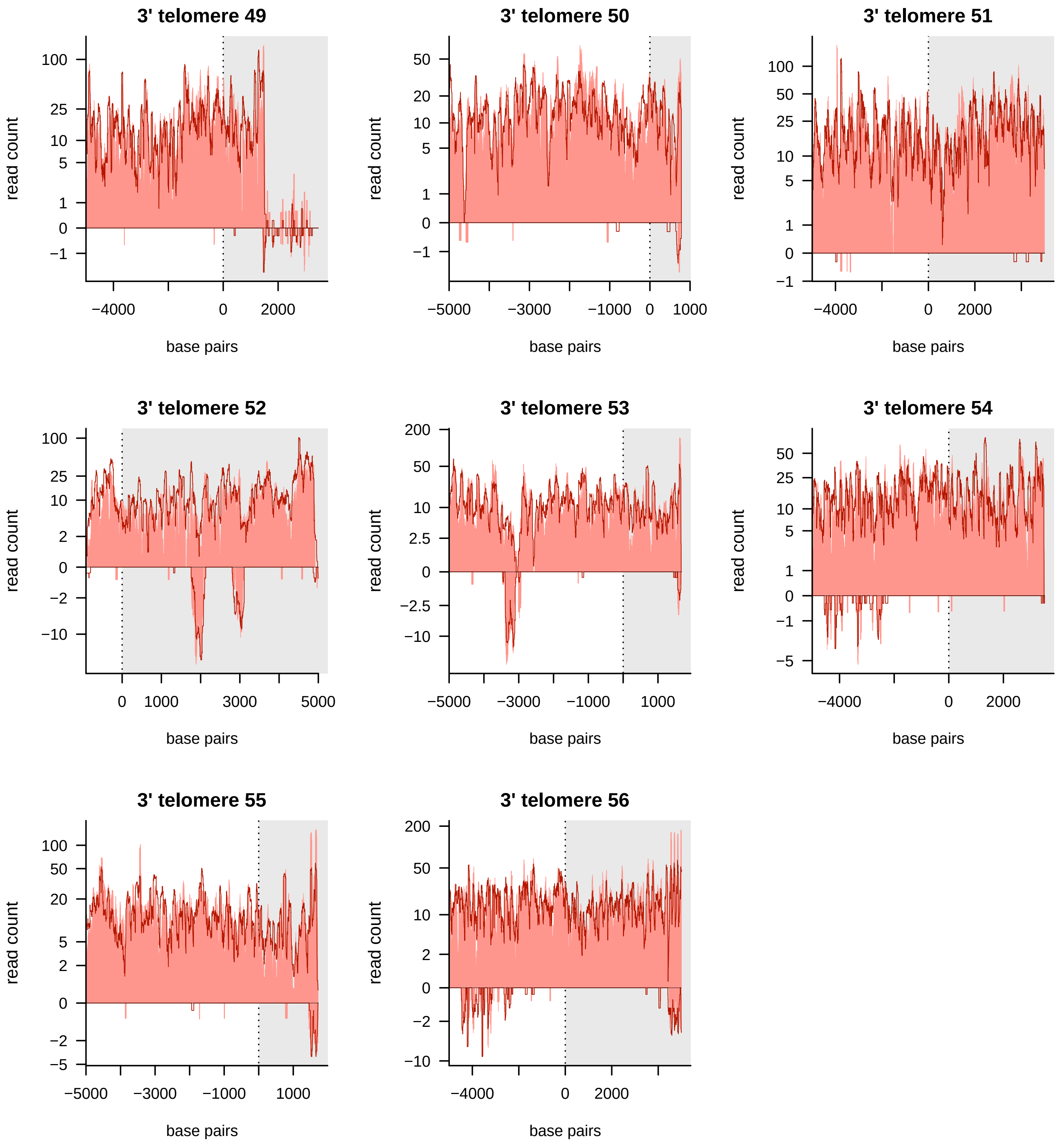
