## Supplementary material for "Genome-wide Quantification of Polycistronic Transcription in *Leishmania major*": Figure S2

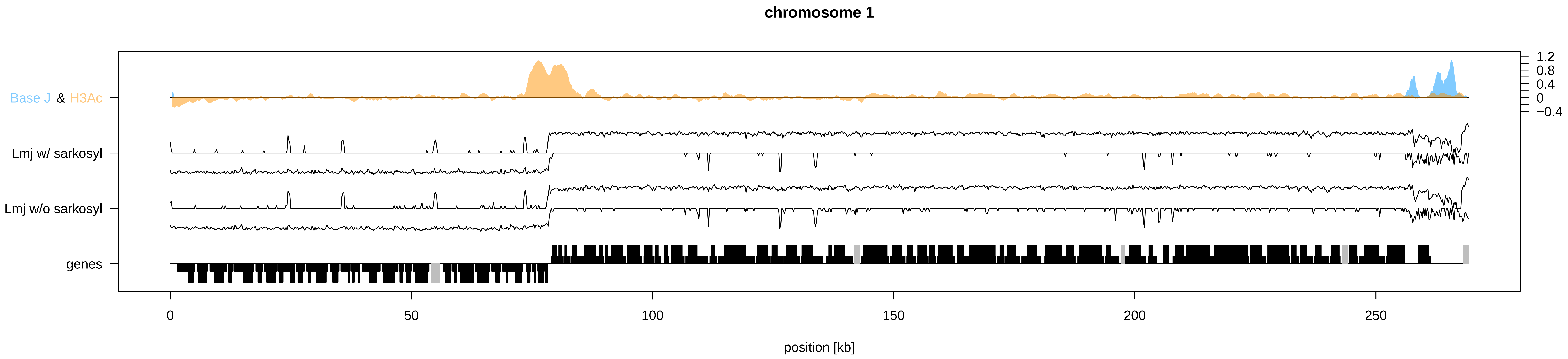

chromosome 1
Base J & H3Ac
Lmj w/ sarkosyl
Lmj w/o sarkosyl
genes
1.2
0.8
0.4
0
−0.4
0
50
100
150
200
250
position [kb]

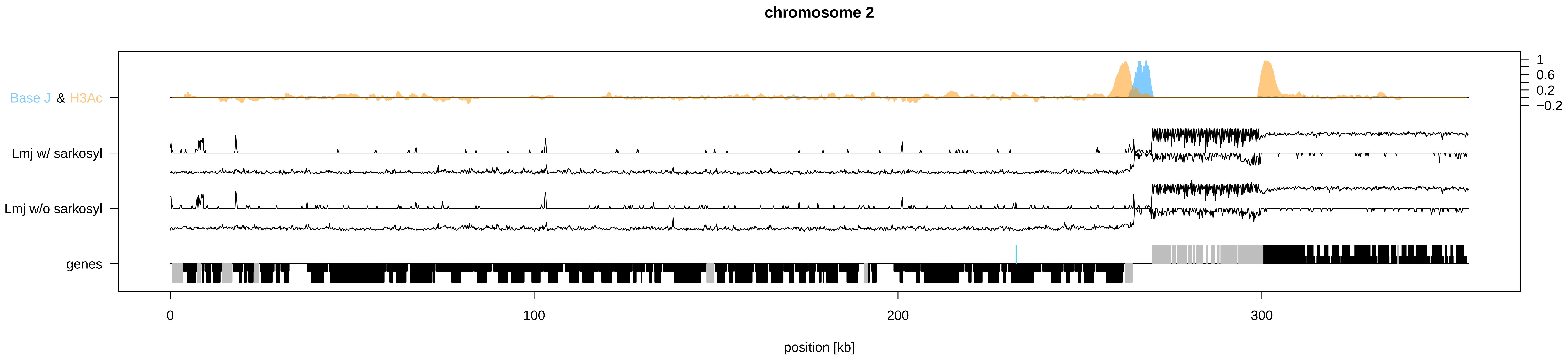
chromosome 2
Base J & H3Ac
Lmj w/ sarkosyl
Lmj w/o sarkosyl
genes
1
0.6
0.2
−0.2
0
100
200
300
position [kb]

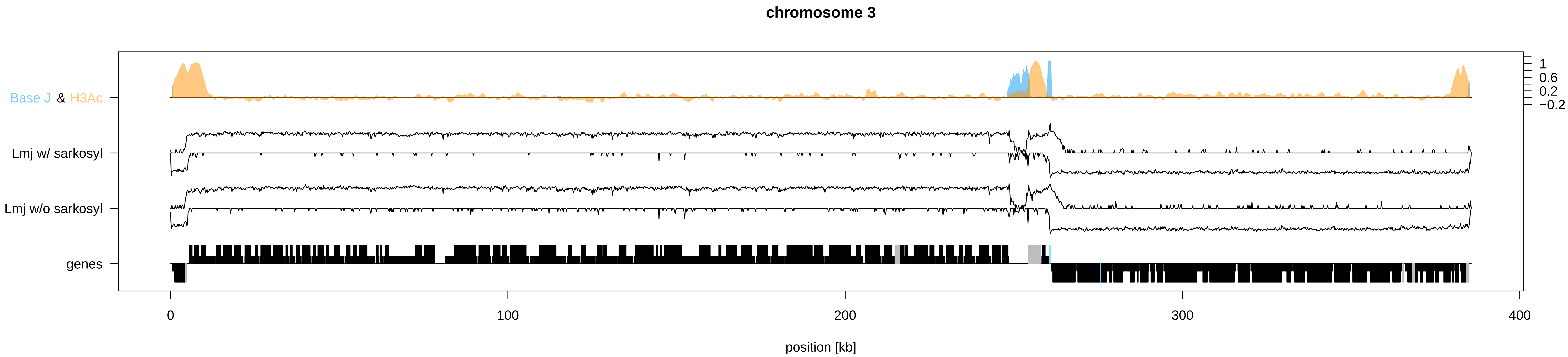
chromosome 3
Base J & H3Ac
Lmj w/ sarkosyl
Lmj w/o sarkosyl
genes
1
0.6
0.2
−0.2
0
100
200
300
400
position [kb]

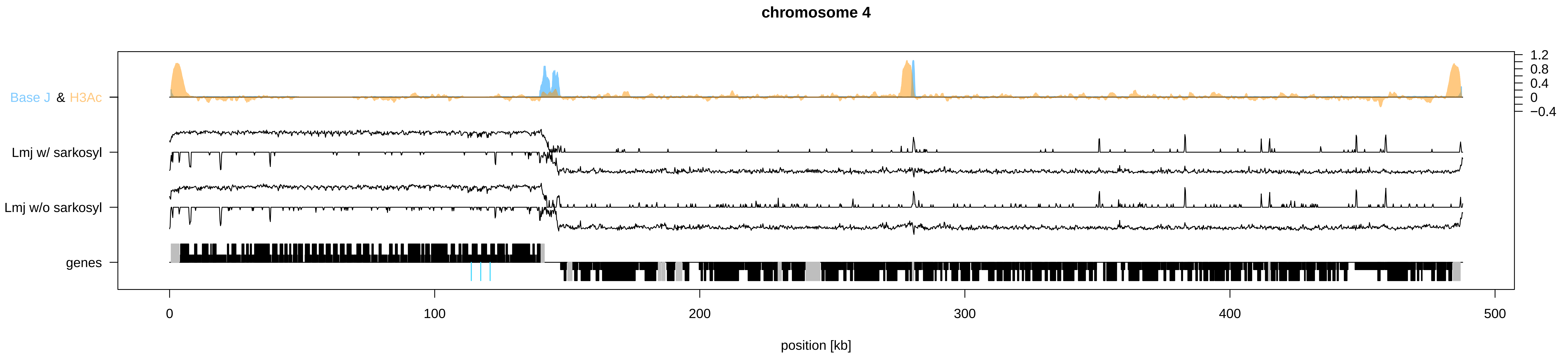

chromosome 4
Base J & H3Ac
Lmj w/ sarkosyl
Lmj w/o sarkosyl
genes
1.2
0.8
0.4
0
−0.4
0
100
200
300
400
500
position [kb]

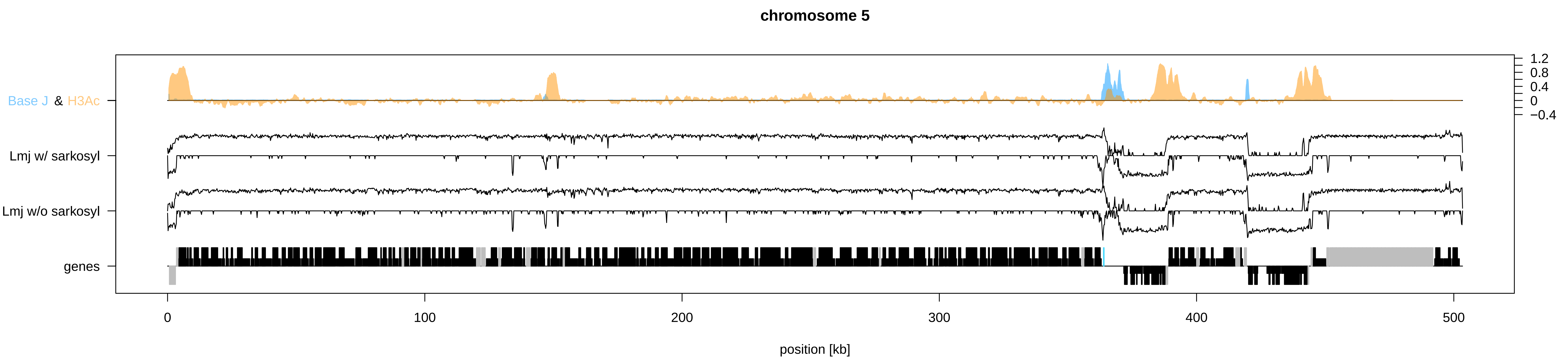

chromosome 5
Base J & H3Ac
Lmj w/ sarkosyl
Lmj w/o sarkosyl
genes
1.2
0.8
0.4
0
−0.4
0
100
200
300
400
500
position [kb]

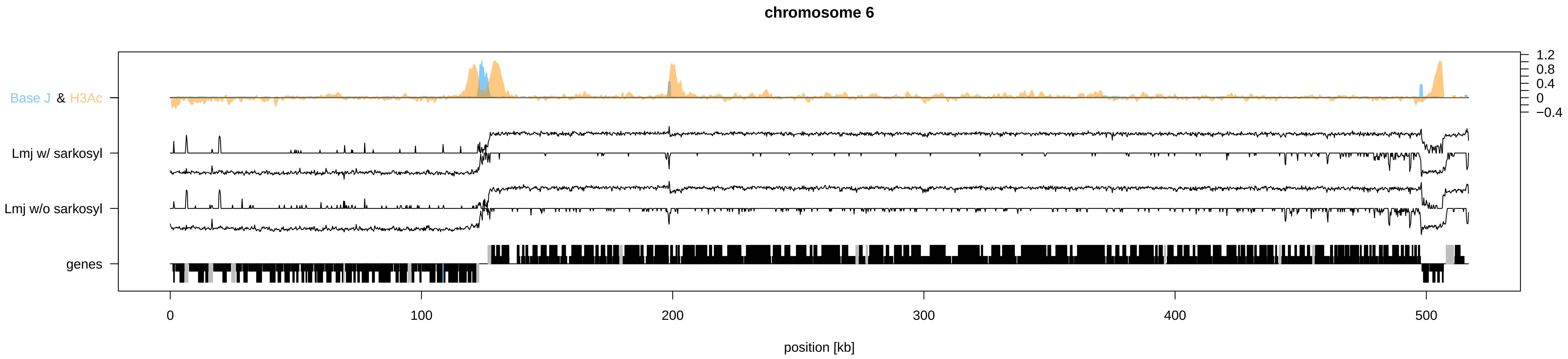

chromosome 6
Base J & H3Ac
Lmj w/ sarkosyl
Lmj w/o sarkosyl
genes
1.2
0.8
0.4
0
−0.4
0
100
200
300
400
500
position [kb]

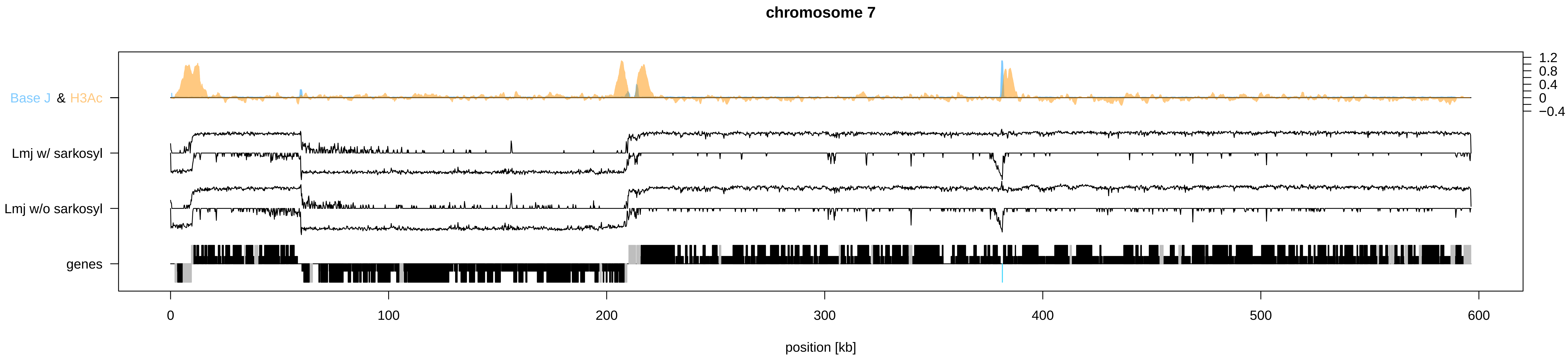
chromosome 7
Base J & H3Ac
Lmj w/ sarkosyl
Lmj w/o sarkosyl
genes
1.2
0.8
0.4
0
−0.4
0
100
200
300
400
500
600
position [kb]

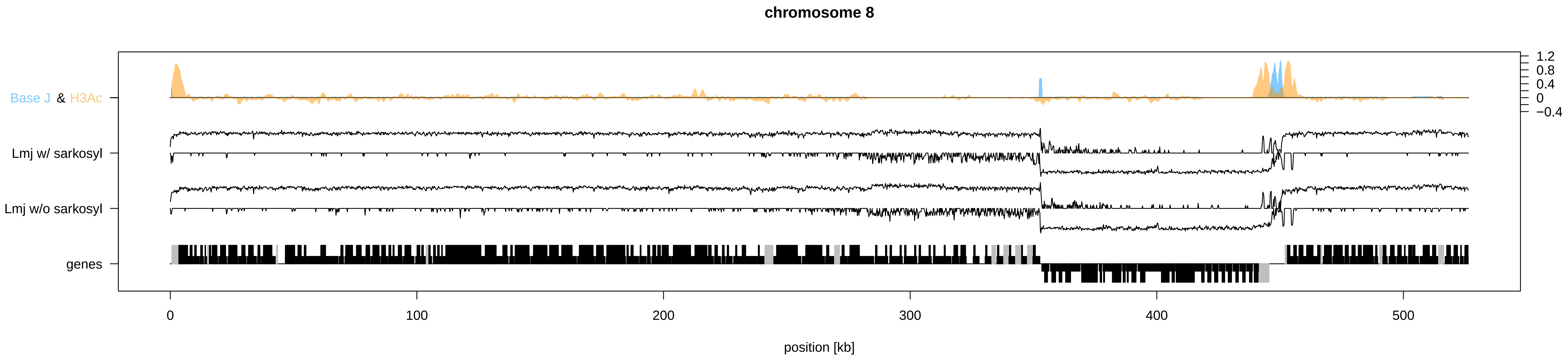

chromosome 8
Base J & H3Ac
Lmj w/ sarkosyl
Lmj w/o sarkosyl
genes
1.2
0.8
0.4
0
−0.4
0
100
200
300
400
500
position [kb]

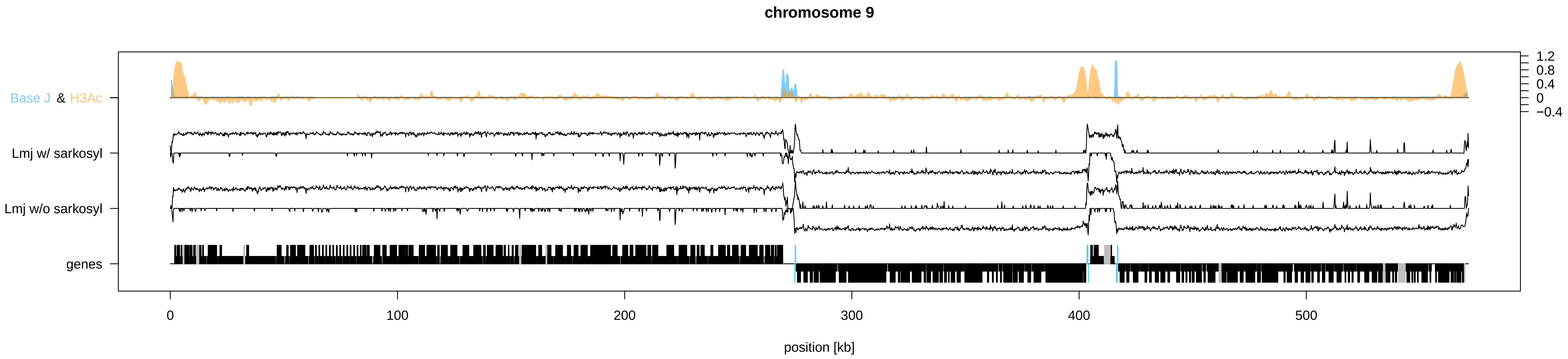
chromosome 9
Base J & H3Ac
Lmj w/ sarkosyl
Lmj w/o sarkosyl
genes
1.2
0.8
0.4
0
−0.4
0
100
200
300
400
500
position [kb]

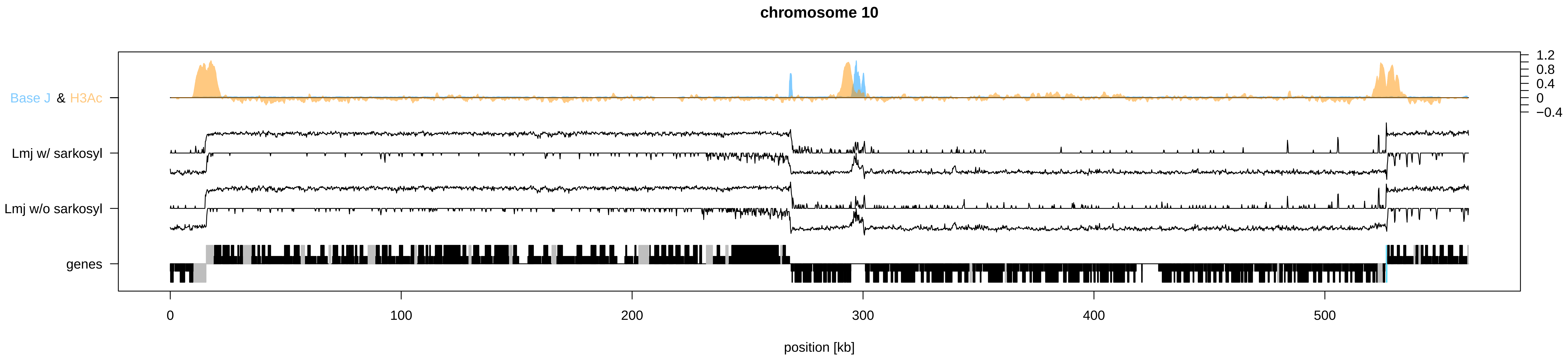
chromosome 10
Base J & H3Ac
Lmj w/ sarkosyl
Lmj w/o sarkosyl
genes
1.2
0.8
0.4
0
−0.4
0
100
200
300
400
500
position [kb]

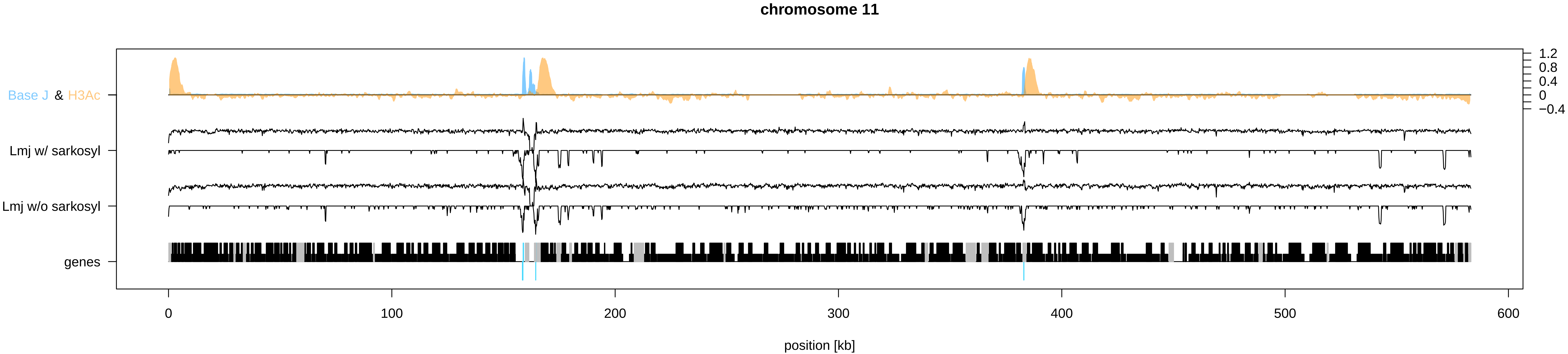
chromosome 11
Base J & H3Ac
Lmj w/ sarkosyl
Lmj w/o sarkosyl
genes
1.2
0.8
0.4
0
−0.4
0
100
200
300
400
500
600
position [kb]

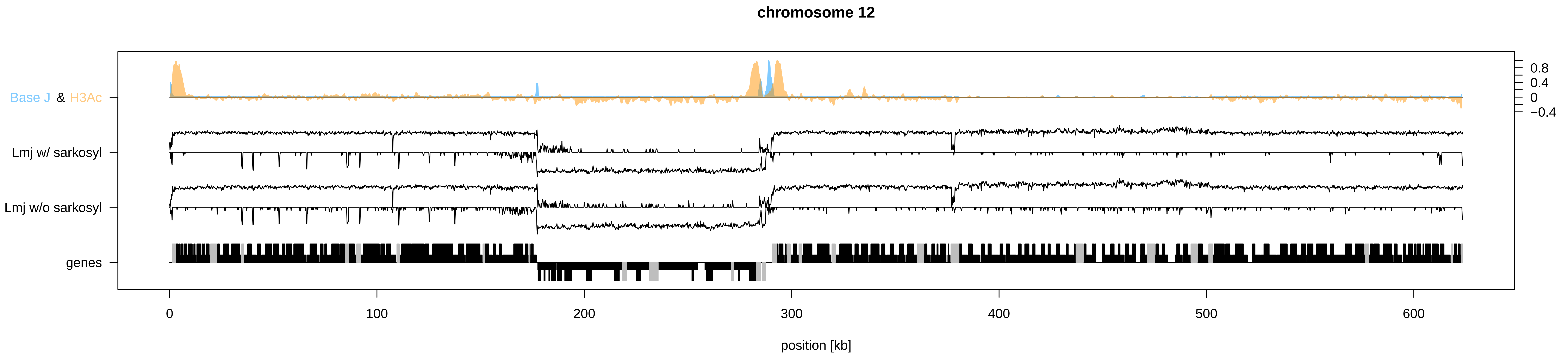
chromosome 12
Base J & H3Ac
Lmj w/ sarkosyl
Lmj w/o sarkosyl
genes
0.8
0.4
0
−0.4
0
100
200
300
400
500
600
position [kb]

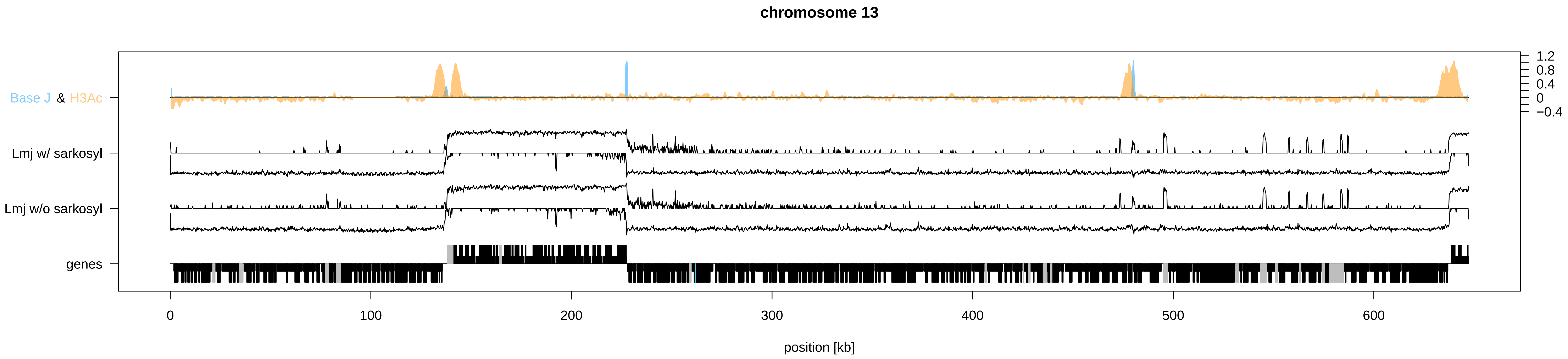

chromosome 13
Base J & H3Ac
Lmj w/ sarkosyl
Lmj w/o sarkosyl
genes
1.2
0.8
0.4
0
−0.4
0
100
200
300
400
500
600
position [kb]

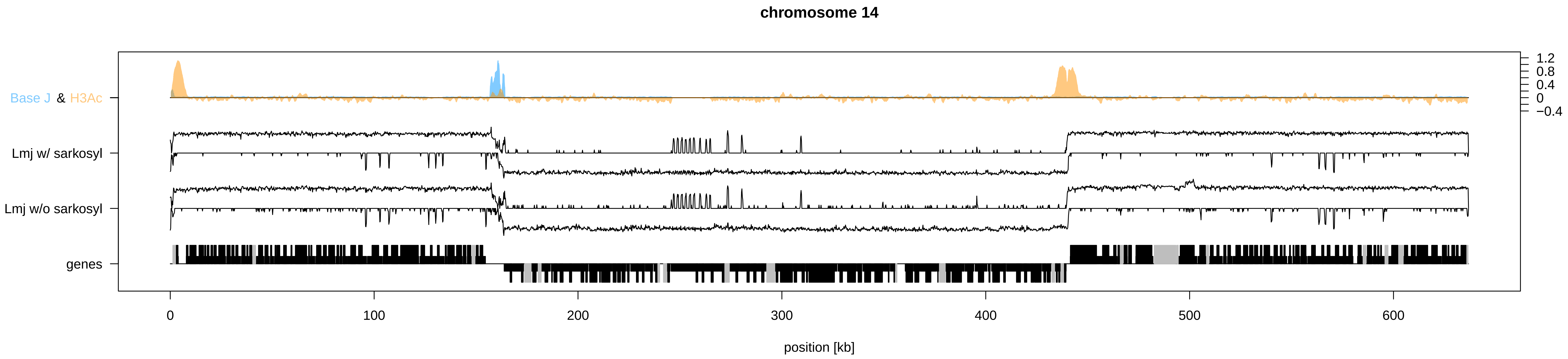

chromosome 14
Base J & H3Ac
Lmj w/ sarkosyl
Lmj w/o sarkosyl
genes
1.2
0.8
0.4
0
−0.4
0
100
200
300
400
500
600
position [kb]

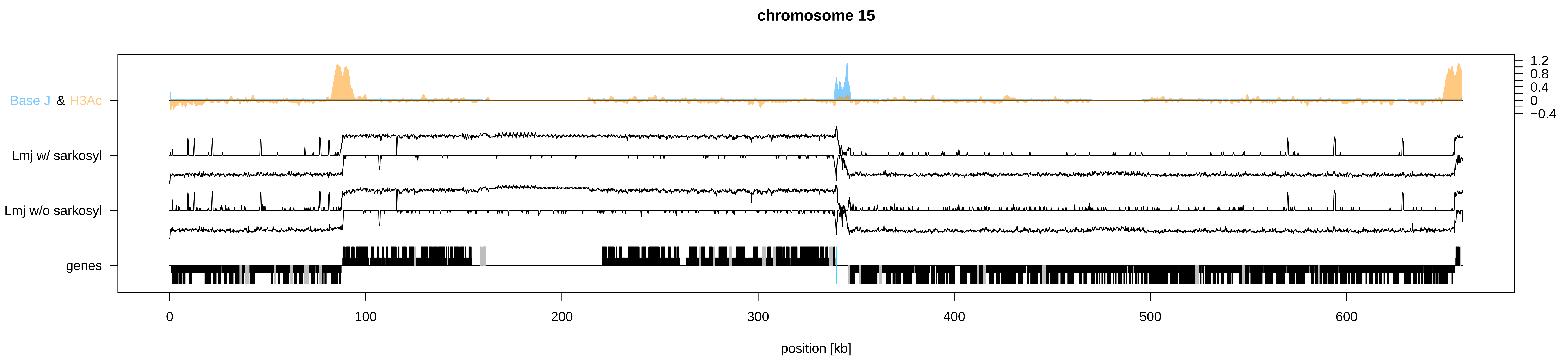

chromosome 15
Base J & H3Ac
Lmj w/ sarkosyl
Lmj w/o sarkosyl
genes
1.2
0.8
0.4
0
−0.4
0
100
200
300
400
500
600
position [kb]

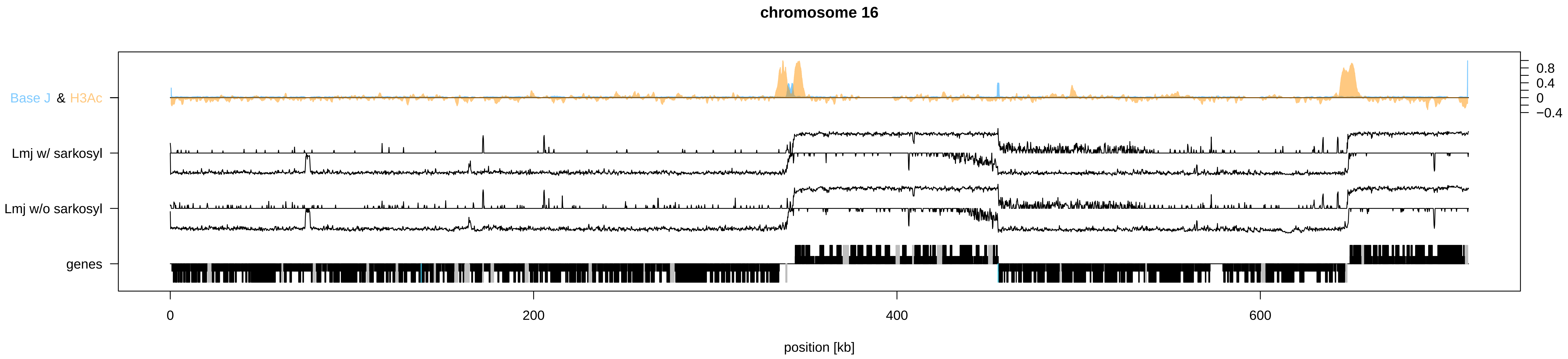

chromosome 16
Base J & H3Ac
Lmj w/ sarkosyl
Lmj w/o sarkosyl
genes
0.8
0.4
0
−0.4
0
200
400
600
position [kb]

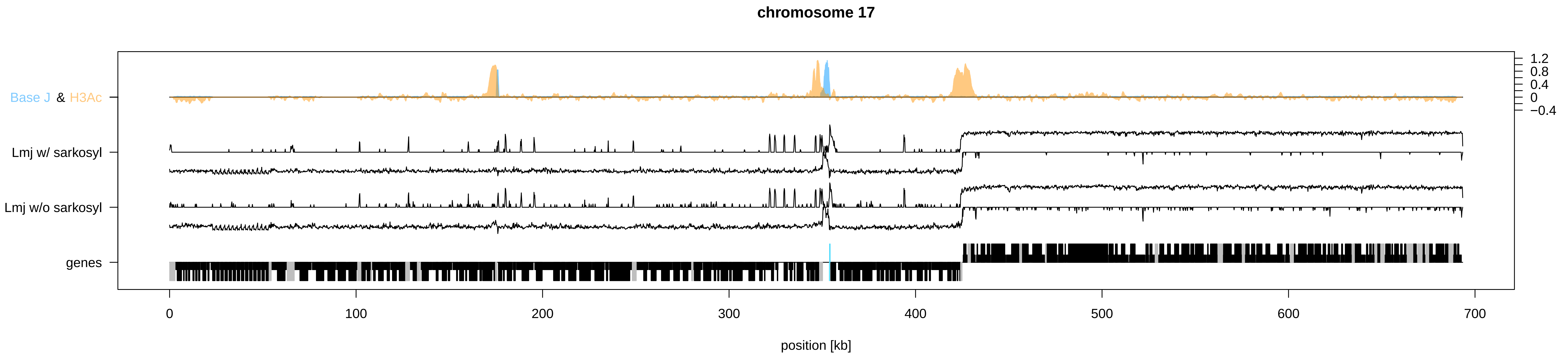

chromosome 17
Base J & H3Ac
Lmj w/ sarkosyl
Lmj w/o sarkosyl
genes
1.2
0.8
0.4
0
−0.4
0
100
200
300
400
500
600
700
position [kb]

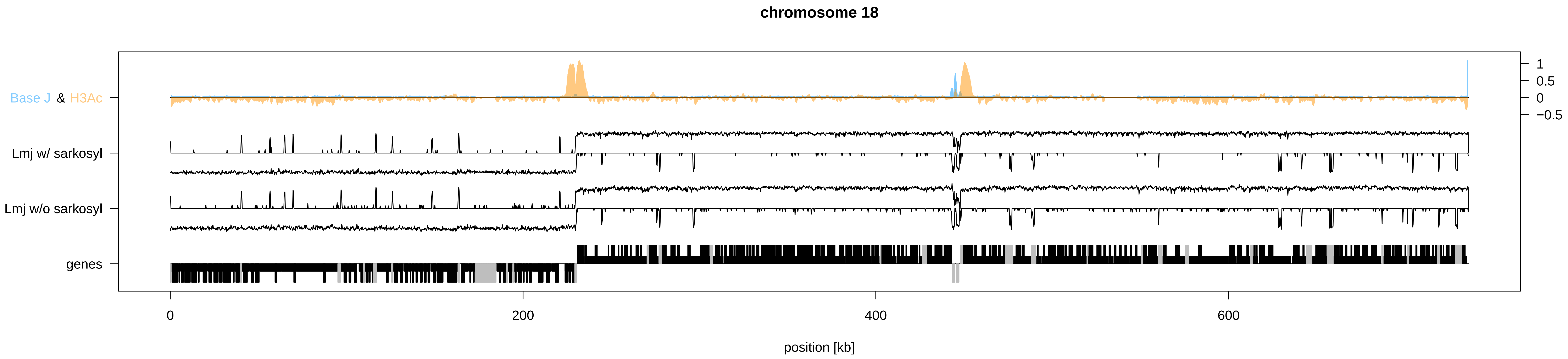
chromosome 18
Base J & H3Ac
Lmj w/ sarkosyl
Lmj w/o sarkosyl
genes
1
0.5
0
−0.5
0
200
400
600
position [kb]

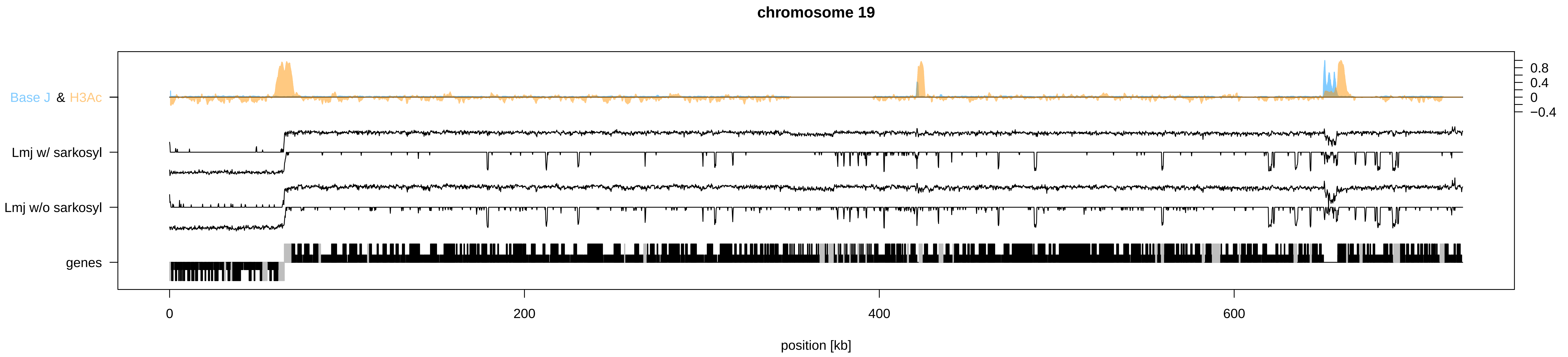
chromosome 19
Base J & H3Ac
Lmj w/ sarkosyl
Lmj w/o sarkosyl
genes
0.8
0.4
0
−0.4
0
200
400
600
position [kb]

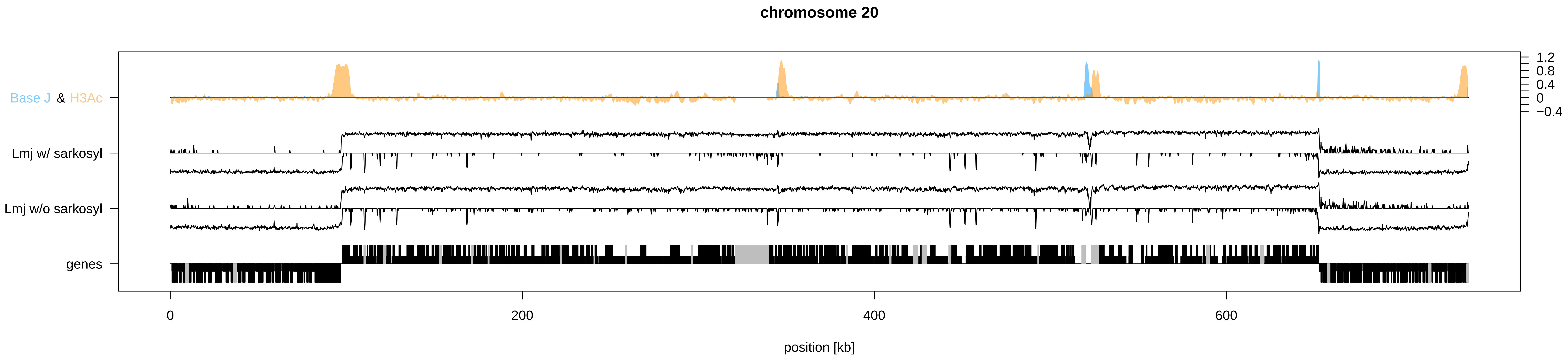

chromosome 20
Base J & H3Ac
Lmj w/ sarkosyl
Lmj w/o sarkosyl
genes
1.2
0.8
0.4
0
−0.4
0
200
400
600
position [kb]

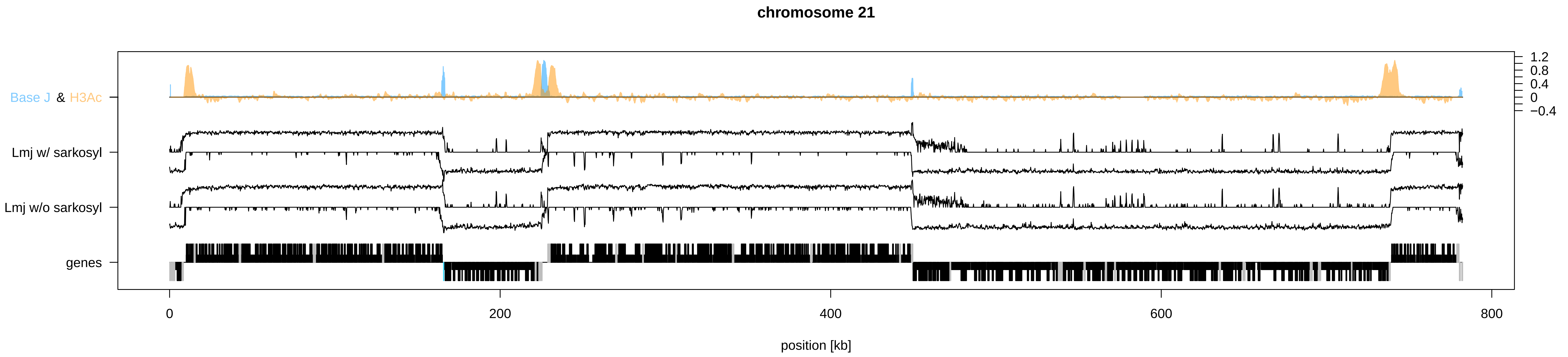

chromosome 21
Base J & H3Ac
Lmj w/ sarkosyl
Lmj w/o sarkosyl
genes
1.2
0.8
0.4
0
−0.4
0
200
400
600
800
position [kb]

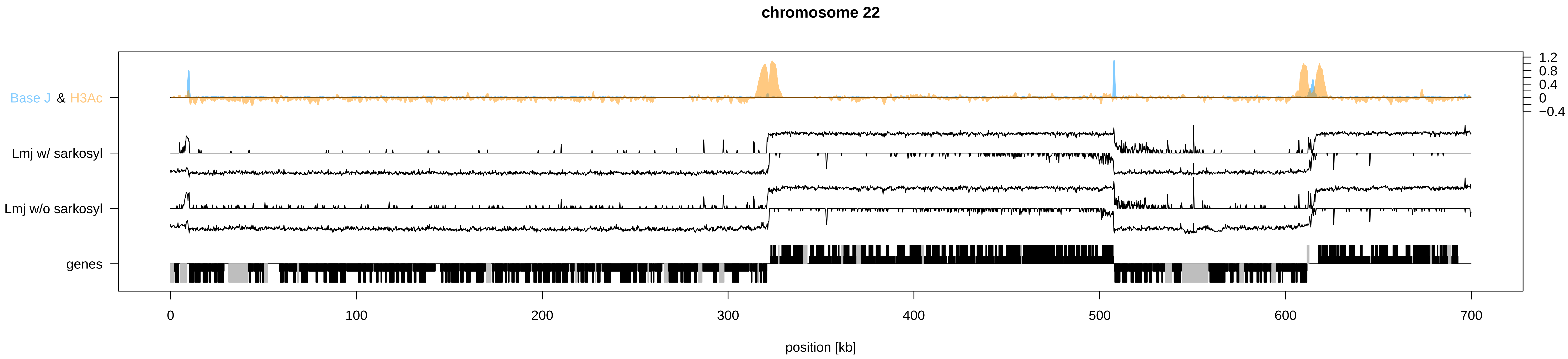

chromosome 22
Base J & H3Ac
Lmj w/ sarkosyl
Lmj w/o sarkosyl
genes
1.2
0.8
0.4
0
−0.4
0
100
200
300
400
500
600
700
position [kb]

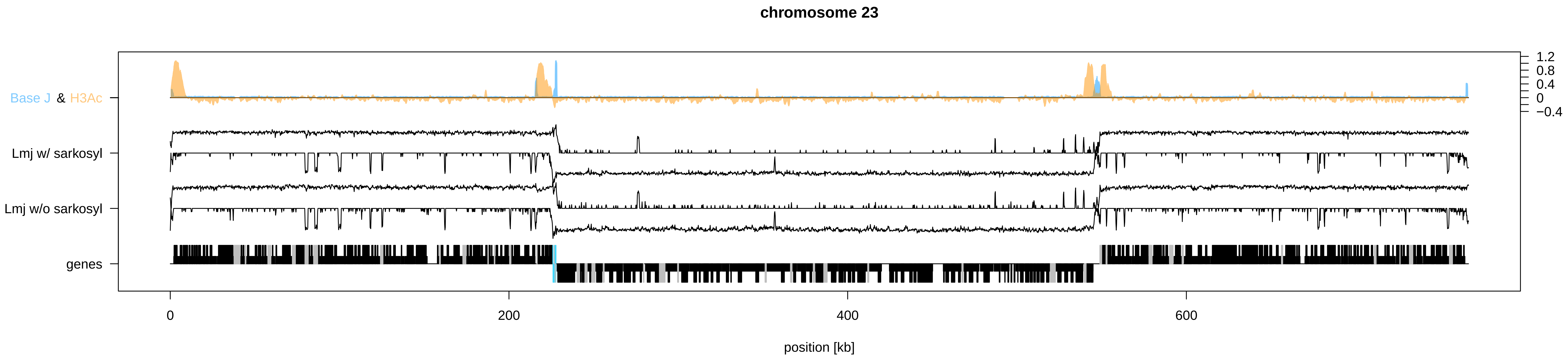
chromosome 23
Base J & H3Ac
Lmj w/ sarkosyl
Lmj w/o sarkosyl
genes
1.2
0.8
0.4
0
−0.4
0
200
400
600
position [kb]

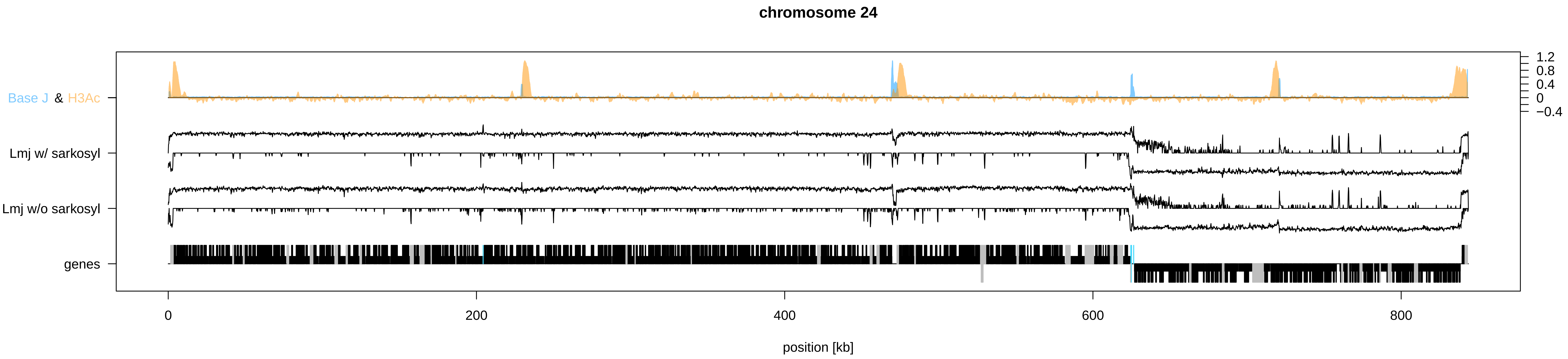
chromosome 24
Base J & H3Ac
Lmj w/ sarkosyl
Lmj w/o sarkosyl
genes
1.2
0.8
0.4
0
−0.4
0
200
400
600
800
position [kb]

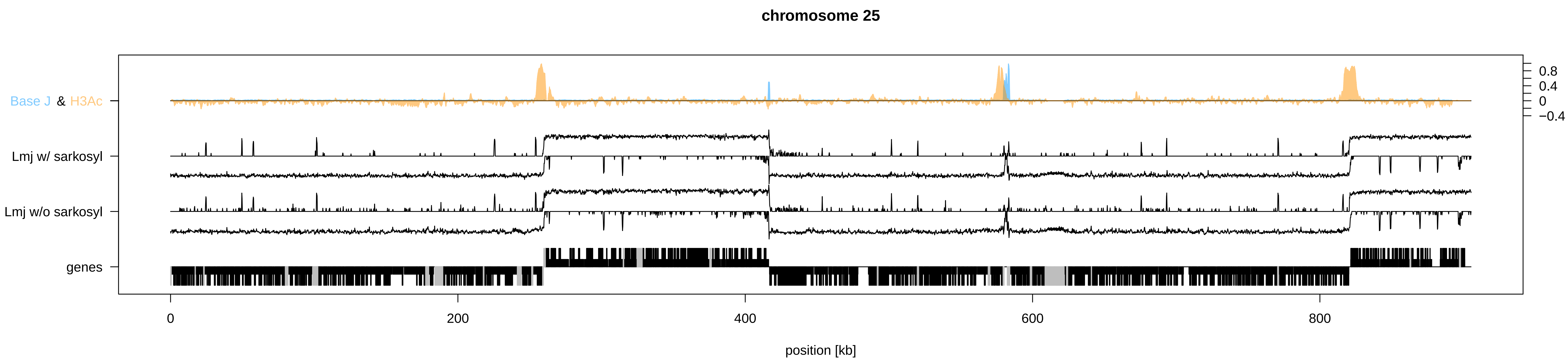
chromosome 25
Base J & H3Ac
Lmj w/ sarkosyl
Lmj w/o sarkosyl
genes
0.8
0.4
0
−0.4
0
200
400
600
800
position [kb]

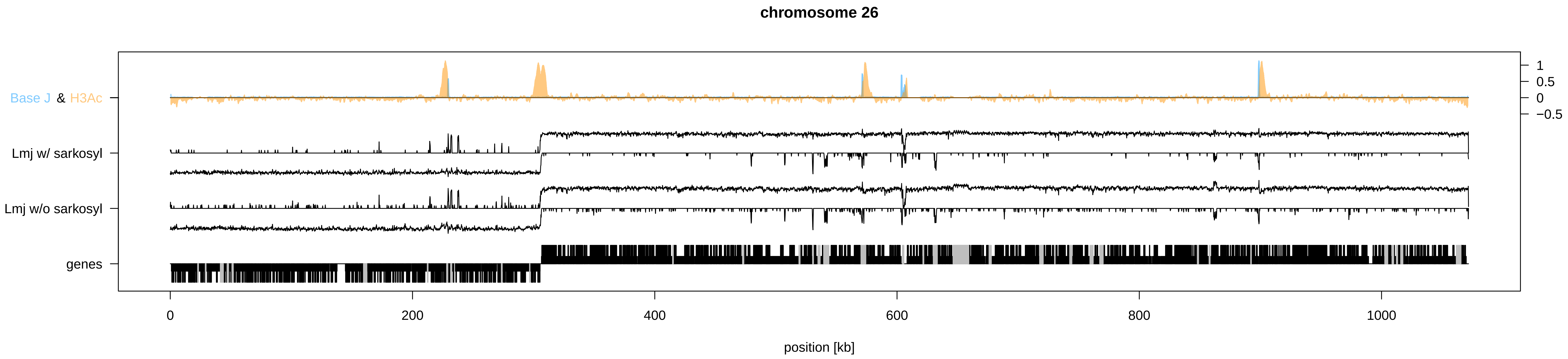
chromosome 26
Base J & H3Ac
Lmj w/ sarkosyl
Lmj w/o sarkosyl
genes
1
0.5
0
−0.5
0
200
400
600
800
1000
position [kb]

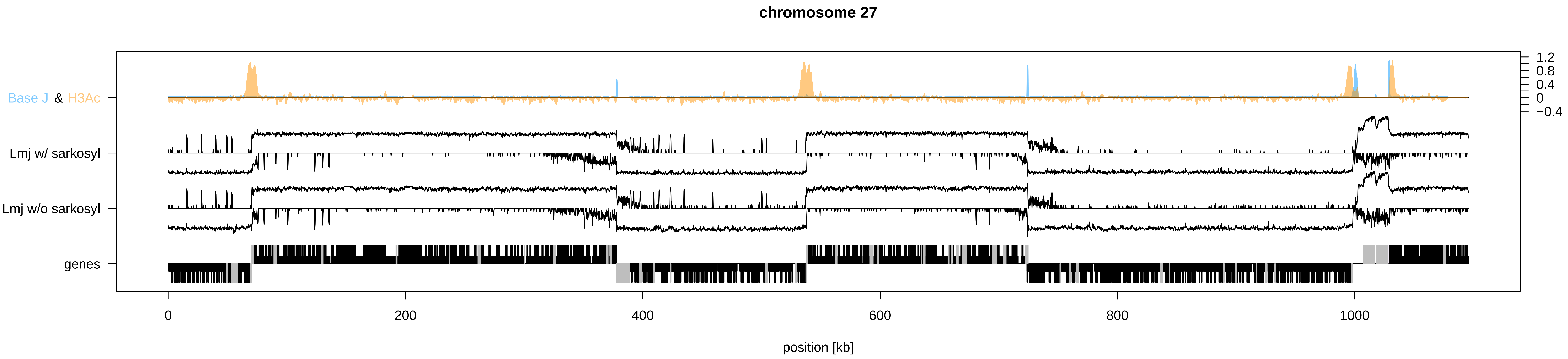

chromosome 27
Base J & H3Ac
Lmj w/ sarkosyl
Lmj w/o sarkosyl
genes
1.2
0.8
0.4
0
−0.4
0
200
400
600
800
1000
position [kb]

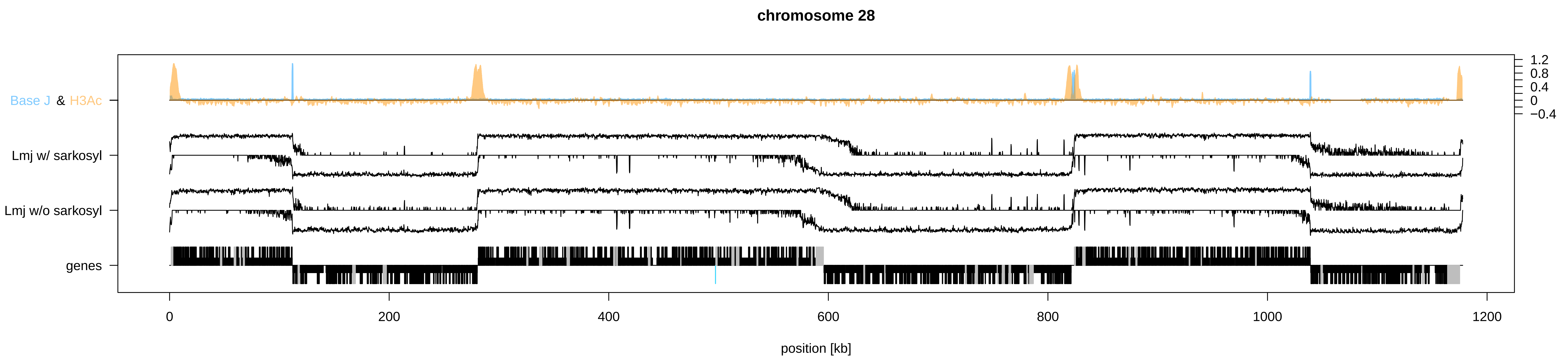

chromosome 28
Base J & H3Ac
Lmj w/ sarkosyl
Lmj w/o sarkosyl
genes
1.2
0.8
0.4
0
−0.4
0
200
400
600
800
1000
1200
position [kb]

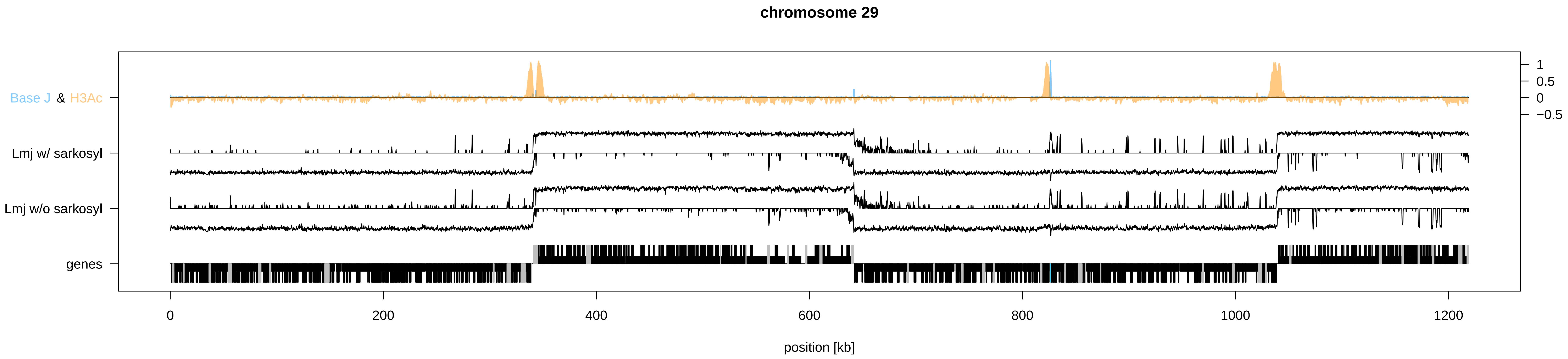
chromosome 29
Base J & H3Ac
Lmj w/ sarkosyl
Lmj w/o sarkosyl
genes
1
0.5
0
−0.5
0
200
400
600
800
1000
1200
position [kb]

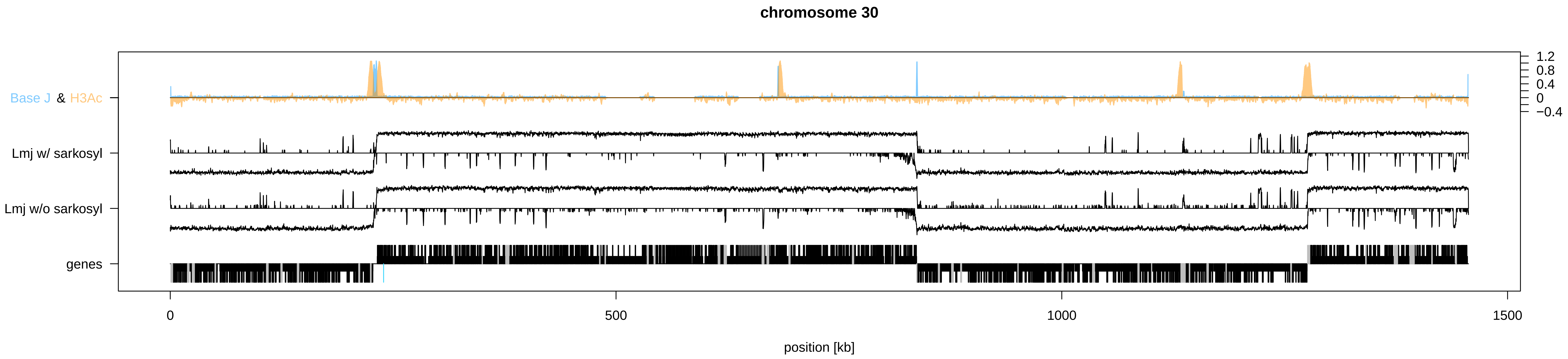
chromosome 30
Base J & H3Ac
Lmj w/ sarkosyl
Lmj w/o sarkosyl
genes
1.2
0.8
0.4
0
−0.4
0
500
1000
1500
position [kb]

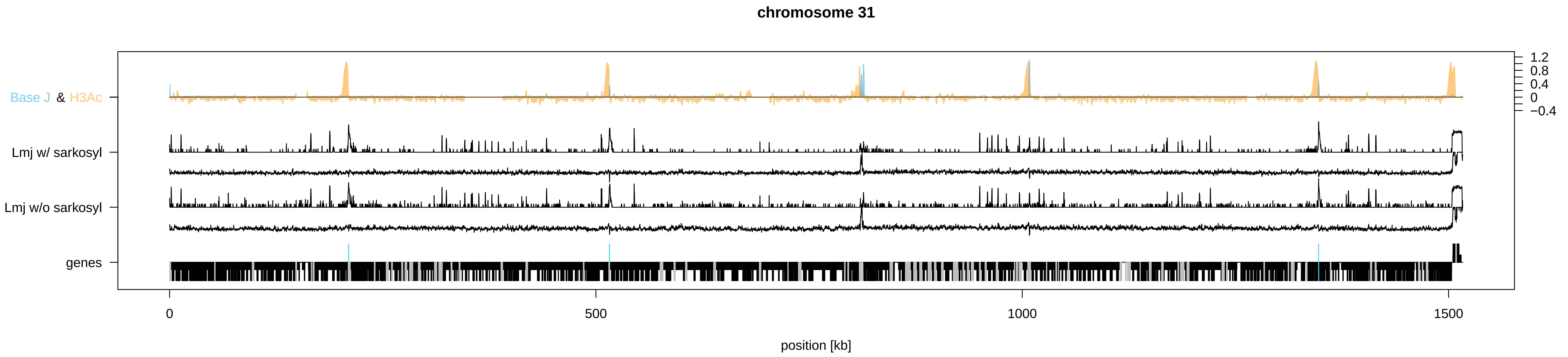
chromosome 31
Base J & H3Ac
Lmj w/ sarkosyl
Lmj w/o sarkosyl
genes
1.2
0.8
0.4
0
−0.4
0
500
1000
1500
position [kb]

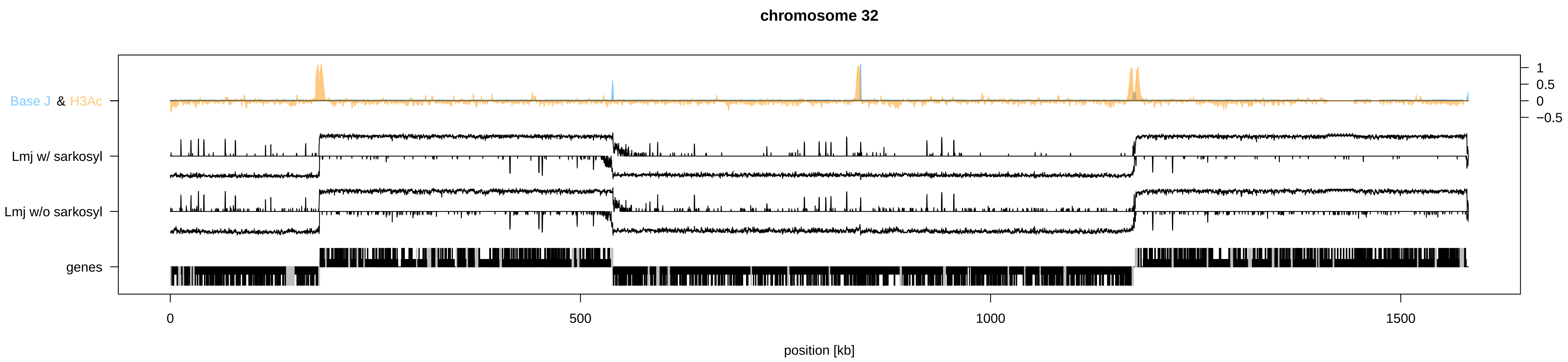

chromosome 32
Base J & H3Ac
Lmj w/ sarkosyl
Lmj w/o sarkosyl
genes
1
0.5
0
−0.5
0
500
1000
1500
position [kb]

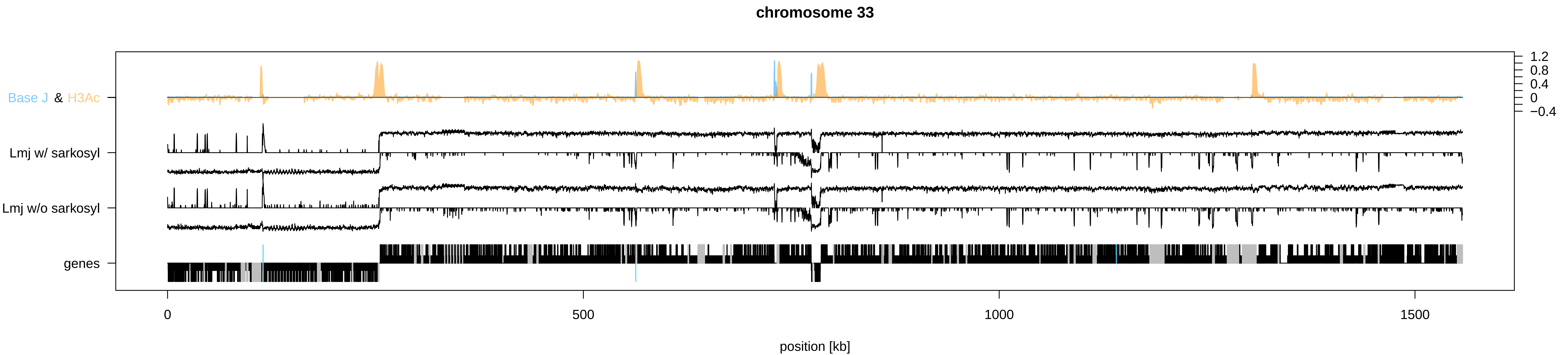

chromosome 33
Base J & H3Ac
Lmj w/ sarkosyl
Lmj w/o sarkosyl
genes
1.2
0.8
0.4
0
−0.4
0
500
1000
1500
position [kb]

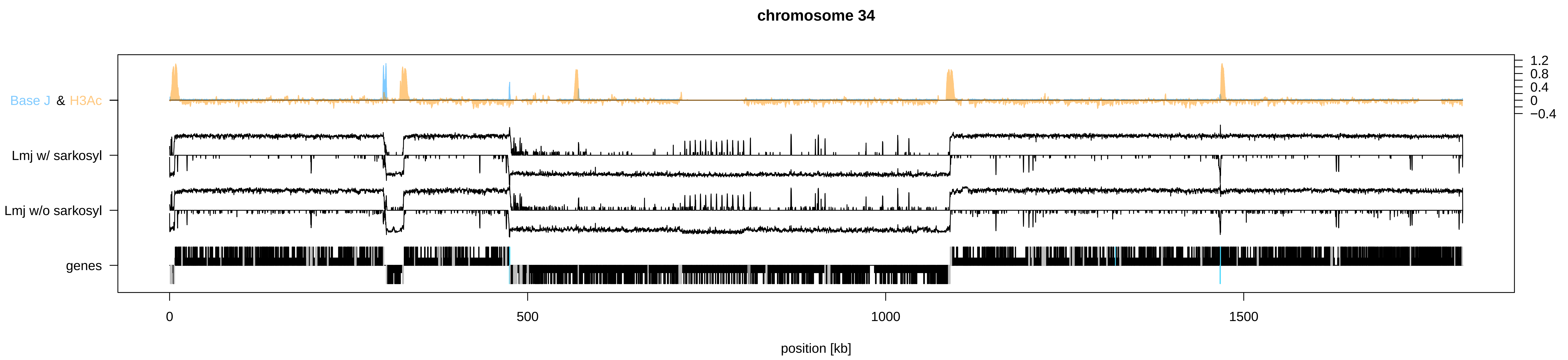
chromosome 34
Base J & H3Ac
Lmj w/ sarkosyl
Lmj w/o sarkosyl
genes
1.2
0.8
0.4
0
−0.4
0
500
1000
1500
position [kb]

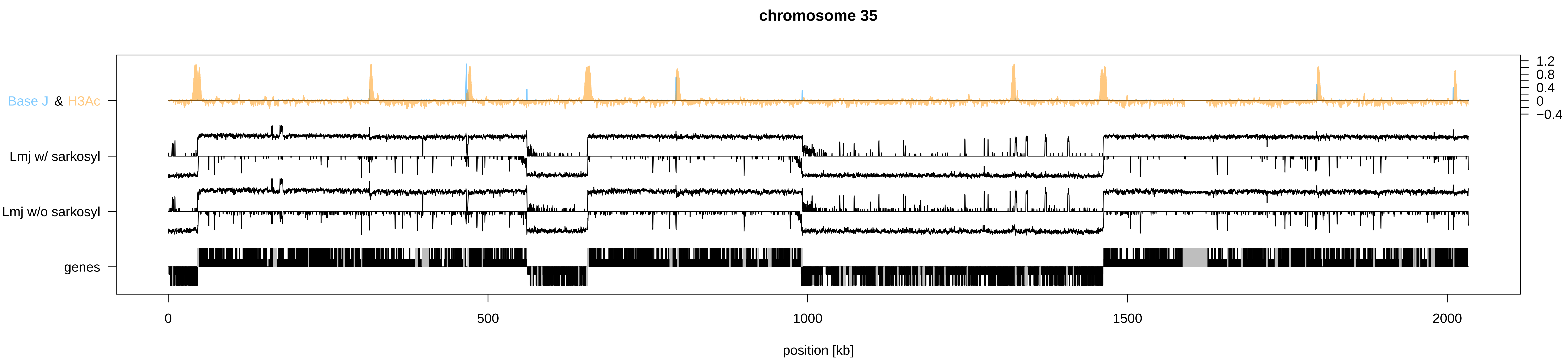

chromosome 35
Base J & H3Ac
Lmj w/ sarkosyl
Lmj w/o sarkosyl
genes
1.2
0.8
0.4
0
−0.4
0
500
1000
1500
2000
position [kb]

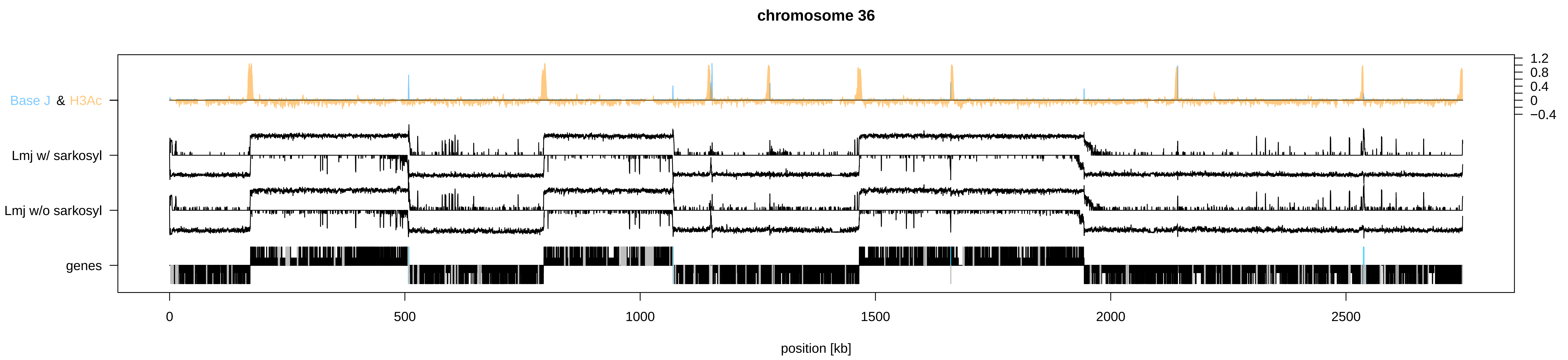
chromosome 36
Base J & H3Ac
Lmj w/ sarkosyl
Lmj w/o sarkosyl
genes
1.2
0.8
0.4
0
−0.4
0
500
1000
1500
2000
2500
position [kb]
