## Supplementary material for "Genome-wide Quantification of Polycistronic Transcription in *Leishmania major*": Table S2

**Table S2:** List of Transcription start and stop sites by chromosomes and position. We defined dSSRs as area between the first protein-coding sequences of diverging PTUs, cSSRs as area between the last protein-coding sequences of converging PTUs, 5' telomers as area between the ends of chromosomes and the first protein-coding sequences of a PTU, and 3' telomers as area between the last protein-coding sequence of a PTU and the ends of chromosomes. See also Figure S2.

| Type | Number | Chromosome | from | to |
|---|---|---|---|---|
| dSSR | 1 | 1 | 78,230 | 79,200 |
| dSSR | 2 | 2 | 261,616 | 300,448 |
| dSSR | 3 | 3 | 4,186 | 5,558 |
| dSSR | 4 | 5 | 387,825 | 389,276 |
| dSSR | 5 | 5 | 443,004 | 445,376 |
| dSSR | 6 | 6 | 121,741 | 127,954 |
| dSSR | 7 | 6 | 506,750 | 511,537 |
| dSSR | 8 | 7 | 5,410 | 10,624 |
| dSSR | 9 | 7 | 208,072 | 215,794 |
| dSSR | 10 | 8 | 441,116 | 452,902 |
| dSSR | 11 | 9 | 402,953 | 405,115 |
| dSSR | 12 | 10 | 9,830 | 19,134 |
| dSSR | 13 | 10 | 525,964 | 527,151 |
| dSSR | 14 | 12 | 282,463 | 293,016 |
| dSSR | 15 | 13 | 135,544 | 141,219 |
| dSSR | 16 | 13 | 636,831 | 638,676 |
| dSSR | 17 | 14 | 439,347 | 441,531 |
| dSSR | 18 | 15 | 87,351 | 88,403 |
| dSSR | 19 | 15 | 654,086 | 654,086 |
| dSSR | 20 | 16 | 335,004 | 344,007 |
| dSSR | 21 | 16 | 646,455 | 649,376 |
| dSSR | 22 | 17 | 424,126 | 425,761 |
| dSSR | 23 | 18 | 229,036 | 230,955 |
| dSSR | 24 | 19 | 61,070 | 68,592 |
| dSSR | 25 | 20 | 96,656 | 97,973 |
| dSSR | 26 | 21 | 6,975 | 10,085 |
| dSSR | 27 | 21 | 223,229 | 230,692 |

| Type | Number | Chromosome | from | to |
|---|---|---|---|---|
| dSSR | 28 | 21 | 737,784 | 739,384 |
| dSSR | 29 | 22 | 320,883 | 323,027 |
| dSSR | 30 | 22 | 611,516 | 617,689 |
| dSSR | 31 | 23 | 544,889 | 550,249 |
| dSSR | 32 | 24 | 838,294 | 839,531 |
| dSSR | 33 | 25 | 258,467 | 261,018 |
| dSSR | 34 | 25 | 820,134 | 821,495 |
| dSSR | 35 | 26 | 305,250 | 306,722 |
| dSSR | 36 | 27 | 68,762 | 72,396 |
| dSSR | 37 | 27 | 537,128 | 538,942 |
| dSSR | 38 | 27 | 996,907 | 1,029,451 |
| dSSR | 39 | 28 | 280,191 | 281,280 |
| dSSR | 40 | 28 | 821,132 | 825,498 |
| dSSR | 41 | 29 | 338,788 | 344,918 |
| dSSR | 42 | 29 | 1,038,709 | 1,040,259 |
| dSSR | 43 | 30 | 227,263 | 232,302 |
| dSSR | 44 | 30 | 1,275,101 | 1,278,983 |
| dSSR | 45 | 31 | 1,503,554 | 1,505,231 |
| dSSR | 46 | 32 | 180,672 | 182,576 |
| dSSR | 47 | 32 | 1,172,102 | 1,179,166 |
| dSSR | 48 | 33 | 252,678 | 255,682 |
| dSSR | 49 | 33 | 784,811 | 785,972 |
| dSSR | 50 | 34 | 5,990 | 7,890 |
| dSSR | 51 | 34 | 322,331 | 327,654 |
| dSSR | 52 | 34 | 1,086,566 | 1,092,822 |
| dSSR | 53 | 35 | 45,582 | 47,975 |
| dSSR | 54 | 35 | 653,805 | 656,897 |
| dSSR | 55 | 35 | 1,461,540 | 1,463,046 |
| dSSR | 56 | 36 | 170,954 | 172,041 |
| dSSR | 57 | 36 | 794,327 | 795,587 |
| dSSR | 58 | 36 | 1,464,913 | 1,465,734 |

| Type | Number | Chromosome | from | to |
|---|---|---|---|---|
| cSSR | 1 | 3 | 259,274 | 261,544 |
| cSSR | 2 | 4 | 139,787 | 148,810 |
| cSSR | 3 | 5 | 362,575 | 371,957 |
| cSSR | 4 | 5 | 417,458 | 420,207 |
| cSSR | 5 | 6 | 497,366 | 498,866 |
| cSSR | 6 | 7 | 56,723 | 61,078 |
| cSSR | 7 | 8 | 350,776 | 354,497 |
| cSSR | 8 | 9 | 269,559 | 276,029 |
| cSSR | 9 | 9 | 414,328 | 414,328 |
| cSSR | 10 | 10 | 266,426 | 269,224 |
| cSSR | 11 | 12 | 175,349 | 177,700 |
| cSSR | 12 | 13 | 227,327 | 228,609 |
| cSSR | 13 | 14 | 153,315 | 166,682 |
| cSSR | 14 | 15 | 339,001 | 347,163 |
| cSSR | 15 | 16 | 455,218 | 456,162 |
| cSSR | 16 | 20 | 652,251 | 653,747 |
| cSSR | 17 | 21 | 164,873 | 166,694 |
| cSSR | 18 | 21 | 448,151 | 449,814 |
| cSSR | 19 | 22 | 507,185 | 508,121 |
| cSSR | 20 | 23 | 225,574 | 228,809 |
| cSSR | 21 | 24 | 622,523 | 627,001 |
| cSSR | 22 | 25 | 414,507 | 417,033 |
| cSSR | 23 | 27 | 377,820 | 390,143 |
| cSSR | 24 | 27 | 720,501 | 723,738 |
| cSSR | 25 | 28 | 111,627 | 112,246 |
| cSSR | 26 | 28 | 587,105 | 596,188 |
| cSSR | 27 | 28 | 1,038,898 | 1,039,552 |
| cSSR | 28 | 29 | 637,891 | 642,271 |
| cSSR | 29 | 30 | 836,948 | 837,884 |
| cSSR | 30 | 32 | 535,701 | 539,885 |

| Type | Number | Chromosome | from | to |
|---|---|---|---|---|
| cSSR | 31 | 33 | 773,888 | 774,327 |
| cSSR | 32 | 34 | 298,096 | 304,186 |
| cSSR | 33 | 34 | 474,060 | 476,124 |
| cSSR | 34 | 35 | 559,643 | 565,916 |
| cSSR | 35 | 35 | 989,052 | 989,864 |
| cSSR | 36 | 36 | 505,456 | 511,743 |
| cSSR | 37 | 36 | 1,068,937 | 1,072,704 |
| cSSR | 38 | 36 | 1,942,310 | 1,943,748 |
| 5`telomers | 1 | 3 | 385,501 | 383,878 |
| 5`telomers | 2 | 4 | 1 | 4,146 |
| 5`telomers | 3 | 4 | 487,861 | 483,664 |
| 5`telomers | 4 | 5 | 1 | 4,442 |
| 5`telomers | 5 | 8 | 1 | 3,439 |
| 5`telomers | 6 | 9 | 1 | 1,949 |
| 5`telomers | 7 | 9 | 571,341 | 569,253 |
| 5`telomers | 8 | 11 | 1 | 1,605 |
| 5`telomers | 9 | 12 | 1 | 3,085 |
| 5`telomers | 10 | 14 | 1 | 2,866 |
| 5`telomers | 11 | 20 | 737,675 | 736,321 |
| 5`telomers | 12 | 23 | 1 | 2,230 |
| 5`telomers | 13 | 24 | 1 | 3,571 |
| 5`telomers | 14 | 28 | 1 | 3,155 |
| 5`telomers | 15 | 28 | 1,177,749 | 1,163,470 |
| 5`telomers | 16 | 36 | 2,748,001 | 2,746,207 |
| 3' telomers | 1 | 1 | 3,766 | 1 |
| 3' telomers | 2 | 1 | 260,876 | 269,237 |
| 3' telomers | 3 | 2 | 4,580 | 1 |
| 3' telomers | 4 | 2 | 355,491 | 356,844 |
| 3' telomers | 5 | 3 | 1,251 | 1 |

| Type | Number | Chromosome | from | to |
|---|---|---|---|---|
| 3' telomers | 6 | 5 | 501,425 | 503,331 |
| 3' telomers | 7 | 6 | 1,339 | 1 |
| 3' telomers | 8 | 6 | 513,467 | 516,716 |
| 3' telomers | 9 | 7 | 3,231 | 1 |
| 3' telomers | 10 | 7 | 591,870 | 596,309 |
| 3' telomers | 11 | 8 | 526,224 | 526,473 |
| 3' telomers | 12 | 10 | 120 | 1 |
| 3' telomers | 13 | 10 | 559,580 | 562,130 |
| 3' telomers | 14 | 11 | 581,800 | 583,214 |
| 3' telomers | 15 | 12 | 622,392 | 623,629 |
| 3' telomers | 16 | 13 | 1,956 | 1 |
| 3' telomers | 17 | 13 | 646,990 | 647,297 |
| 3' telomers | 18 | 14 | 635,637 | 636,703 |
| 3' telomers | 19 | 15 | 1,208 | 1 |
| 3' telomers | 20 | 15 | 657,393 | 659,292 |
| 3' telomers | 21 | 16 | 1,647 | 1 |
| 3' telomers | 22 | 16 | 712,247 | 714,615 |
| 3' telomers | 23 | 17 | 4,071 | 1 |
| 3' telomers | 24 | 17 | 691,349 | 693,320 |
| 3' telomers | 25 | 18 | 1,144 | 1 |
| 3' telomers | 26 | 18 | 734,097 | 735,931 |
| 3' telomers | 27 | 19 | 1,080 | 1 |
| 3' telomers | 28 | 19 | 727,426 | 728,899 |
| 3' telomers | 29 | 20 | 968 | 1 |
| 3' telomers | 30 | 21 | 4,529 | 1 |
| 3' telomers | 31 | 21 | 777,256 | 782,422 |
| 3' telomers | 32 | 22 | 2,388 | 1 |
| 3' telomers | 33 | 22 | 692,474 | 699,868 |
| 3' telomers | 34 | 23 | 764,115 | 766,686 |
| 3' telomers | 35 | 24 | 840,948 | 843,515 |
| 3' telomers | 36 | 25 | 2,096 | 1 |

| Type | Number | Chromosome | from | to |
|---|---|---|---|---|
| 3' telomers | 37 | 25 | 900,841 | 905,260 |
| 3' telomers | 38 | 26 | 1,267 | 1 |
| 3' telomers | 39 | 26 | 1,069,358 | 1,071,765 |
| 3' telomers | 40 | 27 | 2,722 | 1 |
| 3' telomers | 41 | 27 | 1,095,308 | 1,095,864 |
| 3' telomers | 42 | 29 | 1,079 | 1 |
| 3' telomers | 43 | 29 | 1,215,435 | 1,218,866 |
| 3' telomers | 44 | 30 | 3,285 | 1 |
| 3' telomers | 45 | 30 | 1,454,449 | 1,456,173 |
| 3' telomers | 46 | 31 | 2,902 | 1 |
| 3' telomers | 47 | 31 | 1,512,268 | 1,516,535 |
| 3' telomers | 48 | 32 | 1,517 | 1 |
| 3' telomers | 49 | 32 | 1,579,124 | 1,582,589 |
| 3' telomers | 50 | 33 | 786 | 1 |
| 3' telomers | 51 | 33 | 1,548,847 | 1,557,258 |
| 3' telomers | 52 | 34 | 5,068 | 1 |
| 3' telomers | 53 | 34 | 1,804,028 | 1,805,702 |
| 3' telomers | 54 | 35 | 3,507 | 1 |
| 3' telomers | 55 | 35 | 2,031,317 | 2,033,042 |
| 3' telomers | 56 | 36 | 9,590 | 1 |
