## Supplementary material for "Genome-wide Quantification of Polycistronic Transcription in *Leishmania major*": Table S1

Table S1: Sequencing output

| Sample | raw reads | trimmed reads | 5’ dATP | 5’ dCTP | 5’ dGTP | 5’ dTTP | Aligned to Genome |
|---|---|---|---|---|---|---|---|
| Lmj w/ sarkosyl 1 | 2.22E+07 | 1.51E+07 | 1.31E+06 | 8.99E+05 | 1.23E+07 | 6.05E+05 | 92.75% |
| Lmj w/ sarkosyl 2 | 2.14E+07 | 1.47E+07 | 1.29E+06 | 8.69E+05 | 1.20E+07 | 5.62E+05 | 93.64% |
| Lmj w/ sarkosyl 3 | 2.53E+07 | 1.58E+07 | 1.56E+06 | 1.09E+06 | 1.24E+07 | 7.02E+05 | 92.95% |
| **mean** | **2.30E+07** | **1.52E+07** | **1.39E+06** | **9.52E+05** | **1.22E+07** | **6.23E+05** | **93.1%** |
| Lmj w/o sarkosyl 1 | 2.45E+07 | 1.03E+07 | 2.39E+06 | 2.07E+06 | 5.35E+06 | 4.50E+05 | 84.74% |
| Lmj w/o sarkosyl 2 | 2.60E+07 | 9.59E+06 | 2.18E+06 | 1.89E+06 | 5.11E+06 | 4.16E+05 | 85.04% |
| Lmj w/o sarkosyl 3 | 2.34E+07 | 9.89E+06 | 2.25E+06 | 1.97E+06 | 5.25E+06 | 4.10E+05 | 85.39% |
| **mean** | **2.46E+07** | **9.91E+06** | **2.28E+06** | **1.98E+06** | **5.24E+06** | **4.25E+05** | **85.1%** |
